# A Modular Master Regulator Landscape Determines the Impact of Genetic Alterations on the Transcriptional Identity of Cancer Cells

**DOI:** 10.1101/758268

**Authors:** Evan O. Paull, Alvaro Aytes, Prem Subramaniam, Federico M. Giorgi, Eugene F. Douglass, Brennan Chu, Sunny J. Jones, Siyuan Zheng, Roel Verhaak, Cory Abate-Shen, Mariano J. Alvarez, Andrea Califano

## Abstract

Despite considerable pan-cancer efforts, the link between genomics and transcriptomics in cancer remains relatively weak and mostly based on statistical rather than mechanistic principles. By performing integrative analysis of transcriptomic and mutational profiles on a sample-by-sample basis, via regulatory/signaling networks, we identified a repertoire of 407 Master-Regulator proteins responsible for canalizing the genetics of 20 TCGA cohorts into 112 transcriptionally-distinct tumor subtypes. Further analysis highlighted a highly-recurrent regulatory architecture (*oncotecture*) with Master-Regulators organized into 24 modular *MR-Blocks,* regulating highly-specific tumor-hallmark functions and predictive of patient outcome. Critically, >50% of the somatic alterations identified in individual samples were in proteins affecting Master-Regulator activity, thus yielding novel insight into mechanisms linking tumor genetics and transcriptional identity and establishing novel non-oncogene dependencies. Experimental validation of functional mutations upstream of the most conserved MR-Block confirmed their ability to affect MR-protein activity, suggesting that the proposed methodology may effectively complement and extend current pan-cancer knowledge.

## Introduction

Our understanding of cancer as a complex system is constantly evolving: in particular, it is increasingly appreciated that the transcriptional state of cancer cells (i.e. their transcriptional identity) is as tightly regulated as that of their physiologic counterparts, albeit via distinct and aberrant (i.e., *dystatic*) regulatory mechanisms (Califano and Alvarez, 2017). These mechanisms play a key role in determining which transcriptional identities may be compatible with the specific set of somatic and germline variants harbored by cancer cells, as well as their likelihood to plastically reprogram across molecularly-distinct identities. Some mutations effectively restrict the transcriptional identity repertoire accessible to a cancer cell; for instance, activating mutations in *ESR1*, *FOXA1*, and *GATA3* are observed almost exclusively in the luminal subtype of breast cancer (Curtis et al., 2012b). However, most mutations are not as restrictive. In glioblastoma, for instance, there is only weak association between mutational and transcriptional states (Neftel et al., 2019). Indeed, EGFR mutations, while more frequently associated with a proneural identity, are also detected in mesenchymal cells.

While it is reasonable to expect that a tumor cell’s mutational landscape may mechanistically constrain the subset of transcriptional identities occupied by its cells and affect their relative likelihood (E.g., EGFR mutations in GBM may increase the likelihood of a proneural state), the specific regulatory and signaling logic that underlies these relationships is still elusive, with most mutation/transcriptional-subtype relationships based on statistical associations that lack mechanistic rationale. Indeed, the vast majority of studies aimed at elucidating the molecular landscape of large tumor cohorts proceed almost invariably in two steps, first by identifying molecularly-distinct subtypes by gene expression cluster analysis and then by assessing subtype-specific enrichment in recurrent mutations (Hoadley et al., 2018).

To address this challenge, we propose to leverage the *Oncotecture* hypothesis (Califano and Alvarez, 2017). This proposes the existence of Master Regulator (MR) proteins organized in highly modular structures (Tumor Checkpoints) that integrate the effect of upstream signals and genomic alterations to implement specific transcriptional states. We hypothesize that this may pinpoint more specific relationships between a tumor cell’s mutational landscape and its transcriptional identities. Here we distinguish between transcriptional states (which may be transient and thus form a continuum) and identities (i.e., stable states, representing peaks in the probability density of the states thus associated with higher persistence over time). This is important because in TCGA bulk tissue, one is more likely to observe identities than states.

The *Oncotecture hypothesis*, which represents the cancer-specific counterpart of the *Omnigene Hypothesis* in human genetics (Boyle et al., 2017), is supported by a wealth of experimental evidence, from prostate cancer (Aytes et al., 2014b) and breast cancer (Rodriguez-Barrueco et al., 2015; Walsh et al., 2017), to glioblastoma (Carro et al., 2010), neuroblastoma (Rajbhandari et al., 2018a), and neuroendocrine tumors (Alvarez et al., 2018), see (Califano and Alvarez, 2017) for a comprehensive overview, but has not yet been comprehensively and systematically assessed across multiple tumor types.

In this manuscript we thus explore and validate the Oncotecture hypothesis across the entire TCGA repository (Cancer Genome Atlas Research et al., 2013), on a sample-by-sample basis. Specifically, we aim to assess the full range of MR-proteins representing candidate mechanistic determinants of cancer cell identity, their conservation across distinct tumor cohorts, their ability to canalize the effect of specific genetic alterations, and, finally, whether the transcriptional identities they regulate may recapitulate patient outcome and other macroscopic properties. While TCGA does not comprise metastatic samples, the same approach is equally effective in repositories that include samples from metastatic or heavily treated patients, as shown for the Metabric breast cancer repository (Curtis et al., 2012a).

To accomplish this goal, we developed the Multi Omics Master-Regulator Analysis (MOMA) framework. MOMA allows single-sample-based identification of candidate Master Regulators that are downstream of sample-specific, functional genetic alterations—as identified and validated by GISTIC2.0 (Mermel et al., 2011) and CHASM (Carter et al., 2009)—and (b) mechanistically determine a sample’s transcriptional identity, via their regulatory targets. For simplicity, we will use the term *Master Regulator* to indicate a candidate MR and *validated Master Regulator* to indicate one that has been experimentally validated.

MOMA analysis of 9,738 individual primary samples, representing 20 TCGA tumor cohorts of sufficient size to support the analysis, identified 112 transcriptionally distinct, MR-driven tumor identities (or *subtypes*), each one regulated by a distinct Tumor Checkpoint, whose aberrant activity is determined by distinct genomic alteration landscapes. Unexpectedly, the MRs found in the Tumor Checkpoints present a highly recurrent sub-modular structure, implemented by 24 MR sub-modules (*MR-Blocks*), for a total of 407 regulatory proteins (Supplemental Data 6,11). MOMA-inferred subtypes provide novel stratification of TCGA cohorts that have been traditionally difficult to study by gene expression profile alone, while MR-Block activity was found to be highly predictive of patient outcome in virtually all cohorts.

On average, the top 33 MR proteins defining a Tumor Checkpoint were sufficient to account for canalization of genomic alterations detected in individual samples. Furthermore, analysis of the 24 MR-Blocks confirmed their role as highly specific, mechanistic regulators of key cancer hallmark programs. Since each sample was analyzed on an individual basis, these results are agnostic to prior tumor classification schemas, as well as to tumor histology and thus constitute a *bona fide*, unbiased pan-cancer analysis of tumorigenic mechanism conservation. To support these findings, we performed experimental *in vitro* and *in vivo* studies of 5 loss-of function mutations that MOMA identified upstream of the worst prognosis subtype Tumor Checkpoint in prostate cancer. Taken together, this suggests that tumor identities based on the activity of MR-Block proteins are likely to complement and extend prior pan-cancer classification schemas by providing more direct relationships between genetics and tumor subtypes.

The MOMA framework can be accessed on Bioconductor (Gentleman et al., 2004), thus allowing analysis of virtually any cancer cohort with patient-matched transcriptional and mutational profiles. In addition, we provide both the Tumor Checkpoint MRs for the 112 tumor subtypes identified by the analysis as well as the MRs in the 24 MR-Blocks. These represent a comprehensive new collection of candidate tumor dependencies and therapeutic targets and outcome/drug-sensitivity biomarkers, several of which have been validated in previous studies, see for instance (Alvarez et al., 2018; Aytes et al., 2014b; Bisikirska et al., 2016; Carro et al., 2010; Rajbhandari et al., 2018b; Walsh et al., 2017). Given the pan-cancer nature of this work, in the following sections we will use different tumor types to highlight key advantages and novel findings made possible by the MOMA framework.

## Results

### Integrative analysis of genetic alterations and transcriptional state identifies pan-cancer MR proteins

The goal of this analysis is to systematically identify MR proteins that implement a tumor’s transcriptional identity by canalizing the effect of genetic alterations in their upstream pathways, for every tumor sample in the TCGA repository. To accomplish this goal, we first transformed the gene expression profile of each sample to a protein activity profile, using the VIPER algorithm (Alvarez et al., 2016). For each sample, we then prioritized the most aberrantly activated proteins as candidate MRs based on the presence of upstream functional mutations, using the DIGGIT algorithm (Chen et al., 2014), see Figure 1 for a conceptual workflow of the analysis.

**Figure 1:**
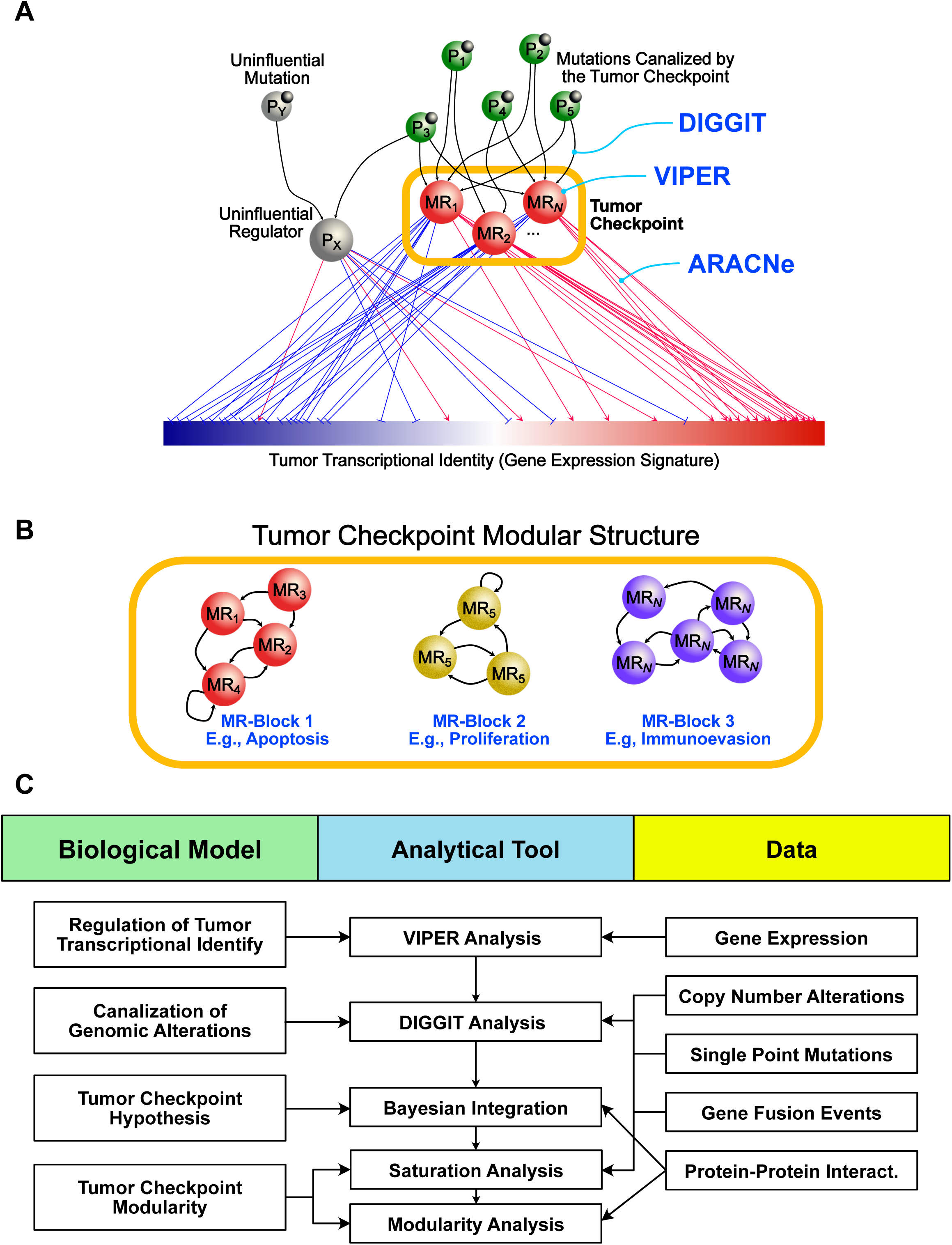
Conceptual overview of the algorithm to find sample “checkpoints” and checkpoint blocks. (A) Diagram illustrating the “bottleneck hypothesis”. Master regulator proteins (‘MR’) integrate the signal from genomic mutations (‘P’) and other “driver” genomic alterations, modulating the “downstream” gene expression signature (red represents upregulated genes, and blue represents downregulated genes). Proteins that assist or co-modulate the signal but are not downstream of genomic alterations, or downstream of only passenger events, are not considered master regulators. The set of master regulator proteins for a given sample is defined as that sample’s “checkpoint”. (B) Checkpoint “blocks” are defined as sets of master regulator proteins (‘MR blocks’) that modulate a specific part of the gene expression signature.. Each sample’s checkpoint may contain several active checkpoint “blocks” that collectively integrate the signal from upstream genomic drivers to modulate the overall gene expression signature. (C) Flow diagram of the inference algorithm to find sample checkpoints and recurrent pan-cancer checkpoint blocks. Multi ‘omics data (gene expression, copy number, SNPs, Fusions and protein-protein interactions) is integrated in a computational pipeline to infer sample checkpoints and checkpoint blocks.

VIPER has been widely validated as an accurate methodology to measure a protein’s transcriptional activity, based on the enrichment of its activated and repressed transcriptional targets (*regulons*) in over and under-expressed genes (see methods) (Alvarez et al., 2016). It is conceptually equivalent to using a multiplexed gene-reporter assay, comprising the transcriptional targets of a protein (i.e., its *regulon*), which are tuned for each specific regulatory protein and each tumor context. We used the ARACNe algorithm (Basso et al., 2005) to dissect accurate regulons for every transcription factor (TF), co-factor (co-TF), and chromatin remodeling enzyme (CRE) (*n* = 2,506). These proteins were selected because they represent the most direct/mechanistic regulators of a cell’s transcriptional state, via physical, on-chromatin interactions. Systematic experimental validation had previously confirmed the accuracy of VIPER activity measurements for >80% of these proteins, including high reproducibility when up to 60% of the targets in a regulon were randomized (Alvarez et al., 2016), thus showing robustness to false positive interactions. Moreover, from other prior studies, on average >70% of ARACNe-inferred targets were validated via biochemical and functional assays, such as Chromatin Immunoprecipitation (ChIP) and RNAi-mediated silencing followed by gene expression profiling—see for instance (Basso et al., 2005; Carro et al., 2010; Lefebvre et al., 2010). This confirms that VIPER not only produces realistic protein activity measurements but also effectively identifies the proteins that mechanistically regulate a sample’s transcriptional state through their physical targets. ARACNe requires *N* ≥100 samples for optimal accuracy, thus restricting the analysis to 20 TCGA cohorts (Table 1), for a total of 9,738 primary tumor samples.

**Table 1:** Data Overview. Data for 20 TCGA tumor types is listed, including the number of samples with RNA, mutational, copy number, and fusion data, respectively.

| TCGA Tissue | # Patients, VIPER Inferences | # Patients, mutation data | # Patients, cnv data | # Patients, fusion calls |
| --- | --- | --- | --- | --- |
| blca | 408 | 130 | 253 | 300 |
| brca | 1093 | 977 | 1051 | 831 |
| coad | 457 | 217 | 437 | 154 |
| gbm | 160 | 291 | 589 | 122 |
| hnsc | 520 | 306 | 513 | 341 |
| kirc | 533 | 293 | 520 | 151 |
| laml | 179 | 196 | 198 | 74 |
| lgg | 516 | 289 | 464 | 0 |
| lihc | 371 | 202 | 193 | 222 |
| luad | 515 | 519 | 496 | 405 |
| lusc | 501 | 178 | 493 | 401 |
| ov | 296 | 141 | 587 | 194 |
| paad | 178 | 168 | 185 | 0 |
| prad | 497 | 425 | 204 | 437 |
| read | 166 | 81 | 164 | 0 |
| sarc | 259 | 255 | 260 | 0 |
| skcm | 468 | 362 | 386 | 338 |
| stad | 274 | 230 | 353 | 183 |
| thca | 501 | 402 | 499 | 144 |
| ucec | 545 | 248 | 543 | 101 |
| cesc | 304 |  |  |  |
| esca | 184 |  |  |  |
| kirp | 290 |  |  |  |
| pcpg | 179 |  |  |  |
| tgct | 150 |  |  |  |
| thym | 120 |  |  |  |

To identify transcriptional tumor identities (i.e., tumor subtypes) implemented by the same subset of regulatory proteins, we performed protein-activity-based unsupervised cluster analysis of the 20 selected TCGA cohorts, using a k-medoids approach (see STAR methods). Within each cohort, the optimal number of clusters was determined using a silhouette-score-based metric (Figure 2A and STAR methods), using the protein activity of the predicted tumor checkpoint proteins. Here we show the 5-cluster optimal solution for KIRC, as an illustrative example (Figure 2B); see Figure S1A-T for all other cohorts. Using the same clustering algorithm (PAM) protein-activity-based clustering significantly outperformed gene-expression-based clustering in all 20 cohorts (*p* ≤ 1.8E-8 in every cohort and *p* < 2.2E-16 in all but one (SKCM), by Wilcoxon rank sum test; see STAR methods, GEX clustering; Figure 2C). Optimal cluster number ranged from *k* = 2 to *k* = 8 per cohort. Whenever two or more statistically-equivalent cluster structures were identified for a given cohort (e.g. *k* = 3 and *k* = 4), we selected the one producing the best association with survival, see Table 2, with twelve cohorts thus further prioritized based on outcome. As an example, we show differential outcome in Cluster 5 (worst) vs. Cluster 3 (best) for KIRC (Figure 2D) (*p* = 1.1E-16).

**Figure 2:**
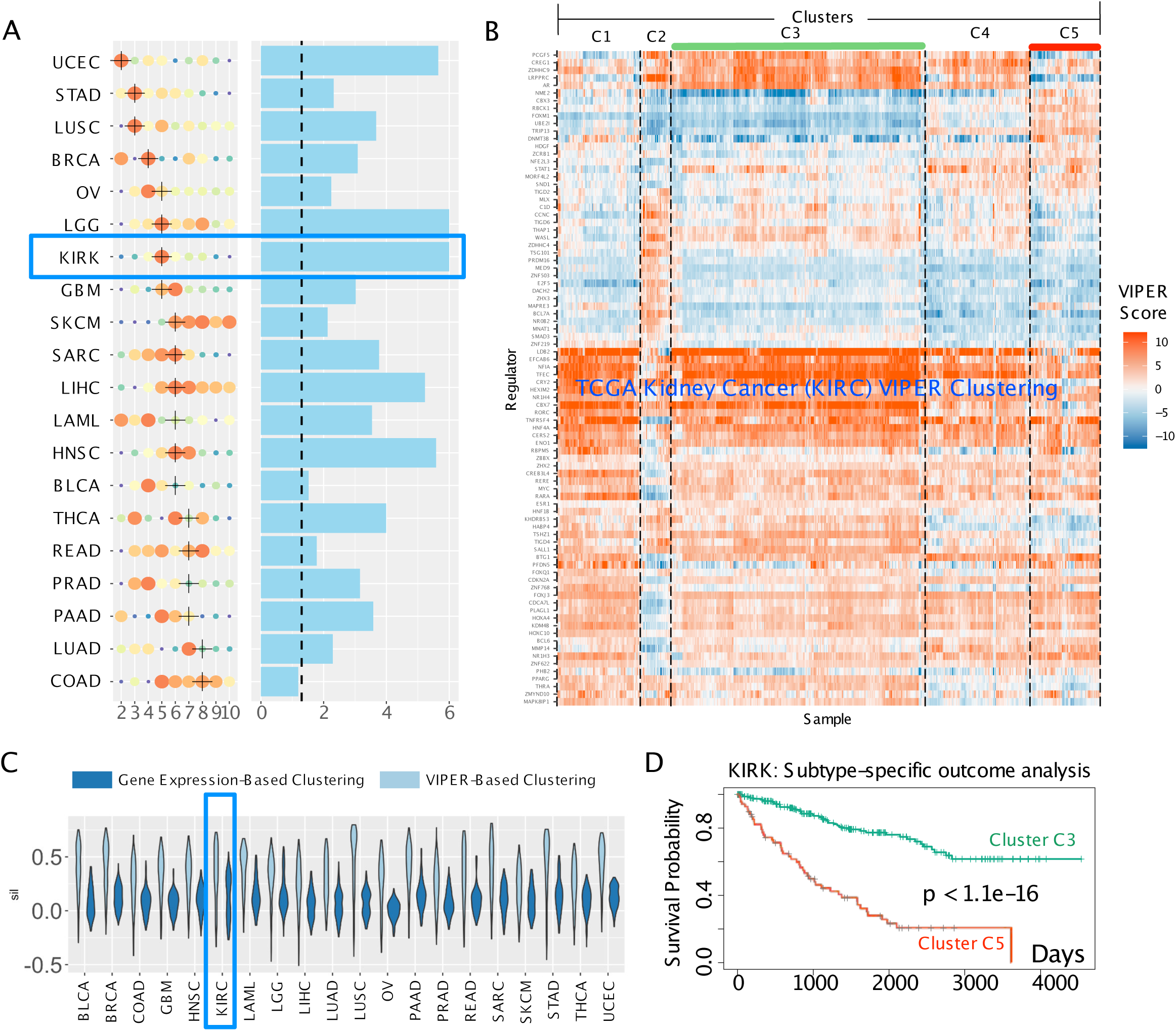
Unsupervised transcriptional subtypes inferred from multi-omics data integration, and associated survival. (A) Analytical score using a modified silhouette measure of each cohort’s cluster fit is shown for each clustering solution from k=2-10, for each tissue type (y-axis). Larger and redder dots represent better mean scores. If multiple statistically equivalent solutions existed the solution with the strongest survival separation between the best and worst surviving clusters was selected (black cross); significance of survival separation is shown in −log_2_(p-value) to the right of each clustering solution (blue bars, X-axis is −log_2_(p-value)). A dashed line represents the canonical threshold for statistical significance (p < 0.05) in log space. (B) Heatmap of VIPER-inferred protein activity for the candidate master regulators of 5 transcriptional subgroups of the TCGA kidney cancer cohort. The best and worst surviving clusters 3 and 5, are highlighted. (C) Violin plots of cluster silhouette scores (y-axis) for each sample, for each of 20 tissue types (x-axis); light blue are VIPER protein activity clusters, dark blue are raw gene expression clustering solutions. (D) Survival probability of patients in unsupervised VIPER-inferred cluster 3 (green solid line) relative to cluster 5 (dashed black line) after fitting a Cox proportional hazards model to the TCGA clinical data (p < 1.1e-16).

**Table 2:** Survival analysis of MOMA sample clustering. Mean silhouette scores, the p-value of survival differences between the best and worst surviving clusters, and the progression-free survival p-value are shown for each clustering solution across the 20 TCGA tumor types.

| tissue | k | mean.silhouette | pval.progression-naive | pval.progression-free | top.analytical.soln |
| --- | --- | --- | --- | --- | --- |
| blca | 2 | 0.608997478 | 0.63232 | 0.37693 | no |
| blca | 3 | 0.706560727 | 0.16872 | 0.11797 | no |
| blca | 4 | 0.759714371 | 0.21336 | 0.26033 | yes |
| blca | 5 | 0.720047253 | 0.1506 | 0.19498 | yes |
| blca | 6 | 0.641998616 | 0.033141 | 0.01937 | yes |
| blca | 7 | 0.69636671 | 0.042014 | 0.027441 | no |
| blca | 8 | 0.633710917 | 0.03032 | 0.0044003 | no |
| blca | 9 | 0.671606656 | 0.051675 | 0.13531 | no |
| blca | 10 | 0.632906191 | 0.18073 | 0.07716 | no |
| brca | 2 | 0.780697516 | 0.013218 | 0.039361 | no |
| brca | 3 | 0.711048776 | 0.11849 | 0.010141 | no |
| brca | 4 | 0.7852366 | 0.020421 | 0.013422 | yes |
| brca | 5 | 0.72477614 | 0.00082055 | 0.0107 | no |
| brca | 6 | 0.719701065 | 0.0051902 | 0.0107 | no |
| brca | 7 | 0.771018746 | 0.0079326 | 0.00025076 | no |
| brca | 8 | 0.759411684 | 0.016372 | 0.033565 | no |
| brca | 9 | 0.723676859 | 0.016372 | 0.00034953 | no |
| brca | 10 | 0.729480844 | 0.0051699 | 0.00049293 | no |
| coad | 2 | 0.646982494 | 0.8179 | 0.8435 | no |
| coad | 3 | 0.686357381 | 0.84978 | 0.086235 | no |
| coad | 4 | 0.645325064 | 0.78718 | 0.18189 | no |
| coad | 5 | 0.722428311 | 0.74365 | 0.095462 | yes |
| coad | 6 | 0.716045021 | 0.4658 | 0.24934 | yes |
| coad | 7 | 0.712651649 | 0.28944 | 0.35243 | no |
| coad | 8 | 0.716932619 | 0.064724 | 0.55033 | yes |
| coad | 9 | 0.714158603 | 0.08393 | 0.11422 | no |
| coad | 10 | 0.698958217 | 0.15085 | 0.41205 | no |
| gbm | 2 | 0.613839743 | 0.48806 | 0.053117 | no |
| gbm | 3 | 0.686719852 | 0.13087 | 0.096051 | no |
| gbm | 4 | 0.639781348 | 0.0068145 | 0.0013133 | no |
| gbm | 5 | 0.737300554 | 0.0039469 | 0.0014076 | yes |
| gbm | 6 | 0.755589927 | 0.0041277 | 0.0023037 | yes |
| gbm | 7 | 0.67755406 | 0.032386 | 0.0067482 | no |
| gbm | 8 | 0.669564142 | 0.032386 | 0.0067482 | no |
| gbm | 9 | 0.652577276 | 0.03294 | 0.013374 | no |
| gbm | 10 | 0.646740114 | 0.00095328 | 0.013374 | no |
| hnsc | 2 | 0.628503024 | 0.23424 | 0.055252 | no |
| hnsc | 3 | 0.667231408 | 0.0049155 | 0.030115 | no |
| hnsc | 4 | 0.703443865 | 0.078984 | 0.019162 | yes |
| hnsc | 5 | 0.722152229 | 0.00012022 | 0.0046603 | no |
| hnsc | 6 | 0.748753543 | 0.00010972 | 0.0079217 | yes |
| hnsc | 7 | 0.744952807 | 0.00035881 | 0.0075361 | no |
| hnsc | 8 | 0.673461776 | 9.25E-05 | 0.0084426 | no |
| hnsc | 9 | 0.647997614 | 0.00010592 | 0.00028122 | no |
| hnsc | 10 | 0.660101673 | 2.61E-06 | 7.89E-05 | no |
| kirc | 2 | 0.589877331 | 7.23E-14 | 1.11E-16 | no |
| kirc | 3 | 0.609216895 | 1.79E-13 | 5.09E-13 | no |
| kirc | 4 | 0.668013519 | 0 | 7.77E-16 | no |
| kirc | 5 | 0.730298651 | 1.11E-16 | 0.00047811 | yes |
| kirc | 6 | 0.67227246 | 6.63E-14 | 5.78E-13 | yes |
| kirc | 7 | 0.648441513 | 0.00066381 | 0.00029486 | no |
| kirc | 8 | 0.649804299 | 0.0006531 | 0.00036198 | no |
| kirc | 9 | 0.59840271 | 4.30E-12 | 6.67E-12 | no |
| kirc | 10 | 0.575458417 | 7.73E-12 | 8.87E-10 | no |
| laml | 2 | 0.801153108 | 0.46044 | NA | no |
| laml | 3 | 0.765486157 | 0.019749 | NA | no |
| laml | 4 | 0.810313106 | 0.0099112 | NA | yes |
| laml | 5 | 0.700628002 | 0.0062179 | NA | yes |
| laml | 6 | 0.71457491 | 0.00090455 | NA | yes |
| laml | 7 | 0.710777817 | 0.00090455 | NA | no |
| laml | 8 | 0.742692413 | 0.00090455 | NA | no |
| laml | 9 | 0.631961571 | 0.0014776 | NA | no |
| laml | 10 | 0.744817443 | 0.00028785 | NA | no |
| lgg | 2 | 0.552213534 | 0 | 0 | no |
| lgg | 3 | 0.677869379 | 0 | 0 | no |
| lgg | 4 | 0.690846507 | 0 | 1.38E-12 | no |
| lgg | 5 | 0.737693354 | 0 | 1.89E-15 | yes |
| lgg | 6 | 0.688394999 | 0 | 0 | no |
| lgg | 7 | 0.704830302 | 0 | 2.22E-16 | yes |
| lgg | 8 | 0.726068521 | 0 | 1.81E-11 | yes |
| lgg | 9 | 0.63463369 | 0 | 1.81E-11 | yes |
| lgg | 10 | 0.676811717 | 0.00011307 | 0.0018331 | no |
| lihc | 2 | 0.561760152 | 5.10E-05 | 1.69E-05 | no |
| lihc | 3 | 0.502660838 | 0.0028464 | 0.0029454 | no |
| lihc | 4 | 0.625995969 | 2.05E-05 | 1.01E-05 | yes |
| lihc | 5 | 0.685422688 | 9.02E-06 | 3.32E-06 | yes |
| lihc | 6 | 0.706792583 | 5.90E-06 | 1.20E-05 | yes |
| lihc | 7 | 0.702901293 | 9.88E-05 | 1.17E-05 | yes |
| lihc | 8 | 0.679974881 | 0.002001 | 2.58E-08 | yes |
| lihc | 9 | 0.677602536 | 0.0053459 | 0.26165 | yes |
| lihc | 10 | 0.667423456 | 0.0024896 | 0.25462 | yes |
| luad | 2 | 0.682103767 | 0.092896 | 0.27301 | no |
| luad | 3 | 0.70962839 | 0.013047 | 0.032815 | no |
| luad | 4 | 0.714583055 | 0.10638 | 0.029843 | yes |
| luad | 5 | 0.612862379 | 0.032239 | 0.029975 | yes |
| luad | 6 | 0.640803279 | 0.022622 | 0.049676 | yes |
| luad | 7 | 0.749596792 | 0.045614 | 0.055438 | yes |
| luad | 8 | 0.656925785 | 0.0051143 | 0.02797 | yes |
| luad | 9 | 0.648309819 | 0.015539 | 0.099825 | yes |
| luad | 10 | 0.639529278 | 0.12293 | 0.019241 | no |
| lusc | 2 | 0.528566491 | 0.19052 | 0.2046 | no |
| lusc | 3 | 0.75505046 | 0.0067527 | 0.0024793 | yes |
| lusc | 4 | 0.676896457 | 0.005613 | 0.0010377 | no |
| lusc | 5 | 0.728830627 | 0.050373 | 0.18249 | no |
| lusc | 6 | 0.631955309 | 0.00027513 | 0.00014193 | no |
| lusc | 7 | 0.688542834 | 0.012909 | 0.0015291 | no |
| lusc | 8 | 0.639857422 | 0.0097389 | 0.00089683 | no |
| lusc | 9 | 0.663310017 | 0.00021324 | 0.02836 | no |
| lusc | 10 | 0.656324988 | 0.0010716 | 0.016821 | no |
| ov | 2 | 0.537390233 | 0.10354 | 0.20396 | no |
| ov | 3 | 0.710201841 | 0.015154 | 0.18232 | yes |
| ov | 4 | 0.782126608 | 0.1204 | 0.22822 | yes |
| ov | 5 | 0.725069551 | 0.0057372 | 0.3668 | yes |
| ov | 6 | 0.6692378 | 0.038902 | 0.34226 | no |
| ov | 7 | 0.669860496 | 0.18572 | 0.31972 | no |
| ov | 8 | 0.666231449 | 0.073733 | 0.24804 | no |
| ov | 9 | 0.648926518 | 0.18572 | 0.28631 | no |
| ov | 10 | 0.666539449 | 0.11984 | 0.26974 | no |
| paad | 2 | 0.769596412 | 0.010689 | 0.0018173 | no |
| paad | 3 | 0.681340558 | 0.00093584 | 5.65E-05 | yes |
| paad | 4 | 0.689516332 | 0.0069061 | 0.0013505 | no |
| paad | 5 | 0.790259533 | 0.0051506 | 0.0034655 | yes |
| paad | 6 | 0.774964909 | 0.0051506 | 0.0034655 | no |
| paad | 7 | 0.756527477 | 0.00026206 | 0.00033836 | yes |
| paad | 8 | 0.681827078 | 0.00035612 | 0.00033836 | no |
| paad | 9 | 0.697770396 | 0.00045162 | 0.00015924 | no |
| paad | 10 | 0.675166787 | 0.00045162 | 0.00015924 | no |
| prad | 2 | 0.564812805 | 0.064563 | 0.001729 | no |
| prad | 3 | 0.681793239 | 0.083578 | 0.002533 | yes |
| prad | 4 | 0.695943936 | 0.16174 | 0.0066142 | yes |
| prad | 5 | 0.648679804 | 0.016743 | 0.0040458 | yes |
| prad | 6 | 0.656139071 | 0.0020547 | 0.058516 | yes |
| prad | 7 | 0.603397574 | 0.00069596 | 0.02439 | yes |
| prad | 8 | 0.625423616 | 0.0070901 | 0.067889 | no |
| prad | 9 | 0.604941287 | 0.0016917 | 7.55E-05 | yes |
| prad | 10 | 0.622967345 | 0.0029194 | 0.00076128 | yes |
| read | 2 | 0.576258894 | 0.14116 | 0.95182 | no |
| read | 3 | 0.663345981 | 0.16911 | 0.1743 | no |
| read | 4 | 0.666606726 | 0.030973 | 0.12663 | no |
| read | 5 | 0.677701654 | 0.23047 | 0.22067 | yes |
| read | 6 | 0.651477825 | 0.36028 | 0.22067 | yes |
| read | 7 | 0.673415931 | 0.016712 | 0.1573 | yes |
| read | 8 | 0.684067301 | 0.18536 | 0.083265 | yes |
| read | 9 | 0.64772557 | 0.21887 | 0.22067 | yes |
| read | 10 | 0.646137155 | 0.24821 | 0.22067 | yes |
| sarc | 2 | 0.661718753 | 0.18972 | 0.28819 | no |
| sarc | 3 | 0.7254593 | 0.0010793 | 0.1934 | yes |
| sarc | 4 | 0.742160411 | 0.005309 | 0.11886 | yes |
| sarc | 5 | 0.752197064 | 0.0094806 | 0.086744 | yes |
| sarc | 6 | 0.757693054 | 0.0001718 | 0.073535 | yes |
| sarc | 7 | 0.722452491 | 0.0019067 | 0.053358 | yes |
| sarc | 8 | 0.643407666 | 0.00082503 | 0.018268 | yes |
| sarc | 9 | 0.635483392 | 0.0001842 | 0.010914 | no |
| sarc | 10 | 0.612152076 | 0.0098517 | 0.066778 | no |
| skcm | 2 | 0.564156317 | 0.26587 | 0.14977 | no |
| skcm | 3 | 0.55969847 | 0.35076 | 0.088105 | no |
| skcm | 4 | 0.565191251 | 0.038617 | 0.12356 | no |
| skcm | 5 | 0.590803727 | 0.0075082 | 0.092109 | no |
| skcm | 6 | 0.64396706 | 0.0094092 | 0.11865 | yes |
| skcm | 7 | 0.645226795 | 0.029668 | 0.179 | yes |
| skcm | 8 | 0.648586086 | 0.029668 | 0.12731 | yes |
| skcm | 9 | 0.635180732 | 0.061822 | 0.18792 | yes |
| skcm | 10 | 0.646480167 | 0.011415 | 0.025271 | yes |
| stad | 2 | 0.723922828 | 0.0088738 | 0.0066771 | no |
| stad | 3 | 0.785824145 | 0.0048198 | 0.0015581 | yes |
| stad | 4 | 0.71885937 | 0.043995 | 0.016671 | yes |
| stad | 5 | 0.739588128 | 0.038258 | 0.030502 | yes |
| stad | 6 | 0.729738863 | 0.18979 | 0.07219 | yes |
| stad | 7 | 0.694404258 | 0.1144 | 0.032081 | no |
| stad | 8 | 0.639886237 | 0.10595 | 0.062994 | no |
| stad | 9 | 0.608928271 | 0.017654 | 0.032331 | no |
| stad | 10 | 0.588086261 | 0.032527 | 0.00015066 | no |
| thca | 2 | 0.650162933 | 0.69403 | 0.75789 | no |
| thca | 3 | 0.697119516 | 0.054398 | 0.29184 | no |
| thca | 4 | 0.640746837 | 0.10751 | 0.28275 | no |
| thca | 5 | 0.609452281 | 0.0081927 | 0.083265 | no |
| thca | 6 | 0.699652501 | 0.0014467 | 0.25926 | yes |
| thca | 7 | 0.647106832 | 0.00010161 | 0.22067 | yes |
| thca | 8 | 0.689565794 | 0.00037157 | 0.26355 | no |
| thca | 9 | 0.63177716 | 0.00039194 | 0.34957 | no |
| thca | 10 | 0.614423854 | 0.00066888 | 0.52709 | no |
| ucec | 2 | 0.742400826 | 1.72E-05 | 4.06E-05 | yes |
| ucec | 3 | 0.68008393 | 2.23E-06 | 0.00048987 | no |
| ucec | 4 | 0.712390962 | 0.0028716 | 0.0022609 | no |
| ucec | 5 | 0.707244791 | 0.0030273 | 0.0023555 | no |
| ucec | 6 | 0.647148064 | 0.019392 | 0.056942 | no |
| ucec | 7 | 0.677781261 | 0.0050285 | 0.027677 | no |
| ucec | 8 | 0.714713798 | 0.0055951 | 0.024932 | no |
| ucec | 9 | 0.668821752 | 0.022302 | 0.0042407 | no |
| ucec | 10 | 0.635964719 | 0.021962 | 0.0036179 | no |

In total, the analysis identified 112 clusters, representing a novel stratification of cancer into distinct transcriptional identities, each one mechanistically regulated by a specific subset of regulatory proteins (Figure 2B, S1A-T, and Table 3, Supplemental Data 4). Supporting the value and novelty of the classification, this analysis identified differential outcome subtypes in TCGA cohorts that had been previously challenging in terms of gene-expression-based stratification, such as prostate cancer. In addition, for each subtype, the analysis provided a repertoire of MR proteins representing its most likely mechanistic determinants. As discussed in the following, this also provides direct links to the specific genetic alterations that, by affecting proteins in upstream pathways, induce aberrant MR-protein activity on an individual sample basis.

**Table 3:** MOMA subtype summary. Cluster identities, sample size and fractions are shown for each of the 20 TCGA tissue types.

| Organ Site | Subtype | Sample Count | Fraction | Subtype Total | Site ID |
| --- | --- | --- | --- | --- | --- |
| blca | 1 | 93 | 23% | 408 | 1 |
| blca | 2 | 46 | 11% |  |  |
| blca | 3 | 81 | 20% |  |  |
| blca | 4 | 77 | 19% |  |  |
| blca | 5 | 62 | 15% |  |  |
| blca | 6 | 49 | 12% |  |  |
| brca | 1 | 337 | 31% | 1100 | 2 |
| brca | 2 | 315 | 29% |  |  |
| brca | 3 | 222 | 20% |  |  |
| brca | 4 | 226 | 21% |  |  |
| coad | 1 | 56 | 12% | 459 | 3 |
| coad | 2 | 50 | 11% |  |  |
| coad | 3 | 31 | 7% |  |  |
| coad | 4 | 78 | 17% |  |  |
| coad | 5 | 105 | 23% |  |  |
| coad | 6 | 87 | 19% |  |  |
| coad | 7 | 38 | 8% |  |  |
| coad | 8 | 14 | 3% |  |  |
| gbm | 1 | 25 | 15% | 166 | 4 |
| gbm | 2 | 8 | 5% |  |  |
| gbm | 3 | 31 | 19% |  |  |
| gbm | 4 | 68 | 41% |  |  |
| gbm | 5 | 34 | 20% |  |  |
| hnsc | 1 | 59 | 11% | 522 | 5 |
| hnsc | 2 | 93 | 18% |  |  |
| hnsc | 3 | 157 | 30% |  |  |
| hnsc | 4 | 64 | 12% |  |  |
| hnsc | 5 | 81 | 16% |  |  |
| hnsc | 6 | 68 | 13% |  |  |
| kirc | 1 | 82 | 15% | **534** | **6** |
| kirc | 2 | 30 | 6% |  |  |
| kirc | 3 | 250 | 47% |  |  |
| kirc | 4 | 103 | 19% |  |  |
| kirc | 5 | 69 | 13% |  |  |
| laml | 1 | 21 | 12% | **179** | **7** |
| laml | 2 | 66 | 37% |  |  |
| laml | 3 | 22 | 12% |  |  |
| laml | 4 | 24 | 13% |  |  |
| laml | 5 | 31 | 17% |  |  |
| laml | 6 | 15 | 8% |  |  |
| lgg | 1 | 221 | 42% | **530** | **8** |
| lgg | 2 | 87 | 16% |  |  |
| lgg | 3 | 25 | 5% |  |  |
| lgg | 4 | 166 | 31% |  |  |
| lgg | 5 | 31 | 6% |  |  |
| lihc | 1 | 81 | 22% | **373** | **9** |
| lihc | 2 | 66 | 18% |  |  |
| lihc | 3 | 59 | 16% |  |  |
| lihc | 4 | 60 | 16% |  |  |
| lihc | 5 | 67 | 18% |  |  |
| lihc | 6 | 40 | 11% |  |  |
| luad | 1 | 41 | 8% | **517** | **10** |
| luad | 2 | 47 | 9% |  |  |
| luad | 3 | 52 | 10% |  |  |
| luad | 4 | 67 | 13% |  |  |
| luad | 5 | 92 | 18% |  |  |
| luad | 6 | 40 | 8% |  |  |
| luad | 7 | 135 | 26% |  |  |
| luad | 8 | 43 | 8% |  |  |
| lusc | 1 | 134 | 27% | **501** | **11** |
| lusc | 2 | 320 | 64% |  |  |
| lusc | 3 | 47 | 9% |  |  |
| ov | 1 | 62 | 21% | **299** | **12** |
| ov | 2 | 40 | 13% |  |  |
| ov | 3 | 96 | 32% |  |  |
| ov | 4 | 62 | 21% |  |  |
| ov | 5 | 39 | 13% |  |  |
| paad | 1 | 71 | 40% | **179** | **13** |
| paad | 2 | 25 | 14% |  |  |
| paad | 3 | 14 | 8% |  |  |
| paad | 4 | 22 | 12% |  |  |
| paad | 5 | 23 | 13% |  |  |
| paad | 6 | 9 | 5% |  |  |
| paad | 7 | 15 | 8% |  |  |
| prad | 1 | 149 | 30% | 498 | 14 |
| prad | 2 | 47 | 9% |  |  |
| prad | 3 | 109 | 22% |  |  |
| prad | 4 | 38 | 8% |  |  |
| prad | 5 | 86 | 17% |  |  |
| prad | 6 | 9 | 2% |  |  |
| prad | 7 | 60 | 12% |  |  |
| read | 1 | 64 | 38% | 167 | 15 |
| read | 2 | 38 | 23% |  |  |
| read | 3 | 11 | 7% |  |  |
| read | 4 | 27 | 16% |  |  |
| read | 5 | 9 | 5% |  |  |
| read | 6 | 13 | 8% |  |  |
| read | 7 | 5 | 3% |  |  |
| sarc | 1 | 50 | 19% | 263 | 16 |
| sarc | 2 | 41 | 16% |  |  |
| sarc | 3 | 49 | 19% |  |  |
| sarc | 4 | 71 | 27% |  |  |
| sarc | 5 | 29 | 11% |  |  |
| sarc | 6 | 23 | 9% |  |  |
| skcm | 1 | 83 | 18% | 472 | 17 |
| skcm | 2 | 131 | 28% |  |  |
| skcm | 3 | 67 | 14% |  |  |
| skcm | 4 | 119 | 25% |  |  |
| skcm | 5 | 44 | 9% |  |  |
| skcm | 6 | 28 | 6% |  |  |
| stad | 1 | 139 | 51% | 274 | 18 |
| stad | 2 | 74 | 27% |  |  |
| stad | 3 | 61 | 22% |  |  |
| thca | 1 | 64 | 13% | 509 | 19 |
| thca | 2 | 55 | 11% |  |  |
| thca | 3 | 89 | 17% |  |  |
| thca | 4 | 94 | 18% |  |  |
| thca | 5 | 55 | 11% |  |  |
| thca | 6 | 27 | 5% |  |  |
| thca | 7 | 125 | 25% |  |  |
| ucec | 1 | 283 | 52% | 546 | 20 |
| ucec | 2 | 263 | 48% |  |  |

As previously reported, identification of tumor subtypes that effectively associate with clinical and other phenotypic properties by gene expression analysis has often been challenging. For instance, with the exception of the neuroendocrine subtype, outcome stratification of prostate cancer cohorts by gene expression profile analysis has been elusive. In contrast, MOMA identified transcriptional clusters strongly associated with outcome in all of the 20 cohorts (Figure 2A), except for COAD, where the *p*-value was just slightly above statistical significance (*p* = 0.07, by Kaplan Meier). Combined with the highly significant improvement in cluster statistics (i.e., *cluster tightness*), this suggests that protein-activity-based clustering significantly outperforms a directly comparable gene-expression-based PAM cluster analysis (Figure 2C). In addition, it provides a far more compact and interpretable subtype stratification, by replacing differential expression signatures comprising thousands of genes with just a handful of their transcriptional regulators.

While producing a largely novel subtype architecture, VIPER-based clustering also showed concordance with the most established molecular subtypes. In breast cancer, for instance, the four protein-activity based clusters were highly concordant with established molecular subtypes (Figure S2A, *p* = 2.2E-16 by Chi^2^ test). Similarly, in high-grade glioma, we found highly significant concordance (*p* = 2.2E-16; by χ^2^ analysis) with published subtypes (Brennan et al., 2013b) and similar outcome differences between clusters associated with best and worst progression-free survival (*p* = 1.4E-3; Figure S2B). Indeed, in agreement with prior literature, the worst survival cluster was comprised almost entirely of mesenchymal tumors, while the best surviving cluster was predominantly comprised of proneural tumors (*p* = 1.3E-3 and *p* = 3E-6 by FET, respectively) (Brennan et al., 2013a; Carro et al., 2010; Chen et al., 2014). Even though MOMA analysis is fully unsupervised, results were consistent with previous supervised analyses in glioblastoma and prostate cancer, for instance, where samples corresponding to poorest and best outcome had been directly compared. Specifically, CEBPβ/CEBPδ/STAT3 and FOXM1/CENPF—previously validated as synergistic Master Regulators of the most aggressive subtypes of GBM (Carro et al., 2010) and prostate cancer (Aytes et al., 2014a), respectively—were among the top MR proteins identified by MOMA for the PRAD and GBM subtypes associated with worst prognosis. See Figure S2C for differential CEBPβ/CEBPδ/STAT3 activity in cluster 2 (mesenchymal) and cluster 3 (proneural) GBM. This is especially noteworthy, since the poor prognosis subtype in PRAD includes only nine samples, a result of the TCGA tissue selection criteria.

To further prioritize MR-proteins based on the genetic alterations that determine their aberrant activation, we computed a *genomic score* based on the enrichment of genomic alteration in their upstream pathways—on a sample-by-sample basis—using the DIGGIT algorithm (Alvarez et al., 2015; Chen et al., 2014; Torres-Garcia et al., 2014). This includes three steps. First candidate modulators of MR activity were identified by the CINDy algorithm (Giorgi et al., 2014). Further improving the original MINDy algorithm (Wang et al., 2009), CINDy uses the Conditional Mutual Information (CMI) between MRs, their downstream targets and potential upstream modulators, to identify *MR-modulator* proteins, whose abundance is associated with differential MR activity. Activity Quantitative Trait Locus (aQTL) analysis was then used to determine whether genetic alterations in CINDy-inferred MR-modulators were effectively associated with their differential activity. Finally, conditional analysis was used to assess which ones of the aQTLs identified by the analysis were statistically independent of other aQTLs, thus efficiently distinguishing between driver and passenger alterations (e.g., same-amplicon genes with no functional effect on the MR). DIGGIT was shown not only to recapitulate known driver mutations but also to infer novel, highly penetrant mutations that were missed by traditional approaches and were then experimentally validated (Chen et al., 2014).

Finally, to generate a refined repertoire of MR proteins that are responsible for determining a tumor’s transcriptional identity by canalizing the effect of genetic alterations in their upstream pathways, we used a Bayesian evidence integration approach. Specifically, we ranked MRs by integrating evidence from (a) their protein activity (*VIPER score*), (b) their upstream genetic alterations (*genomic score*), and (c) additional structure and literature-based evidence supporting direct protein-protein interactions between the MRs and their MR-modulators harboring genetic alterations, such as the PrePPI algorithm (Zhang et al., 2012) (see STAR Methods*, Integrated Rankings*).

### Identification of MR-proteins in Tumor Checkpoints

We have defined Tumor Checkpoint modules as the minimum repertoire of regulatory proteins necessary to implement a tumor’s transcriptional identity by canalizing the effect of upstream genomic events (i.e., mutations, copy number alterations, etc.). Based on this definition, we used saturation analysis to identify Tumor Checkpoint MRs from the full ranked-list of aberrantly activated proteins, for each of the 112 subtypes (Figure 3A,B). Specifically, this was accomplished by assessing how many of the most aberrantly activated proteins are needed to capture a substantial proportion of, or saturate, the number of genomic alterations they canalize. If, as postulated, Tumor Checkpoints comprise only a handful of MRs, saturation should occur rapidly. In contrast, if mutations were randomly distributed across all proteins, saturation would be gradual.

**Figure 3:**
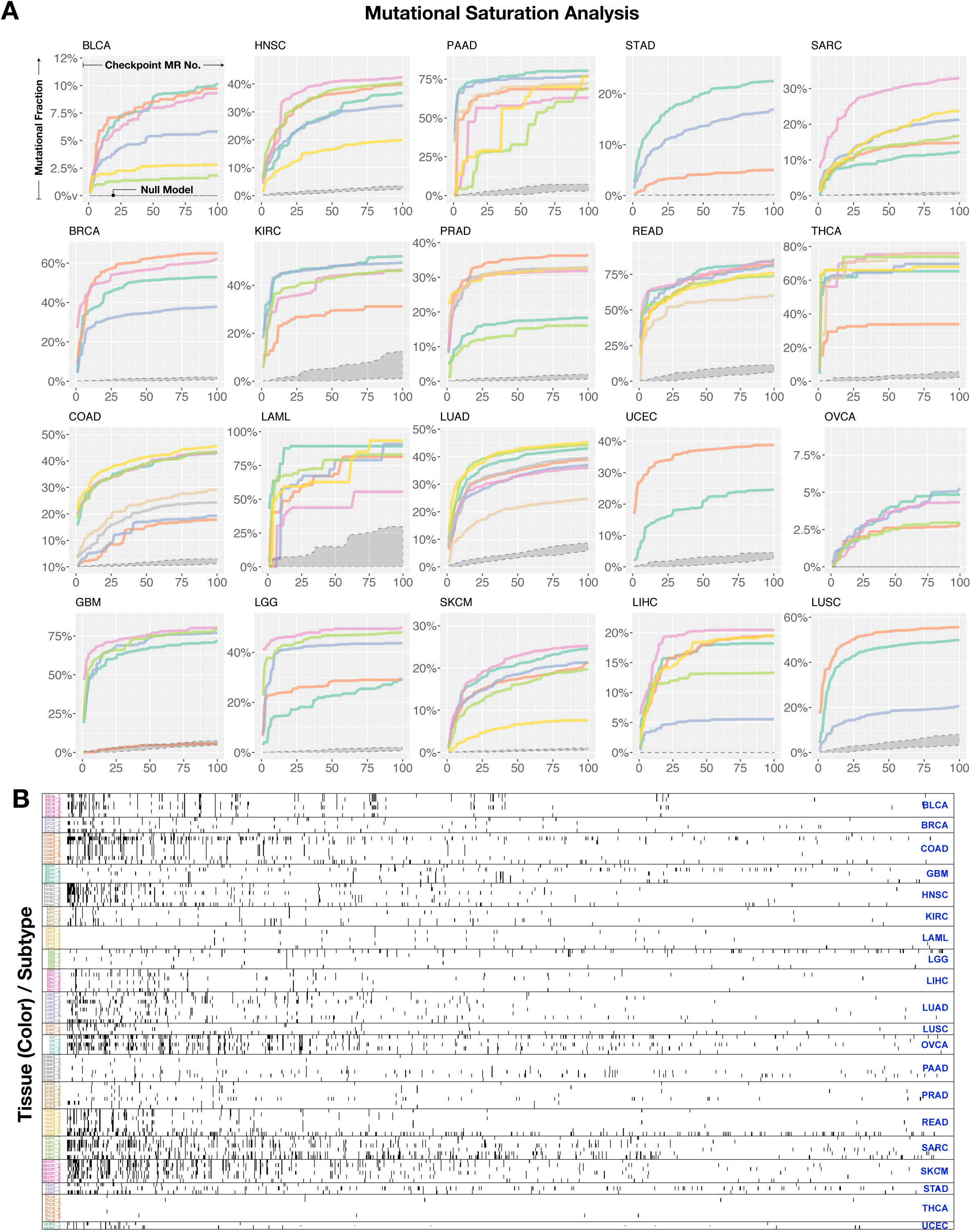
Genomic saturation analysis of candidate master regulators across all tissues of origin. (A) The mean fraction of genomic copy number, SNP and fusions events in each patient (y-axis) and linked candidate Master Regulators (x-axis) is shown as a separate curve for each transcriptional subgroup. Vertical dashed lines indicate the saturation point covering 85% of all events associated with some candidate Master Regulators or the estimated inflection point. (see figure S4; methods). (B) Identities of Master Regulators derived from the saturation analysis in (A) are shown as black tick marks for each transcriptional subtype (row). Color of the y-axis subtype labels represents tissue of origin. Columns (Master Regulators) are sorted by frequency of recurrence in multiple subtypes, from left (highest) to right (lowest). Grey ribbons at the bottom of the plots represent the null-model genomic coverage for 1000 randomly chosen transcription and co-transcription factors that were not ranked in the top 50% by the MOMA algorithm.

To test this hypothesis, we first identified all proteins harboring genetic alterations detected by GISTIC2.0 (Mermel et al., 2011) and non-silent SNVs in a specific subtype—including functional CNVs significant associated with differential gene expression, non-silent SNVs, and focal SCNAs. We then assessed how many of these occurred in CINDy-inferred modulators of the *N* most statistically significant MRs (on a sample-by-sample basis), as ranked by the previously described Bayesian evidence integration, with *N* ranging from 1 to 100. Finally, we plotted both the fraction and total number of mutations as a function of *N*, averaged over all samples in the subtype (Figure 3A).

Consistent with the Tumor Checkpoint hypothesis, we observed extremely rapid saturation of the genetic events canalized by the top MR proteins, across virtually all 112 subtypes, (Figure 3A). For each subtype, we estimated the inflection point of these saturation curves using a simple heuristic (see STAR methods) and found that only a handful of MRs were required to virtually saturate the vast majority of mutations in individual samples. This ranged from 4 MRs (THCA subtype 6) to 86 (LAML subtype 3), with Ovarian cancer representing an outlier with 170, 140, and 140 MRs in subtypes 1,3 and 4, respectively. The latter is likely due to the extremely large number of structural events in this tumor.

Between 14 (0.6%) and 52 (2%) MRs were sufficient to account for the first and third quantile of the mutational burden of each sample and a median of 33 (1.3%) MRs per Tumor Checkpoint. In contrast, when MRs were chosen at random from all 2,506 regulatory proteins, saturation increased very gradually, with no evidence of ever reaching a plateau. Specifically, on average, only 0.4% of the mutations/fusions/CNVs were found upstream of the first 130 (5%) randomly selected MRs (Figure 3A). This confirmed that rapid saturation observed upstream of inferred MRs does not arise from lack of analysis specificity.

At the saturation inflection point, the ratio of genomic events to MRs ranged from r = 0.02 (i.e., one event affecting 50 MRs) to r = 32 (i.e., 32 events affecting a single MR), with an average of 5 events per MR. This is consistent with the hypothesis that the handful of MR-proteins in each Tumor Checkpoints represent critical regulatory bottlenecks, responsible for canalizing the effect of multiple functional mutations (Supplemental Data 4,6). Once saturation was achieved, about half (50%) of all mutations were reported upstream of top MR proteins. Remaining events likely are either non-functional (passenger), too infrequent to be effectively analyzed, or false negatives (i.e., proteins that the analysis failed to identify as MR-modulators). The most significant mutations for each subtype are shown in Supplemental Data 10.

Taken together, these data strongly support the Oncotecture hypothesis and suggest that a much larger and finer-grain mutational repertoire than previously suspected may functionally affect MR-protein activity and, through them, tumor transcriptional identity. In kidney cancer (KIRC), for instance, the analysis identified between 15 and 45 MRs for each of five transcriptional subtypes (Figure 4A-E). These accounted for 40% to 55% of the total number of non-silent SNV and focal GISTIC2.0-detected SCNAs in individual samples of each respective subtype (Figure 4F-J), suggesting significant intertumoral genomic heterogeneity. Specifically, between 40 and 80 genomic alterations per sample were identified as functional determinants of KIRC MR dysregulation. Interestingly, the genetic alterations identified for each subtype are highly distinct, both in terms of their type (e.g., amplifications vs. deletions, Figure 4A,B) and identity. As purely illustrative examples, for instance, TSC1 deletions were detected in >50% of subtype 4 and 5 samples, but only in <30% of subtypes 1, 2, and 3 samples; similarly, BRAF amplifications were detected exclusively in subtype 4 and 5, while KRAS amplifications were exclusive to subtype 5. Such highly subtype-specific mutational landscape co-segregation is pervasive across all tumor cohorts (Figure S4A-T).

**Figure 4:**
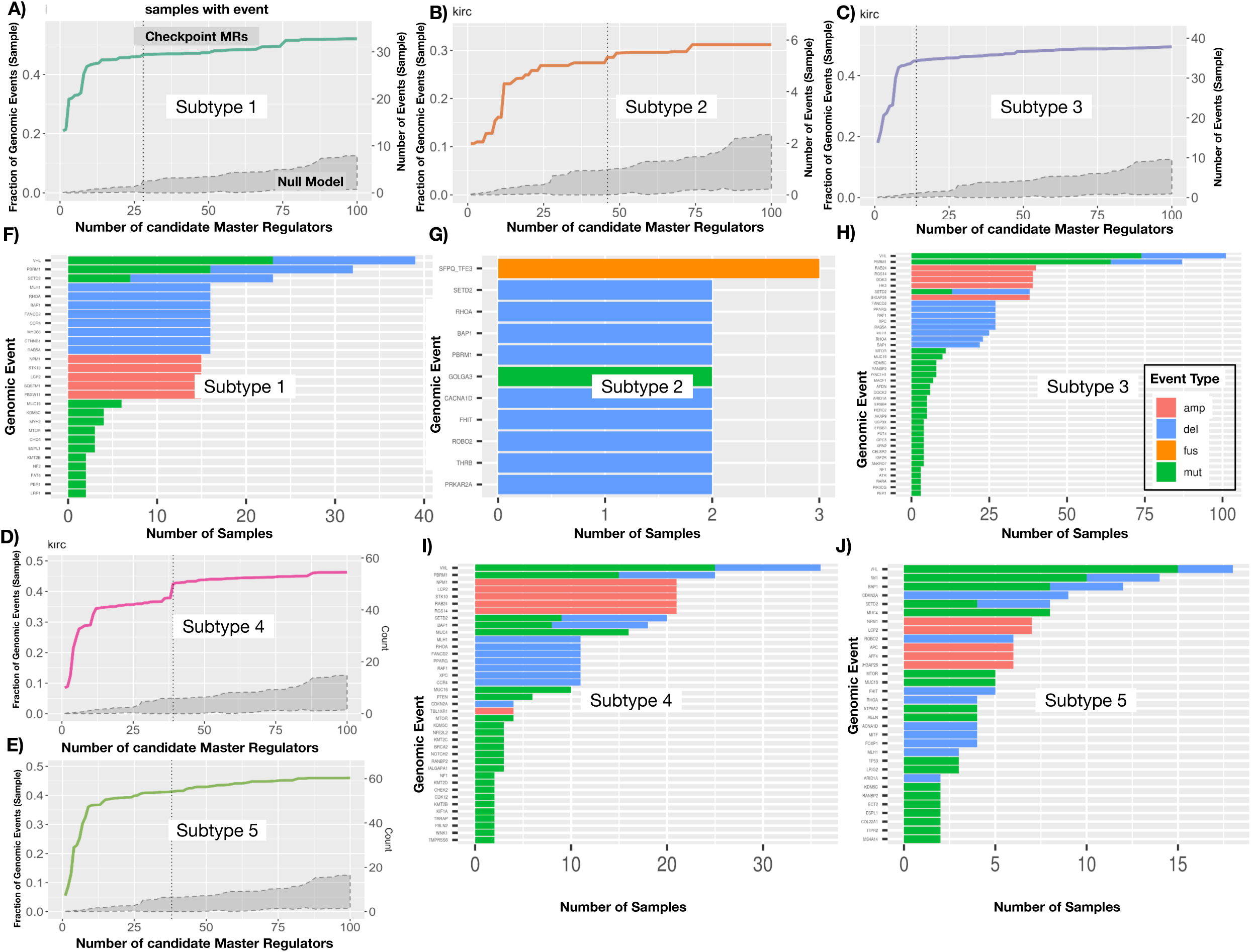
Genomic events upstream of KIRC transcriptional subtype checkpoints. (A-E) Genomic saturation curves for KIRC transcriptional subtypes 1-5; dashed line indicates the point where 85% of all events associated with some Master Regulator are covered, which defines the MR checkpoint for each subtype. Grey ribbons at the bottom of the plots represent the null-model genomic coverage for 1000 randomly chosen transcription and co-transcription factors that were not ranked in the top 50% by the MOMA algorithm. (F-J) Frequency bar plots of genomic events found in samples of each subtype that are downstream of each set of checkpoint MRs, respectively. The number of samples within each subtype with that alteration is plotted on the x-axis; genomic location or gene name is indicated on the y-axis, with studied cancer driver genes indicated if located within a focal amplification or deletion. Bar colors indicate the type of event (focal deletion: blue; focal amplification: red; fusion: yellow; mutation: green).

It should be noted that this analysis is not meant to identify all genetic alterations but rather those that functionally contribute, mechanistically or stochastically, to implementing distinct transcriptional identities. For instance, TP53 mutations, which are completely ubiquitous in ovarian cancer—thus providing no specific contribution to implementing individual transcriptional identities in this cohort—are not reported (Figure S4L). In addition, the proposed cluster analysis may over-stratify some cohorts, to avoid missing rare, molecularly distinct subtypes or subtypes where largely overlapping MR proteins are dysregulated by different genetic events. For instance, in PRAD (Figure S1O), the most aggressive subtype (C6) would be missed due to its small size if a smaller clustering solution were selected. As a result, at first sight, cluster C3 and C7 may appear similar in terms of their MR activity and suggest overstratification. However, closer inspection of the mutational events that co-segregate in these subtypes (Figure S4N) shows that C3 is dominated by TMPRS-ERG fusion events, PTEN mutations and deletions, and ERG, RB1, FOXO1, and SORBS3 deletions. In contrast, C7 is largely devoid of TMPRS-ERG fusions and is instead most enriched in ZNF292, SYNCRIP, MAP3K7, SNX14 deletions and SPOP mutations, suggesting that, albeit similar, their transcriptional identity is driven by an almost orthogonal mutational landscape. In rare cases, subtypes with largely overlapping MR activity and mutational events may be inferred, due to overstratification, as we observed to some degree with pancreatic adenocarcinoma (PAAD), finding high similarity between the mutational events and MR checkpoints of subtypes 3,4 and 5 (Figure S4M). This, however, is not surprising, given the complexity of identifying a common strategy to analyze highly heterogenous cluster structures, as well as the known complexity of pancreatic cancer stratification (Birnbaum et al., 2017).

Finally, there may be biologically relevant subtypes that are missed at the selected level of clustering granularity. For instance, in breast cancer, we identify a basal-like cluster (C4), a Luminal-B enriched cluster (C2), and two Luminal-A clusters (C1 and C3). However, while a more granular 8-cluster solution splits Claudin low/high expressing subtypes in basal cancers (Fig. S1-V), HER2 positive tumors are split between C2 (HR+) and C4 (HR-) and are not identified as forming distinct sub-clusters (Fig. S1-C). This suggests that while HER2+ tumors may present a classic oncogene dependency, their transcriptional identity is actually consistent with that of other basal and Luminal B breast cancers. This highlights the complementarity of this approach, whereas drugs targeting oncogene dependencies would benefit from mutational analysis, while drugs targeting core identity-based dependencies, may target Luminal B HER2+ and Luminal B HER2-with the same approach. Since manual selection of the number of clusters is possible in MOMA, one can explore different clustering solutions to identify the one that makes the most biological/clinical sense. This is of course best accomplished at the individual tumor level rather than across all tumors.

To estimate MOMA’s ability to differentiate between likely driver and passenger mutations, we computed the differential enrichment of mutations upstream of MRs in either GISTIC2.0/CHASM predicted events or all genomic events. When averaged across all MOMA-inferred subtypes of a specific TCGA cancer cohort, differential enrichment of GISTIC2.0 events—i.e., focal amplifications and deletions (confidence 99%)—and significant CHASM events (*p* < 0.05) was highly statistically significant across all but one (LAML) of the tumor subtypes (*p* = 1E-7 to *p =* 1E-156, Figure S3A,B). Our data suggest that low SNV and high fusion-event rates, may have contributed to the LAML discrepancy, since CHASM only assesses candidate SNVs. Even though the vast majority of inferred events were novel, MOMA also effectively recovered all 200 high confidence, pancancer driver genes harboring genetic alterations, as recently identified (Bailey et al., 2018), as well as 92%-100% of the tissue-specific, high-confidence driver genes (98.8%, on average; Supplemental Data 5).

A key novelty of the approach is that it effectively co-segregated genetic alterations—both novel and previously reported—with tumor subtypes, while identifying the specific MR proteins dysregulated by these events and thus responsible for canalizing their effect. Additionally, MOMA inferred a large number of mutational events missed by CHASM and GISTIC, suggesting that the actual repertoire of functional alterations contributing to a tumor’s transcriptional identity may be much larger than previously suspected. See Table 3, and Supplemental Data 4 for a complete account of MOMA-inferred Tumor Checkpoints and MRs, and Figures S4A-U for Master Regulator saturation analysis and upstream genomic event for each of the 112 subtypes. For convenience, we labeled individual Tumor Checkpoints using their two most significant MRs or, when possible, using experimentally validated MRs (e.g., CEBPβ/δ-STAT3 for subtype 2 of high-grade glioma (Carro et al., 2010)).

### Tumor Checkpoints are hyperconnected and modular

Analysis of MOMA-inferred MRs shows that Tumor Checkpoints represent hyperconnected modules of regulatory proteins. This was assessed based on literature-curated regulatory and signaling networks, including HumanNet 2.0 (Hwang et al., 2018) (*p* < 5.0E-42, by Kolmogorov-Smirnov) and Multinet (Khurana et al., 2013) (*p* < 2.0E-37) (Figure S3C,D), as well as on protein-protein interactions predicted by PrePPI using 3D-structure information (Zhang et al., 2012) (*p* = 9.0E-44) (Figure S3E), compared to equal-size sets of regulatory proteins selected at random, as a null model.

To further explore Tumor Checkpoint modularity, we tested whether MR sub-modules could be recurrently identified across multiple Tumor Checkpoints, suggesting the existence of pan-cancer, core regulatory structures (*MR-Blocks* or MRB for short). To accomplish this goal, we first identified and then clustered a subset of recurrent MR proteins included in at least 4 of 112 MOMA-inferred Tumor Checkpoints—a statistically significant threshold based on a random permutation null model (Figure S5A). From the analysis, *k* = 24 MR-Blocks emerged as the optimal solution (Figure 5A, Figure S5B), providing an initial tessellation, where each recurrently-inferred MR was assigned to one and only one MR-Block. To allow a more biologically plausible solution, we then used a “fuzzy” clustering method (Miyamoto et al., 2008) (see STAR methods, fuzzy clustering). such that individual MRs could be included in more than one MR-Block, see Supplemental Data 6. Clustering parameters were optimized to ensure uniqueness and specificity of the MR-Block solution (see methods *Checkpoint Generation*; Figure S5C)

**Figure 5:**
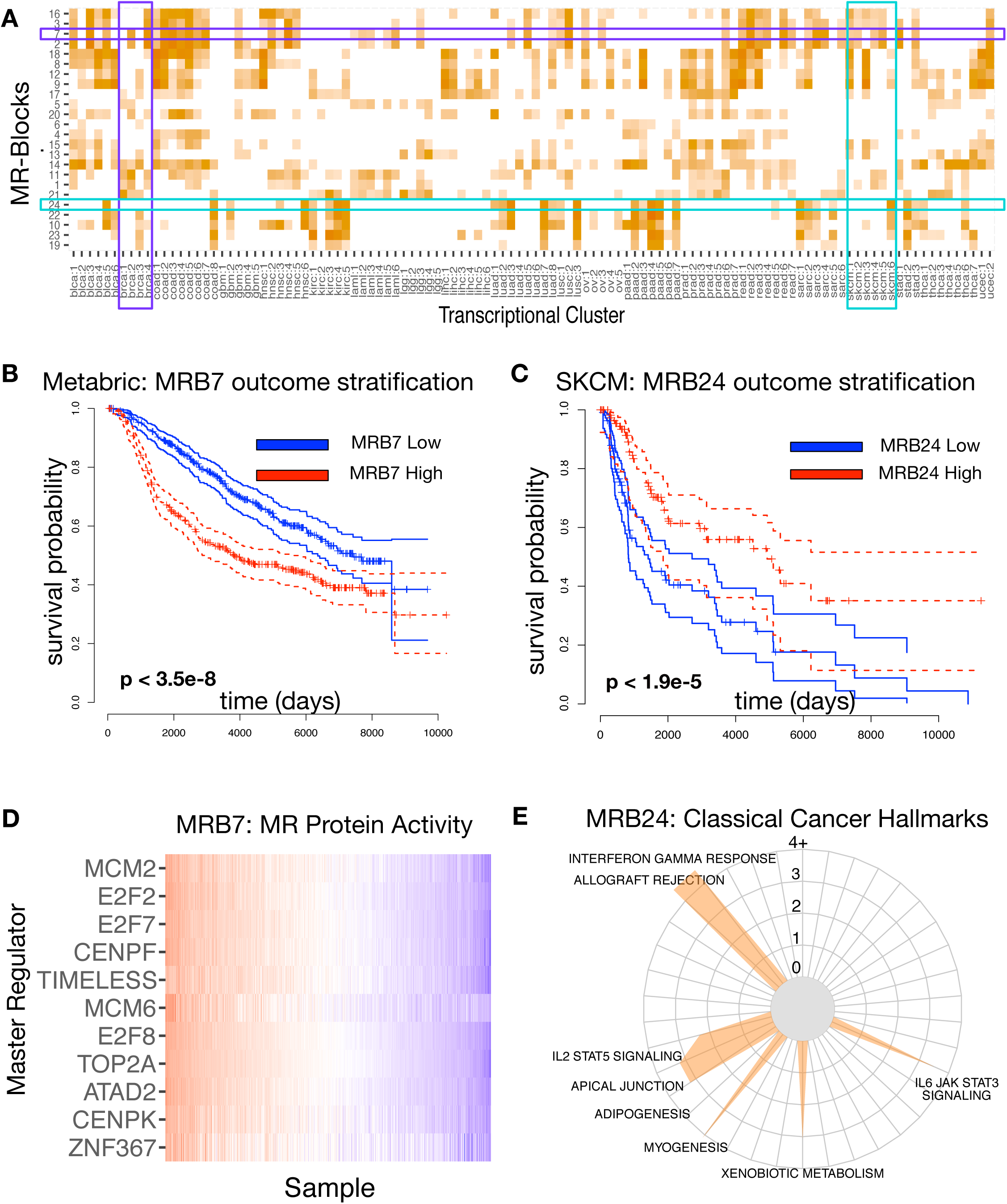
Checkpoint blocks are closely related to known cancer biology. (A) Active checkpoint blocks (y-axis) are shown in a heatmap summary across identified transcriptional subtypes (x-axis). Darker shades indicate higher mean activity across a subtype, for a given checkpoint. Breast cancer (BRCA) and melanoma (SKCM) transcriptional sub-type columns are highlighted along with checkpoint block 3 and 12 (rows). (B) Activity of MRB7 significantly stratifies Metabric breast cancer patients by outcome (p < 3.5E-8; Kaplan-Meier estimator). (C) Activity of MRB24 significantly stratifies TCGA melanoma patients by outcome (p < 1.9E-5; Kaplan-Meier estimator). In this case, high checkpoint activity leads to better outcome (D) VIPER Predicted activity for checkpoint proteins in checkpoint block 3 across Metabric breast cancer samples (columns). VIPER activity is highly correlated in all but one (E2F1) protein. (E) Enrichment “radar” plot for MRB24. Several hallmarks of cancer, including inflammatory/immune response hallmarks, apical junction IL/JAK/STAT transition are enriched within the checkpoint proteins of this block.

Thus, each Tumor Checkpoint comprises and is defined by a set of aberrantly-activated and/or inactivated MR-Blocks. This suggests that each MR-Block may regulate a set of complementary genetic programs required to implement and maintain a tumor cell’s transcriptional identity. Consistent with this hypothesis, we found highly significant enrichment of Cancer Hallmarks—as defined by the Broad Institute collection (Drake et al., 2016; Liberzon et al., 2015)—in MR-Block-specific MRs, with most hallmarks enriched in the MRs of at least one MR-Block (Figure S5D, Supplemental Data 7; see methods *Checkpoint Generation*). Confirming specificity, most MR-Blocks were enriched in only a handful of hallmarks (*N ≤ 5 for >50% of MR-Blocks*). In terms of clinical applicability, most hallmark blocks were able to significantly stratify patients by outcome, see Figure 5B and 5C, for BRCA stratification in the Metabric cohort using MRB2—an MR-Block comprised of classic cell growth, DNA repair, and cell division regulators (Figure 5D)—and for SKCM stratification in TCGA using MRB24—an MR-Block highly enriched in immune-related hallmarks (Figure 5E). See also Figure S6A for a comprehensive analysis across all TCGA cohorts. These results represent an initial attempt to elucidate how specific cancer hallmarks may be mechanistically regulated in each tumor subtypes.

We then assessed whether MR-Blocks could effectively stratify tumor cohorts based on outcome. For this purpose, we used a sparse Lasso COX proportional hazards regression model (Tibshirani, 1997), using the mean MR-Block activity of each sample as a predictor. In most cases, survival separation was more statistically significant than using the entire tumor-checkpoint (Figure S6B vs. S2E, Supplemental Data 8). For instance, in melanoma (SKCM) we observed striking survival separation (*p* < 1.6E-7), using a 6 MR-Block model—including MRB10, associated with strong inflammatory/immune phenotype (Supplemental Data 7). In contrast, the best outcome separation by full Tumor Checkpoint analysis was much less significant (*p* = 9.4E-3). Similarly, in colorectal cancer (COAD), significant outcome separation was achieved using a 3 MR-Block model (*p <* 3.5E-3)—with MRB6 providing the greatest contribution. In contrast, differential outcome by Tumor Checkpoint analysis was not statistically significant in this tissue type (*p* = 0.07).

To assess whether the MR-Block landscape emerging from this analysis would generalize to non-TCGA cohorts, we assessed VIPER-inferred activity of breast cancer relevant MRs from a large compendium of breast cancer samples with considerable long-term survival data (Curtis et al., 2012a). Considering *N* = 7 MR-Blocks with high differential activity in the TCGA breast cancer cohort (MRB2, 3, 7, 11, 14, 16, and 21), all of them but MRB11 provided statistically significant survival stratification, with 5 of the 6 MR-Blocks in the *p* = 1.88E-8 to 9.13E-7 range (Bonferroni corrected), as well as highly correlated activity of MR-Block MRs (Figure S6C). This suggests that MR-Block proteins may play a key role in tumor outcome by regulating key cancer hallmark programs.

### Cell line-specific MOMA-inferred tumor checkpoints are enriched in experimentally validated tumor dependencies

We further assessed whether MRs in MR-Block associated with viability-related cancer hallmarks were enriched in essential proteins, based on existing pooled RNAi screen data from the Achilles Project (Cowley et al., 2014), see Figure S2D for a conceptual workflow. Specifically, we used VIPER to transform RNASeq profiles of all Cancer Cell Line Encyclopedia (CCLE) into protein activity profiles, then matched the average protein activity profile of each of the 24 MR-Blocks to a set of best-matched cell lines, by MR enrichment analysis. Finally, we assessed essentiality of the corresponding MR-Block MRs based on their Achilles’ Project score. As expected, the three MR-Blocks enriched for growth and proliferation-related hallmarks (G2M, E2F, etc.) (Figure S5D) had the highest ratio of essential MRs (MRB2: 50%; MRB7: 43.8%; MRB3: 30.4%), including proteins such as E2F1, E2F2, E2F7, TOP2A, PTTG1, FOXM1, MYBL2, UHRF1, DNMT3B, ZNF695, TCF19, RBL1, and ZNF367. Interestingly, however, we also found a large fraction of essential proteins in additional blocks, including MRB6 (31.3%; ZNF436, HES1, HOXB7, TP63, TRIM29, GRHL1, PBX4, IKZF2, RARG, IRX5, HHEX, RUNX2, STAT5A, HDAC1, HOXC6) and MRB14 (18.8%; GRHL2, OVOL1, ZBTB7B), for instance. Not surprisingly, we found no Achilles validated MR proteins in immune-related MR-Blocks (MRB10, 22, 23, and 24)—consistent with lack of *in vitro* immune function. However, we already addressed the pan-cancer role of these proteins and of their upstream mutations in regulating immunity and inflammation in a prior publication (Thorsson et al., 2018a). Overall, we found MOMA-inferred MR proteins to be significantly enriched in essential genes, compared with 10^6^ randomly chosen, identically sized regulatory protein sets, not included in any Tumor Checkpoint (*p* = 7.1E-6; Figure S2E).

### MRB2 canalizes the effect of driver mutations in MAP3K7, SORBS3, BCAR1, PTEN, and TP53

As discussed, MRB2 mechanistically regulates the transcriptional identity of several highly aggressive subtypes, including in UCEC, STAD, SKCM, SARC, READ, PRAD, PAAD, LUAD, LIHC, LGG, and KIRC. Moreover, FOXM1 and CENPF—two of its core MR proteins—rank 2^nd^ and 17^th^ as most recurrently inferred across all TCGA tumor samples. Consistently, an MRB2-based regularized COX regression model produced several of the largest regression coefficients for outcome stratification across all TCGA samples (Supplemental Data 8), and is one of the most significant and effective single-block predictors of outcome across the TCGA cohorts (Figure S6A).

We thus sought to investigate the specific mutational events upstream of this MR-Block that determine its aberrant activation. MOMA identified 7 molecularly-distinct prostate adenocarcinoma (PRAD) subtypes, with significant survival separation (*p* = 6E-3; Cox proportional hazard model; Figure 6A, Figure S6D) between subtype 6 and subtypes 1, 3 and 5, driven by checkpoint proteins that include a majority subset of MRB2 (Figure 6A). Interestingly and consistent with (Aytes et al., 2014a), outcome difference was most significantly driven by MRB2 MRs (Figure S6B, Supplemental Data 8), with the lowest and highest MRB2 activity associated with best (subtype 1, 3, and 5) and worst (subtype 6) survival, respectively. Further supporting MRB2 as a key molecular determinant of disease outcome, we also observed high enrichment of negative prognosis samples, based on Gleason score and biochemical recurrence (Figure 6B,C), in subtype 6. This subtype also had the worst survival outcome of any cluster, which was significantly worse compared to cluster 3, the best outcome subtype, with 0 of 109 deaths (*p* < 7E-4; Figure S2B). To further study this malignant phenotype, we computed the differential gene expression signature between subtypes 6 and 1 and confirmed its highly significant enrichment in “G2M” (*p =* 1.6E-24), “E2F-Targets” (*p =* 1.8E-31), “Mitotic Spindle” (*p =* 2.6E-5), and “DNA Repair” (*p =* 2.2E-5) hallmarks (Figure 6D), which is consistent with the hallmark enrichment analysis of the proteins in MRB2 (Figure 6E).

**Figure 6:**
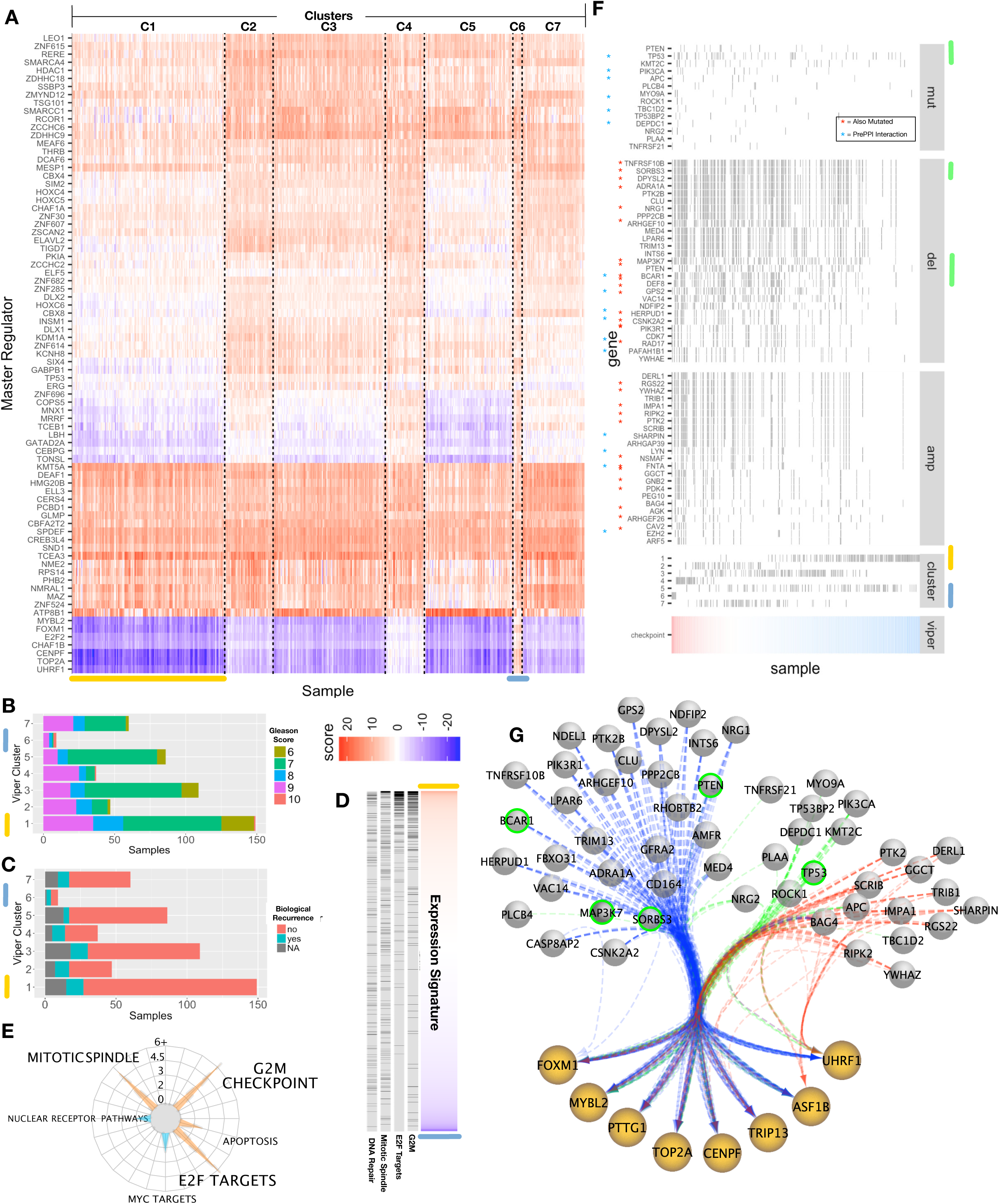
Proliferative checkpoint block 2 and associated causal genomic events drive the aggressive subtype of prostate cancer. (A) Heatmap of VIPER protein activity scores for the checkpoint proteins in all TCGA prostate cancer subtypes. Dashed vertical lines indicate subtype demarcation, rows are checkpoint proteins. Color indicates the VIPER inferred protein activity (red is high activity; blue is negative activity). (B) Clinical Gleason scores for TCGA prostate samples, grouped by the 7 clusters identified in our analysis (Figure 6A). All but one of the grade 10 samples were found in cluster 6, with the remaining sample found in cluster 4, which we found to have moderate activity in checkpoint block 2. (C) Clinical recurrence status of TCGA prostate samples, grouped by the 7 clusters in Figure 6A. Missing values are shown in grey, whereas recurrent samples are shown in blue/green. The largest fraction of recurrent samples was shown in cluster 6, with the second largest fraction in cluster 4. (D) Gene expression signature between 9 samples in cluster 6 and 149 samples in cluster 1 is sorted from highest differential gene expression (t-test on variance stabilized gene expression; red) to lowest (blue). Genes in significantly enriched respective hallmarks (GSEA; “DNA Repair”, NES = 2.6, *p* = 2.2E-16; “Mitotic Spindle,” NES = 3, *p* = 2.2E-16; “E2F Targets,” NES = 6.3, *p* = 2.2E-16; “G2M,” NES = 6, *p* = 2.2E-16) are shown as grey ticks. (E) Hallmark enrichment wheel of checkpoint block 2 proteins, from MSigDB 2.0 hallmark categories. Orange radii indicate enrichments that are statistically significant after multi-hypothesis correction (Benjamini-Hochberg FDR < 0.01). (F) Sample copy number and mutation events statistically associated with checkpoint 2 activity. Samples (columns) are sorted by checkpoint 2 VIPER activity (bottom); grey ticks indicate samples with a SNV/copy number/fusion event. Copy number events that are also mutated in one or more samples in the cohort are marked with a red star to their left. Genes are ordered from most (top) to least frequently altered in the cohort. The five genes selected for experimental validation are highlighted in green. (G) Network diagram of checkpoint 2 proteins and DIGGIT interactions highlighted in (F), with deletions (blue), mutations (green) and amplification events (red) shown as bundled edges. Green-circled events were selected for experimental follow-up (Figure 7).

We then considered the repertoire of genetic alterations identified by DIGGIT as upstream of MRB2, ranking them based on their combined statistical significance in the PRAD cohort, as well as across all pancancer cohorts. To visualize the genomic events with the strongest overall checkpoint association, we combined the strongest individual interactions of each of the eight MRs in MRB2 with equal weights (Figure 6F,G; Supplemental Data 8). Critically, we found that most of these genomic events would have been missed by existing mutation assessment algorithms (Table 4, Supplemental Data 9). When ranking all samples by the overall activity of MR-Block:2, clusters 6 was the one most enriched in samples with high activity, while cluster 1 was the most enriched in samples with low-activity.

**Table 4:** Putative mutational drivers in PRAD cohort. Mutational drivers upstream of MRB2, detection status for the MutSig2.CV algorithm, Clinical Correlation via the Broad TCGA Firehose pipeline, and Mutation-Assesor algorithms are shown in the respective columns.

| Node | MSIG.2CV | Clin.Correlate | Mut.Assesor |
| --- | --- | --- | --- |
| KMT2C | no | no | NA |
| TRIM13 | no | no | NA |
| PPP2CB | no | no | 0.2 |
| AGK | no | no | 1.1 |
| BAG4 | no | no | NA |
| ARL3 | no | no | NA |
| NDEL1 | no | no | NA |
| RAD17 | no | no | 1.375 |
| GGCT | no | no | 1.265 |
| TNFRSF10B | no | no | 1.545 |
| PTK2B | no | no | NA |
| EZH2 | no | no | 1.04 |
| TRIB1 | no | no | NA |
| MED4 | no | no | NA |
| PLCB4 | no | no | 1.525 |
| NSMAF | no | no | 1.7175 |
| CAV2 | no | no | NA |
| SORBS3 | no | no | 1.39 |
| MMS19 | no | no | NA |
| INTS6 | no | no | NA |
| YWHAZ | no | no | 3.735 |
| RGS22 | no | no | NA |
| GPS2 | no | no | 0.805 |
| ROCK1 | no | no | 0.715 |
| NDFIP2 | no | no | NA |
| PAFAH1B1 | no | no | NA |
| TBC1D2 | no | no | 1.8 |
| DEF8 | no | no | 1.84 |
| MAP3K7 | no | no | 2.195 |
| ARF5 | no | no | NA |
| VAC14 | no | no | 1.5 |
| DERL1 | no | no | NA |
| DEPDC1 | no | no | 0.855 |
| PEG10 | no | no | NA |
| PIK3R1 | no | no | 3.015 |
| NRG1 | no | no | 1.905 |
| LYN | no | no | 0.5625 |
| NRG2 | no | no | 1.645 |
| PTK2 | no | no | 0.375 |
| CASP8AP2 | no | no | NA |
| PDK4 | no | no | 0.9075 |
| ARHGEF10 | no | no | 0.9075 |
| DPYSL2 | no | no | 1.625 |
| FNTA | no | no | NA |
| GFRA2 | no | no | NA |
| HERPUD1 | no | no | 2.35 |
| GNB2 | no | no | 1.245 |
| IMPA1 | no | no | 3.51 |
| AMFR | no | no | NA |
| RHOBTB2 | no | no | NA |
| PIK3CA | yes | yes | 1.7625 |
| APC | no | yes | 1.1975 |
| CLU | no | no | NA |
| PLAA | no | no | 0.49 |
| TRIM23 | no | no | NA |
| BCAR1 | no | no | 1.655 |
| CSNK2A2 | no | no | 0.405 |
| ADRA1A | no | no | 0.895 |
| TP53 | yes | yes | 2.9925 |
| TP53BP2 | no | no | 2.06 |
| ERBIN | no | no | NA |
| BTRC | no | no | 1.6175 |
| CD164 | no | no | 1.1 |
| SCRIB | no | no | 0.345 |
| PTEN | yes | yes | 3.435 |
| RIPK2 | no | no | 0.345 |
| ADGRA2 | no | no | NA |
| TNFRSF21 | no | no | 1.325 |
| FBXO31 | no | no | NA |
| MYO9A | no | no | 1.83 |
| CDK7 | no | no | 1.775 |
| ARHGAP39 | no | no | 0.69 |
| YWHAE | no | no | NA |
| LPAR6 | no | no | NA |
| SHARPIN | no | no | NA |
| ARHGEF26 | no | no | NA |

We selected 6 DIGGIT-inferred loss-of-function events for experimental validation, including TP53^Mut^ (strongest pancancer association with MRB2 among single-point mutations), PTEN^Del^ (strongest pancancer association among deletions, also associated with PTEN^Mut^), MAP3K7^Del^ (strongest PRAD-specific association, among deletions), SORBS3^Del^ (one of the most significant associations, both pancancer and PRAD-specific, among deletions) (Figure 6B) and BCAR1^Del^, the strongest pancancer association, among deletions supported by a direct protein-protein interaction with one of the MRs (i.e., FOXM1)). These are visualized as green circles in the context of other statistically significant deletion (blue lines), mutation (green lines) and amplification (red lines) events in Figure 6D.

For experimental validation, 22Rv1 human prostate cancer cells were chosen, which present low MRB2 activity—thus providing an ideal model to detect MRB2 activity increase, following loss-of-function assays for the selected genes. Pools of 5 shRNAs/target were used to individually silence PTEN, TP53, MAP3K7, SORBS3 and BCAR1. Functional and tumorigenic effects were subsequently assessed both in vitro and in vivo (Figure 7A). VIPER analysis of gene expression profiles, following shRNA-mediated silencing of each candidate gene, confirmed significant increase MRB2 MR activity (Figure 7B). In addition, of the 5 candidates, MAP3K7, PTEN and TP53 showed the most pronounced and significant increase in cell migration using scratch assays at the indicated time points and relative to control cells infected with scramble shRNAs (Figure 7C-D), as further confirmed by Boyden chamber migration assays (Figure 7E). Finally, control and shRNA-silenced 22rv1 cells for each individual gene were engrafted in immune deficient mice to assess the relative capacity for tumor growth *in vivo.* As shown by these *in vivo* assays—and even discarding the expected effect on tumorigenesis associated with loss of PTEN and TP53—MAP3K7 silencing resulted in a marked and significant increase in tumor growth (p <0.01) (Figure 7F). As a result, all of the predicted loss of function events induced activation of MRB2 MR proteins, while three out of five had additional significant effects in terms of increased *in vitro* migration and *in vivo* tumorigenesis. Several of the phenotypes associated with MRB2, such as increased metastatic progression or reduced immunosurveillance, cannot be fully assessed in these assays or may require additional co-segregating events to be fully revealed.

**Figure 7:**
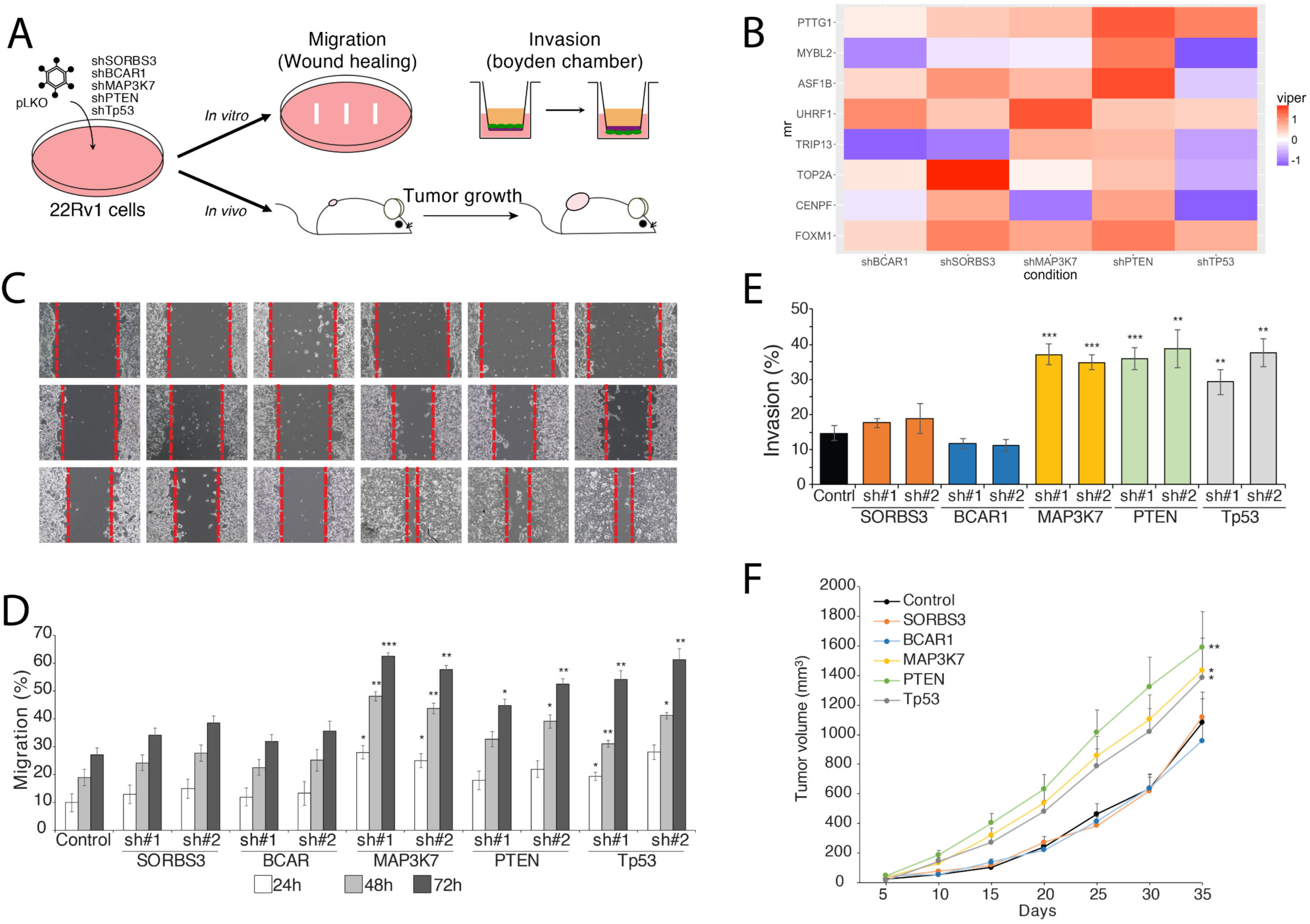
Functional validation of predicted candidates. (A) Schematics for functional assays. Androgen independent 22rv1 prostate cancer cells were infected with lentiviral control vectors and vectors containing shRNAs to silence recurrent genomic events upstream of FOXM1, namely, SORBS3, BCAR1, MAP3K7, PTEN and TP53. After selection for stable silencing, cells were used in *in vitro* in *in vivo* assays. (B) VIPER analysis of the MR-Block proteins (y-axis) in each silencing condition (x-axis). Red indicate increased activity relative to the control condition, blue decreased activity. *(C)* Migration of 22Rv1 cells was assessed in wound healing assays at 24 and 48 hours after scratching a confluent culture of control and silenced 22rv1. Quantification of the migration **assay is shown in** (D). Invasion capabilities of control and individually silenced 22rv1 cells was carried out in Boyden chamber invasion assays and quantification of the percent of invading cells is shown in *(E). (F)* Functional validation *in vivo.* Control and individually silenced 22rv1 cells where engrafted in mice and tumor growth was monitored over time until euthanasia. Tumor growth curves are shown. All *in vitro* experiments were done in triplicate in two independent replicates and significance was assessed using standard T-Student test comparing silencing to control. The *in vivo* assays where performed in two independent replicates and significance assessed using the Two-way analysis of variance (ANOVA). The p-values are indicated by * < 0.05, ** < 0.001 and *** < 0.0001.

## Discussion

The repertoire of transcriptional identities accessible to a cancer cell in response to endogenous and exogenous perturbations (i.e., its plasticity), is constrained by the cell’s genetic alteration landscape and by the baseline epigenetics of its tissue of origin. Yet, the specific mechanistic rationale of these constraints is largely unexplored. For instance, it is unclear why EGFR or NF1 mutations may alter the probability of individual GBM cells adopting a proneural or a mesenchymal identity. In this manuscript, we attempt to address this challenge by identifying Master Regulator proteins that mechanistically regulate the tumor cell’s transcriptional identity by integrating the effect of multiple genomic alterations in their upstream pathways. To achieve this goal both systematically and in a completely unbiased fashion, we analyzed 9,738 individual samples, representing the 20 largest TCGA cohorts using a novel methodology (MOMA).

MOMA revealed a highly modular regulatory architecture where 112 distinct tumor subtypes— representing distinct transcriptional identities—are implemented by combinations of only 24 regulatory modules comprised of Master Regulator proteins (MR-Blocks). Furthermore, an average of 33 Master Regulator proteins per subtype was sufficient to account for the effect of a majority of genomic alterations identified on a sample by sample basis, suggesting the existence of cross-tumor commonalities yielding a relatively small and yet highly universal repertoire of non-oncogene dependency mechanisms. Thus, by connecting MR proteins to genomic alterations in their upstream pathways, MOMA produced a comprehensive map of interactions between cancer’s genomic landscape and the MR proteins presiding over the transcriptional identity of distinct tumor subtypes. The fact that a large number of genomic events were found upstream of actual but not randomly selected MR proteins suggests that many more alterations than previously appreciated may be required to make a cancer cell. While some “passenger” genomic events may have been erroneously identified as MR modulators (false positives), we also expect that a large number of weakly-additive events may cooperate to provide a potentially large contribution to tumorigenesis, as is the case for other complex diseases (Boyle et al., 2017).

If further confirmed, these findings would have several relevant consequences for the study of cancer: **First**, they reduce the complexity arising from the extraordinary diversity of mutational patterns detected in cancer cells—even within the same mass—by providing a small number of highly universal modular regulatory dependencies, as well as the specific Master Regulator proteins comprising them. This was independently validated by assessing statistically significant overlap of MR proteins in proliferation related MR-Blocks with Achilles’ project dependencies, suggesting that MR-Blocks associated with other hallmarks (e.g., immunoevasion or migration) may be critical to tumor survival and progression *in vivo*. **Second**, they may redirect the search for new cancer drugs development, from the development of inhibitors of signaling proteins that only indirectly affect MR activity and whose effect can be easily bypassed by alternative mutations, to direct MR protein activity inhibitors inducing Tumor Checkpoint collapse, which was shown to abrogate tumor viability *in vivo*, see for instance (Alvarez et al., 2018; Califano and Alvarez, 2017). This is especially relevant because, over the last decade, regulatory proteins are relinquishing their status as undruggable targets, for instance as a result of novel covalent inhibitors targeting protein cystines (Singh et al., 2011) or via activation of degron mechanisms (Gan et al., 2015). **Third**, these findings dramatically expand the number of genetic alterations mechanistically linked to specific tumor subtypes. This stems from abandoning a purely associative, statistical methodology in favor of one that leverages the tumor-specific transcriptional-regulation and signal-transduction architecture to limit the number of genomic events inducing aberrant activity of Master Regulator proteins. **Fourth**, they represent a much finer-grain tumor-subtype molecular characterization, whose novelty and potential value is also supported by statistically significant association with patient outcome across every analyzed TCGA cohort. **Finally**, as previously shown for regulation of programs presiding over immune infiltration and immunoevasion (Thorsson et al., 2018c), the analysis provides direct mechanistic hypotheses for the specific proteins that regulate virtually each classic tumor hallmark, in different tumor subtypes, as well as for the specific genomic events that determine their aberrant activity.

Over the last 50 years, a number of cancer hallmarks, representing programs necessary for cancer cell survival and proliferation, have emerged (Hanahan and Weinberg, 2011), thus spurring research aimed at identifying the specific proteins and protein-modules that comprise them. This has led to development of several methods to ‘decompose’ the 20,000+ dimensional gene-expression data space into orthogonal programs, either using 2-dimentional matrices (Kim et al., 2017) or higher dimensional tensors (Sankaranarayanan et al., 2015), thus creating a simplified representation of the underlying cellular states and shared oncogenic alterations (Kim et al., 2017; Malta et al., 2018). These studies are encouraging and confirm that cancer hallmarks may be indeed implemented by coordinated activity of specific gene modules. However, the high complexity of these solutions combined with lack of direct biological interpretability continue to be critical roadblocks in terms of reducing these models to a set of hypotheses that may be experimentally validated. In addition, since these models arise from application of “non-convex” optimization problems, their stability and reproducibility are a concern, as multiple (and arbitrarily selected) sub-optimal solutions may exist. In contrast, we have shown that due to the use of large regulons, VIPER-based protein activity measurements are extremely reproducible, robust, and highly conserved within tumor subtypes (Alvarez et al., 2016; Califano and Alvarez, 2017). Indeed, based on their reproducibility, two VIPER-based algorithms (OncoTarget and OncoTreat (Alvarez et al., 2018)) have achieved NY State CLIA certification.

As compared to these other models, MOMA analysis produced 112 distinct Tumor Checkpoints, each comprising an average of only 33 proteins, which account for the effect of dozens of genomic alterations in their upstream pathways. More critically, each Tumor Checkpoint was shown to result from the superposition of only 24 independent, pancancer MR-Blocks, each implementing critical tumor hallmark functions. This modular organization yields biologically interesting findings, linking tumor hallmarks with their candidate mechanistic determinants and generating straightforward hypotheses that can be efficiently validated, as shown for mutations upstream of the MRB2 MR-Block in prostate cancer.

MRB2 was specifically selected for validation because it emerged as the most stable and robust pancancer MR-Block across all tumor subtypes, for clustering solutions ranging from *k* = 2 to *k* = 100 (Figure S5E). This MR-Block comprises 14 regulators of cell growth, DNA repair, and cell division, including: CENPK, HELLS, E2F2/7, MCM6, TIMELESS, TOP2A, PTTG1, FOXM1, MYBL2, ASF1B, CENPF, TRIP13, UHRF1 (Supplemental Data 6). Among these, FOXM1 and CENPF were previously validated as synergistic MRs of the most aggressive subtype of prostate cancer (Aytes et al., 2014a). However, their effect in regulating aggressive cancer across several distinct tumor cohorts could not have emerged without a systematic, pancancer study. TRIP13 is also known to play a critical role in chromosomal structure maintenance during meiosis (Roig et al., 2010), facilitated by the DNA topoisomerase 2-alpha subunit, TOP2A, which enables chromosome condensation and chromatid separation, and already represents a key cancer therapeutic target (Jain et al., 2013). FOXM1, CENPF, MYBL2, and TRIP13 have all been implicated as part of a core “proliferation cluster,” associated with poor outcome, whose activity is dependent on p53 inactivation (Brosh and Rotter, 2010). Indeed, MOMA identified mutations in TP53 as the most significant event upstream of aberrant FOXM1 and CENPF activation. UHRF1, also a candidate therapeutic target, is overexpressed in many cancers (Unoki et al., 2009), where it regulates gene expression and peaks in G1 phase, continuing through G2 and M, while ASF1B—a core member of the histone chaperone proteins responsible for providing a constant supply of histones at the site of nucleosome assembly—plays an essential role in many cancers and is predictive of outcome in some (Corpet et al., 2011). In addition, MRB2 comprises multiple proliferation-related proteins, such as E2F2, E2F7, and TIMELESS, and is associated with proliferative cancer hallmarks, including “E2F Targets” (*p* = 4.26E-09), “Mitotic Spindle” (*p* = 4.65E-07), “G2/M Checkpoint” (*p* = 5.96E-06), and “peroxisome” (*p* = 3.64E-04). Consistently, the proteins harboring the 100 most statistically significant recurrent genomic alterations upstream of MBR2 MRs were also enriched in these hallmarks—e.g., “E2F Targets” (*p* = 2.2E-03) and “Mitotic Spindle” (*p* = 5.7E-5). Thus, while the individual role of these proteins may be established in some cancer context, our study suggests that their ability to form a hyper-connected, synergistic core “subunit” represents a universal determinant of highly aggressive cancer subtypes, from melanoma and glioblastoma, to colorectal, prostate, and ovarian adenocarcinoma (Figure 5D). Not surprisingly, whenever MRB2 was predictive of survival, we found negative regression coefficients in the respective COX proportional hazards models, meaning higher MRB2 activity was predictive of worse survival (Figure S6B; ucec, stad, skcm, sarc, paad, luad, lihc, lgg, kirc).

Experimental validation of the 5 top recurrent mutations upstream of MRB2 not only confirmed its predicted functional properties but also showed that activity of MRB2 MRs was dysregulated following shRNA-mediated silencing of the five mutated genes. Interestingly, activity of MRB3 and MRB7 was correlated with MRB2 activity. These MR-blocks control complementary, yet distinct aspects of the proliferation hallmark, via established proliferative MRs such as E2F(1/2/7/8), as well as chromatin modification enzymes involved in mitotic progression (SUV39H1), assembly (CHAF1B), and mini-chromosome maintenance (MCM2/3/6/7).

At the other end of the functional spectrum, MRB24 emerged as significantly associated with inflammatory response and immune function, including via the immune-regulatory MR STAT1 (Figure 5B), with high activity in a subset of Cutaneous Melanoma (Figure 5C). Indeed, MRB24 activity was a highly-significant survival predictor in this tumor, based on Kaplan Meier analysis (Figure 5C), confirming that higher immune infiltration, may be associated with increased immunosurveillance and thus better outcome. MRB19 also emerged as highly enriched in the “immune activity” hallmark, including via (a) the MHC trans-activator CIITA, whose inactivation in cancer abrogates HLA-DR presentation thus promoting immunoevasion (Yazawa et al., 1999), (b) Cluster of Differentiation 86 (CD86), the canonical CTLA-4 ligand involved in immune checkpoint activation, as well as (c) several additional proteins—such as NOTCH4, MITF, etc.— commonly associated with an immunoevasive microenvironment, as reported in a recent analysis of master regulators of tumor immune response (Thorsson et al., 2018b) (Figure 5E).

Clearly—consistent with other large-scale, high-throughput analyses, both experimental and computational—one cannot expect all MOMA inferences to be correct. However, as shown in a large body of literature, experimental validation rates of the methodologies used by the MOMA framework—including ARACNe, VIPER, and DIGGIT—compare favorably with those of high-throughput experimental assays, see (Califano and Alvarez, 2017) for a comprehensive review. As a result, it is reasonable to assume that a significant subset or even the majority of these predictions will be eventually validated and will complement the existing knowledge on tumor subtype genetics and transcriptomics. In that sense, MOMA inferences represent high-likelihood hypotheses that may be further investigated by the research community to elucidate the mechanistic regulation of tumor hallmark programs across all cancers, including both the MR proteins that control these programs and the genetic alterations that determine their aberrant activity. It should also be noted that a number of significant improvements are possible and will be investigated in future work. For instance, ARACNe networks can be further improved by use of epigenetic data, such as that derived by ATAC and ChIP-Seq methodologies, while VIPER is being improved using results from systematic drug and CRISPRi perturbations.

To make the MOMA results available to the research community, as an interactive resource that can be easily queried and visualized, we have developed a publicly accessible graphical web interface that allows users to easily navigate the ∼2 million tumor-specific molecular interactions emerging from the MOMA analysis. Users can also execute advanced queries through this interface, using an efficient graph database based on Neo4j (neo4j.org).

## Acknowledgments

We acknowledge support from the genomic and small animal imaging facility, which are shared resources of the Herbert Irving Comprehensive Cancer Center at Columbia University, supported in part by NIH/NCI grant #P30 CA013696. This research was also supported by funding from the National Cancer Institute Outstanding Investigator Award to AC (CA197745), U54 Cancer Systems Biology Centers to AC and CAS (CA209997), R01 to CAS (CA173481 and CA196662), and by two S10 NIH Shared Instrumentation Grants (OD012351 and OD0217640). AA was supported by grant from the Spanish ISCIII-MINECO (PI16/01070, CP15/00090), EAURF/407003/XH, and a Fundación BBVA-Young Investigator Award.

## Author Contributions

Conceptualization and Methodology, E.O.P., A.A., F.M.G., M.J.A., C.AS. and A.C.; Investigation, E.O.P, A.A., F.M.G, S.J.J., M.J.A., and A.C.; Resources, E.O.P., B.C., S.J. and P.S.; Formal Analysis E.O.P., F.M.G., E.F.D., S.J. and M.J.A.; Writing – Original Draft, E.O.P, P.S., and A.C.; Writing – Review and Editing, all authors.

---

***Figure S1: Heatmap(s) of MOMA clustering each of the 20 TCGA subtypes.*** Checkpoint proteins for all subtypes are shown on the y-axis, samples on the x-axis. VIPER protein activity scores are plotted (red = high activity; blue = low activity) with the scale bar shown on the right. Established subtype identities are shown for select tissues, where available (BRCA, COAD, GBM, STAD).

***Figure S2: Functional validation of MOMA subtypes and survival segregation.*** (A) Similarity plot between MOMA identified sample clusters (bottom) and classical breast cancer subtypes (upper). Classical breast cancer subtypes are shown (light blue: luminal A; dark blue: luminal B; basal: red; her2: yellow). (B) Kaplan-Meyer survival plot, displaying differential outcome for the best and worst surviving subtype of each of the 20 TCGA Tissue types, with survival time in days plotted on the x-asis, and survival probability plotted on the y-axis. P-values for the COX proportional hazard model test between subtypes are displayed above each plot. Legends display the subtype identities (C) VIPER inferred protein activity heatmap for STAT3, CEBPD and CEBPB in Glioblastoma MOMA clusters 2 and 3. The black vertical line separates samples from subtype 2 (left) and subtype 3 (right). VIPER activities are colored by score (red=high; blue=low). (D) Illustration of how Achilles single gene essentiality screens are used in conjunction with patient samples and cell line models. Patient sample clusters are matched to the nearest cell line models by comparison with VIPER inferred protein activity profiles. Achilles K.O. scores for those specific cell lines are then used to assess single gene essentiality (E) Density plot of the number of Master Regulators identified as significantly essential in Achilles (Bonferroni corrected p-value < 1e-5) for each sample clusters checkpoint, as compared with randomly selected cMR checkpoints (black distribution; p < 1.6E-3) of the same size. The null model was constructed with 1E6 randomly selected checkpoints, and fitted to a normal distribution to asses statistical significance of the true number of significantly essential Master Regulators (153: blue vertical line).

***Figure S3: Checkpoint proteins are highly interconnected, downstream of known genomic drivers.*** (A) Significance of the enrichment for genomic drivers (CHASM: single point mutation events; GISTIC 2.0: focal copy number) upstream of predicted checkpoint proteins in each tissue of origin. Log10 p-values are shown in a bar plot, with the horizontal dashed line representing the canonical significance level of 0.05. (B) Enrichment ratios for genomic drivers upstream of predicted checkpoint proteins in each tissue type. The distribution of enrichment ratios is shown in each violin plot. (C) Density plots of the mean shortest path distance between all pairs of predicted checkpoint proteins in each tissue of origin (blue), compared with pairwise distances between random pairs of transcriptional and co-transcriptional proteins, in the HumanNet network. (D) Density plots of the mean shortest path distance between all pairs of predicted checkpoint proteins in each tissue of origin (blue), compared with pairwise distances between random pairs of transcriptional and co-transcriptional proteins, in the Multinet network. (E) Density plots of the mean shortest path distance between all pairs of predicted checkpoint proteins in each tissue of origin (blue), compared with pairwise distances between random pairs of transcriptional and co-transcriptional proteins, in the PrePPI protein-protein interaction network.

***Figure S4a-t: Genomic saturation and identity plots for each of the 112 identified subtypes, within 20 TCGA tissues of origin.*** Left column: genomic events are shown on the y-axis, with frequency of alteration in the respective cohort displayed on the x-axix; deletion events are shown as blue marks, amplifications are red, mutations green. All events are identified as interacting with the candidate Master Regulator (cMR) proteins that are selected via the genomic saturation analysis shown on the right column, in the respective subtype/tissue. Saturation curves on the right each correspond to a single sample cluster, with the quantity of cMRs used to explain genomic events on the x-axis and the average number of genomic events (and fraction of all non-silent SNV and GISTIC2.0 identified events) per-sample on the y-axis. The dashed line indicates the identified inflection point, and defines the cluster checkpoint as all cMR proteins to the left of that line. (D) No saturation detected above the null distribution for subtype (2) due to low mutational burden.

***Figure S5: Checkpoint block discovery and hallmarks of cancer enrichment.*** (A) Density plot of the number of different checkpoints (of the 112 identified pan-cancer subtypes) each cMR participates in (solid red line), with the fraction of all ∼2500 transcription factors (TF) and co-factors (coTF) considered shown on the y-axis, compared to a null model constructed by randomly placing (TF/coTF) proteins into bins the same size as the 112 checkpoints, permuted 100 times (dashed black line). The vertical dotted line represents cMRs that are found in four or more checkpoints. The real and null distributions are significantly different, according to a non-parameteric Kolmogorov–Smirnov test (p < 2.2E-16). (B) Plot of the analytical clustering score for k=2 to k=100 checkpoint clusters, for the 407 highly recurrent candidate Master Regulator (cMR) proteins across tissue types. The 24 cluster solution of checkpoint “blocks” was found to be the highest scoring (green line). (C) Relative score representing the specificity of enrichment in the classical hallmarks of cancer (y-axis) across all 24 checkpoint blocks, as the blocks are “expanded” with additional nearest neighbor cMRs (x-axis). The coverage score (blue line) represents the Eigen-trace of the covariance matrix of all hallmark enrichments for all checkpoint blocks, while the dashed black line is the delta with the previous expansion factor. We selected k=6 as it is both an early absolute maximum and has one of the highest rates of improvement over the previous score (k=5). (D) Hallmark enrichments that are significant after multiple-hypothesis correction (Benjamini-Hochberg FDR) for each of the final 24 checkpoint blocks. (E) Violin plots of the Jaccard concordance index of each of the 24 checkpoint blocks with the most similar cluster found in each of the other clustering solutions (k = 2 to 100, excluding 24). Sorted left to right, from most to least concurrent.

***Figure S6: Analysis of survival/outcome predictions from MR-Block activities.*** (A) The negative log p-values of single-variable cox regression models are shown for each MR block (columns), representing the ability of each MR-block to predict patient outcome, across each of 20 TCGA tissue types (rows). Bars represent the −log10(pvalue) significance of each predictor, truncated at (log10(p)=5) for visual clarity; values less than (log10(p)<1) are not shown. The dashed line represents the canonical statistical significance level of p=0.05. (B) Survival plots of all 20 TCGA cohorts using a regularized cox proportional hazards model trained on the mean activity of the 24 MR-blocks. P-values for the fitted cox regression models (coefficients) are shown above each plot. Censors are shown and ticks along each axis. (C) Analysis of Pan-cancer checkpoint block activity in the Metabric breast cancer dataset. VIPER activities of the 7 MR-Blocks that were found to be highly active in the TCGA breast cancer cohort (Figure 5C), Differential survival outcomes shown for Metabric samples with positive mean activity of the proteins in each checkpoint (red) and negative mean activity (blue). Some checkpoint proteins were not inferred by the ARACNe regulon generated from Metabric data, and are omitted from the heatmaps. Survival separation was most significant for blocks 2, 3, 7 and 16, as well as block 21 (p < 2E-8, p < 2E-8, P<3E-8, P<3E-8, respectively). In contrast, we found the separation with block 14 to be only marginally significant (p < 0.006), and non-significant in block 11 (p < 0.3). (D) Censored survival plot of the TCGA PRAD (prostate cancer) cohort subtypes 3 (best survival; n=109, deaths=0) and subtype 6 (worst survival; n=9, deaths=1). Separation is significant according to a cox proportional hazards model (p < 7E-4).

## STAR* Methods

### Pan-cancer protein inference

RNA-Seq raw gene counts were downloaded from the TCGA firehose (gdac.broadinstitute.org), transformed to RPKM using the average transcript length for each gene and log2 transformed. Transcriptome-wide expression signatures were computed by two non-parametric transformations. First, each column (tumor sample) was rank transformed and scaled between 0 and 1. Then each row (gene) was rank transformed and scaled between 0 and 1. Finally, the activity of ∼2,500 regulatory proteins was estimated by the VIPER algorithm, using tissue-matched ARACNE regulons (Giorgi et al., 2016; Lachmann et al., 2016).

### DIGGIT

We identified statistically associated SNP events with the DIGGIT algorithm. Instead of using the mutual-information computation outlined in the published DIGGIT method (Alvarez et al., 2015) we computed the aREA enrichment (Alvarez et al., 2016) of the sample set with non-silent coding mutations in a given gene, against the ranked protein-activity signature inferred by VIPER for a given MR. This was performed for each VIPER Inferred Protein (VIP) / mutated gene pair with at least 4 samples with a non-silent alteration. Similarly SNP6 copy number profiles were downloaded from the Broad Institute and we picked a threshold value of 0.5, the mean value that we found to be optimally sensitive for detection with DIGGIT while maintaining high specificity for functional events as explored in recent literature (Jerby-Arnon et al., 2014).

### DIGGIT Null Model

A null model was constructed specific to each TCGA tissue type by considering the 1253 VIPs with the lowest absolute mean activity as a ‘null set’; we then computed the empirical p-values and q-values of the each DIGGIT/aREA score against the distribution generated with aREA on the null set of VIPs using the ‘*q*-value’ Bioconductor package (3.5) (Kall et al., 2008). Positive DIGGIT/aREA z-scores with an uncorrected empirical p-value of less than 0.05 over the background were combined using Stouffer’s method to generate three separate rankings for each VIP (Jerby-Arnon et al., 2014) based on SNV mutations, amplification events, and deletion events, respectively. CINDy was run using gene expression and the computed VIPER profiles separately within each TCGA tissue type. For most tissue types the number of CINDy interactions between genes with genomic alterations and VIPs with significant DIGGIT scores was large—hundreds to tens of thousands—and only these interactions were retained when computing the SNV/Amplification/Deletion rankings detailed above. In the few cases where overlap was less than 100 total interactions, all significant DIGGIT interactions were retained and the CINDy data utilized at a later step. Fusion calls were detected with the PRADA algorithm (Torres-Garcia et al., 2014), aREA and null-model aREA scores were computed in the same way.

We used the PrePPI database 1.2.0 (Zhang et al., 2013) to incorporate structural information into the rankings. We first converted all high-confidence (probability > 0.5) PPI interactions into empirical p-values by ranking and binning the likelihood scores, and assigning the lowest bin the probability of interaction based on the count of all possible pairs within the PrePPI database. Significant DIGGIT interactions with corresponding PrePPI interactions were considered for each VIP; the PrePPI empirical p-values were combined using Fisher’s method to generate rankings for SNV, Amplification and Deletion based DIGGIT interactions, respectively.

### Integrated rankings

Integrated rankings were generated by first removing the conditional dependency of the DIGGIT-based score for each MR by conditioning it on the rank of the VIPER score, and then converting the rank to an empirical one-tailed p-value. Similarly, PrePPI scores were conditioned on the DIGGIT scores for each, as were CINDy scores for several tissue types with a small number of CINDy predictions (see above). This conditional model was applied separately for each of the SNV, Fusion, Amplification and Deletion data types; the p-values from all conditionally independent tests were combined using Fisher’s method to generate a single ranking of candidate MRs for each tissue.

### Survival analysis

Clinical data was downloaded from the Broad Institute GDAC website (gdac.broadinstitute.org). We used the ‘survival’ R/CRAN package version 2.41-3 to fit a Cox proportional hazards model to each sample grouping defined by the initial cluster. We then defined the “best” survival clusters as the one with the lowest proportion of observed to expected death events, and the “worst” survival as the highest observed/expected ratio. We then fit a second Cox model exclusively to samples from those two clusters and calculated the significance of survival differences between “best” and “worst” clusters in that model.

### Sample clustering

Each tissue-specific VIPER activity matrix was clustered using k-medoids clustering with k ranging from 2 to 10 clusters, using a distance matrix defined by the weighted Pearson correlation between sample VIPER profiles. Weights were defined by the negative log p-values of the integrated scores described above in *Integrated Rankings*; to increase the contribution of high scoring Master Regulators we also transformed the negative log p-values with a square operation before generating the distance matrix. A silhouette-like score was calculated for each sample at each *k* value, using the aREA function described in (Alvarez et al., 2016) to determine the enrichment in similarity between each sample and it’s assigned cluster. We then chose the *k* that maximized the mean score across all samples.

### GEX clustering

Each tissue-specific gene expression matrix was clustered using k-medoids clustering with *k* set as the same value chosen for the tissue-specific VIPER activity clustering (see methods, Sample clustering). Distance between samples was defined using Pearson correlation between gene expression profiles. Silhouette scores were computed using the ‘cluster’ package in R.

### Candidate drivers

Mutation and SNP6 copy-number data was downloaded from the Broad Firehose platform (gdac.broadinstitute.org), as described in (methods: clustering/DIGGIT). We downloaded analysis results from Firehose and characterized each SNP as a candidate driver event if it achieved a p-value of 0.05 (uncorrected) or less according to the CHASM algorithm. Similarly, focal copy number events were considered “candidate drivers” if they were considered a high confidence (99% interval) event according to the GISTIC2.0 algorithm.

### Genomic coverage

Genomic events considered “candidate drivers” (see above) were used in the sample-specific analysis if they had a sufficient number of events to be detected by the DIGGIT algorithm (4 events, in each TCGA tissue type).

### Checkpoint Generation

Proteins were clustered with the VIPER protein activity matrix on the gene level, using a Euclidean distance metric and partitioning around medoids (PAM) for a predefined set of clusters *k,* from 2 to 100. The cluster fit was defined as the mean cluster reliability of each proteins fit to its respective cluster, which is calculated as the aREA enrichment score of the cluster member set on the distance vector between the protein and all other protein in the matrix. We chose an optimal *k* of 24 as shown in Figure S5B. Each “core” cluster was expanded by the *n candidate Master Regulator* proteins with the best similarity (outside of the original cluster), for all *n* in the range of 0 to 100. For each *n* in this range, we computed the trace of the covariance matrix calculated from hallmark enrichment across the 20 checkpoints expanded by *n* to approximate the total variance across the space defined by hallmark enrichment. We found an optimal increase in this variance at an expansion number of 6 (Figure S5C) and defined the “fuzzy” checkpoints at that threshold.

### Hallmark Enrichment

Cancer Hallmarks were defined as the 25 gene sets defined by the Broad Institute and refined/simplified by others (Drake et al., 2016; Liberzon et al., 2015). We computed the p-value of the hypergeometric overlap between each hallmark gene set and each checkpoint, using the cardinality of all candidate MRs (2506) as the “universe” size.

### Achilles Essentiality Validation

Achilles shRNA DEMETER knockout scores were downloaded from The Broad Institute for all cell lines. Transcription Factor (TF’s) Achilles dependencies scores were re-normalized by fitting bimodal normal mixture models using the R package ‘mixtools’. The most positive (least dependent) sub-population was set as the reference distribution for the re-normalized “dependency score” as a z-score. By binning Achilles-scores into distinct sub-populations, this procedure assumes discrete transcriptional-states with resolvable effects on cell-viability. In the context of orthogonal transcriptional programs (e.g. basal vs luminal breast cancer) this bias should boost meaningful signal for causal transcription factors. In cases of more continuous relationships between TF dependency and viability (e.g., house-keeping programs) this bias would most likely destroy information.

For each of the 112 TCGA subtypes, we matched the centroid sample to all CCLE VIPER profiles, using the ‘viperSimilarity’ algorithm included with the VIPER algorithm (Alvarez et al., 2016), after weighting each patient-sample by the MOMA scores for the corresponding tissue. Cell lines that were significant matches (FWER < 0.01; Bonferroni correction) were compared with non-matches (*p* = 1) using a non-parametric rank-based Mann-Whitney-Wilcox test; significant FDRs after multiple hypothesis correction (Benjamini-Hochberg FDR < 0.05) were retained for each subtype.

### METABRIC Breast cancer analysis

We ran ARACNE with 100 bootstrap iterations and a M.I. threshold of 1e-8, separately for the candidate TF and coTF regulators. Protein activity was inferred across all samples, using the VIPER algorithm. Survival analysis was performed by first calculating the mean VIPER activity across checkpoint proteins and binning samples into “high” and “low” quantiles, for each checkpoint. Clinical data was downloaded from the Broad Institute GDAC website (gdac.broadinstitute.org). We used the ‘survival’ R/CRAN package version 2.41-3 to fit a Cox proportional hazards model to each sample grouping, using the last known follow-up date, and testing for significant survival differences with that model.

### Interaction rankings

CINDy interactions were converted to empirical p-values by ranking and binning the number of significant triplets (Giorgi et al., 2014), and assigning the lowest bin the probability of interaction based on the count of all modulator-TF interactions. For each modulator-TF interaction, the CINDy based p-value, PrePPI p-value, p-value based on the DIGGIT/aREA score and the p-value generated by the DIGGIT null model were integrated using Fisher’s method to create a single ‘integrated’ p-value. For each MR, we computed the Benjamini-Hochberg false discovery rate of all integrated p-values and removed those above a threshold of 10%. Integrated p-values were combined across tissue types using Fisher’s method to generate a pancancer ranking.

### Proliferation cluster interactions

For each of the candidate Master Regulator proteins in checkpoint 2 we computed the rankings based on the integrated p-value in prostate cancer, as well as the cross-pancancer rankings for the same interactions. We created a combined rank from these two lists, using an additive mean and retained the top 20 interactions for each MR. Interactions were visualized with the Cytoscape software package (Shannon et al., 2003).

### ARACNe and VIPER analysis of the Sboner dataset

Clinical data and gene expression microarray data for 281 prostate cancer samples was downloaded from the Gene Expression Omnibus (GEO) (ID GSE16560). The expression profiles for 6100 transcriptionally informative genes (Gene Expression Omnibus Platform GPL5474) was used to generate ARACNe networks for the same TF and co-TF definitions used for the TCGA analysis, respectively. VIPER scores were computed for 563 TFs and 254 co-TFs across all 281 samples; representative candidate Master Regulator in the Pan-cancer checkpoint 2, identified through our TCGA based analysis included TRIP13, TOP2A, PTTG1, MYBL2, FOXM1 and CENPF. We computed the mean VIPER activity across these candidate Master Regulators and selected the top and bottom quantiles of samples with highest and lowest mean activity, respectively, for further analysis.

### Perturbation dataset VIPER analysis

We generated a signature for count data from each experimental condition, using the control condition as a reference, and performing a t test, using 100 permutations of the samples (columns) as a null model. This signature and null model were inputted to the ‘msviper’ function in the VIPER Bioconductor package, along with the TCGA Prostate cancer regulon. A second null model was constructed by re-running this same analysis on 100 permutations of the column labels, and a t-test was performed between the VIPER scores from each condition and this null, to assess the overall ability in reverting the signature for checkpoint 2 proteins.

