## Supplemental Figures for "A Modular Master Regulator Landscape Determines the Impact of Genetic Alterations on the Transcriptional Identity of Cancer Cells"

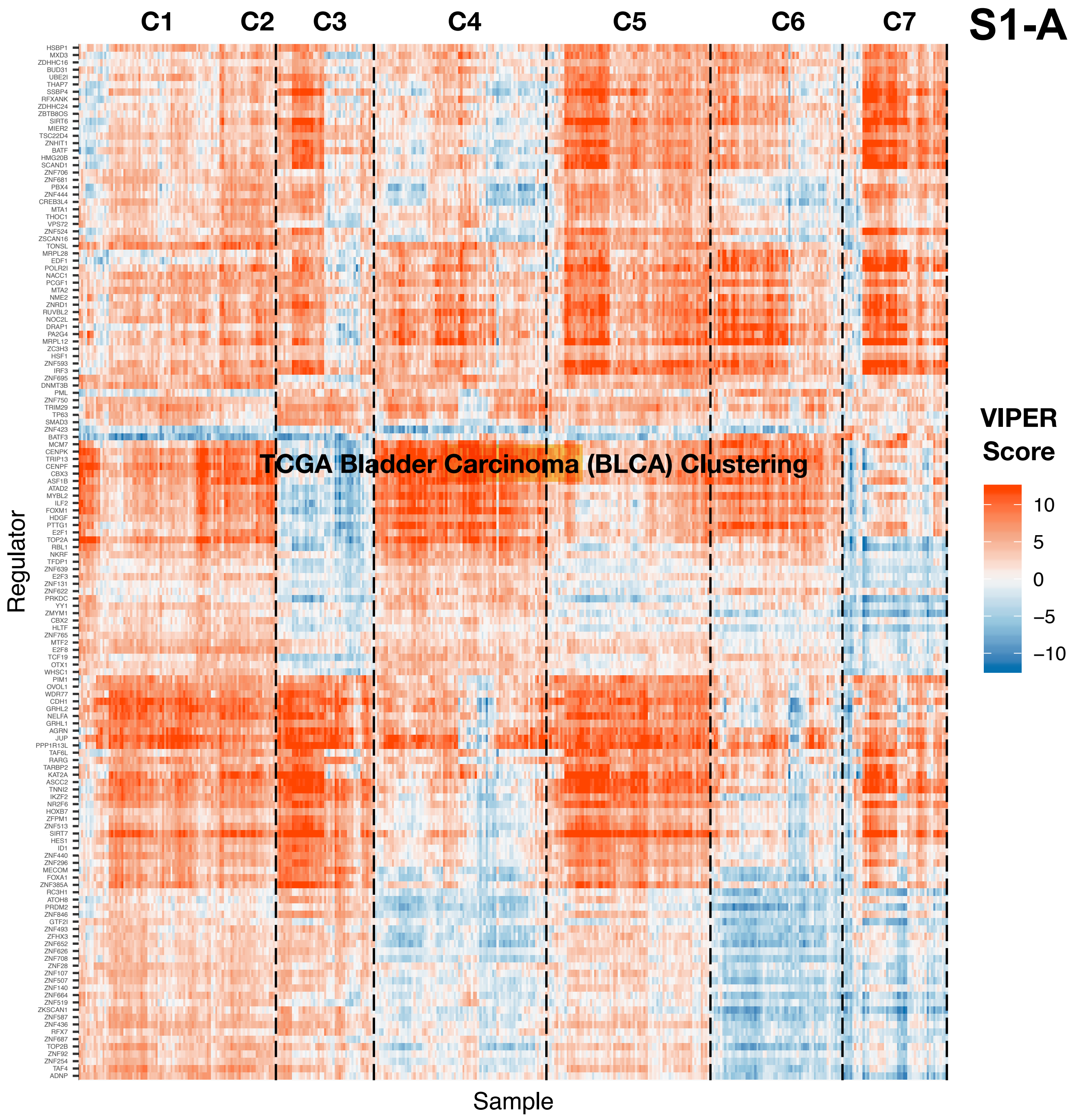
S1-A
C1
C2
C3
C4
C5
C6
C7
TCGA Bladder Carcinoma (BLCA) Clustering
Regulator
Sample
VIPER Score
10
5
0
−5
−10
HSBP1
MXD3
ZDHHC16
BUD31
UBE2I
THAP7
SSBP4
RFXANK
ZDHHC24
ZBTB8OS
SIRT6
MIER2
TSC22D4
ZNHIT1
BATF
HMG20B
SCAND1
ZNF706
ZNF681
PBX4
ZNF444
CREB3L4
MTA1
THOC1
VPS72
ZNF524
ZSCAN16
TONSL
MRPL28
EDF1
POLR2I
NACC1
PCGF1
MTA2
NME2
ZNRD1
RUVBL2
NOC2L
DRAP1
PA2G4
MRPL12
ZC3H3
HSF1
ZNF593
IRF3
ZNF695
DNMT3B
PML
ZNF750
TRIM29
TP63
SMAD3
ZNF423
BATF3
MCM7
CENPK
TRIP13
CENPF
CBX3
ASF1B
ATAD2
MYBL2
ILF2
FOXM1
HDGF
PTTG1
E2F1
TOP2A
RBL1
NKRF
TFDP1
ZNF639
E2F3
ZNF131
ZNF622
PRKDC
YY1
ZMYM1
CBX2
HLTF
ZNF765
MTF2
E2F8
TCF19
OTX1
WHSC1
PIM1
OVOL1
WDR77
CDH1
GRHL2
NELFA
GRHL1
AGRN
JUP
PPP1R13L
TAF6L
RARG
TARBP2
KAT2A
ASCC2
TNNI2
IKZF2
NR2F6
HOXB7
ZFPM1
ZNF513
SIRT7
HES1
ID1
ZNF440
ZNF296
MECOM
FOXA1
ZNF385A
RC3H1
ATOH8
PRDM2
ZNF846
GTF2I
ZNF493
ZFHX3
ZNF652
ZNF626
ZNF708
ZNF28
ZNF107
ZNF507
ZNF140
ZNF664
ZNF519
ZKSCAN1
ZNF587
ZNF436
RFX7
ZNF687
TOP2B
ZNF92
ZNF254
TAF4
ADNP

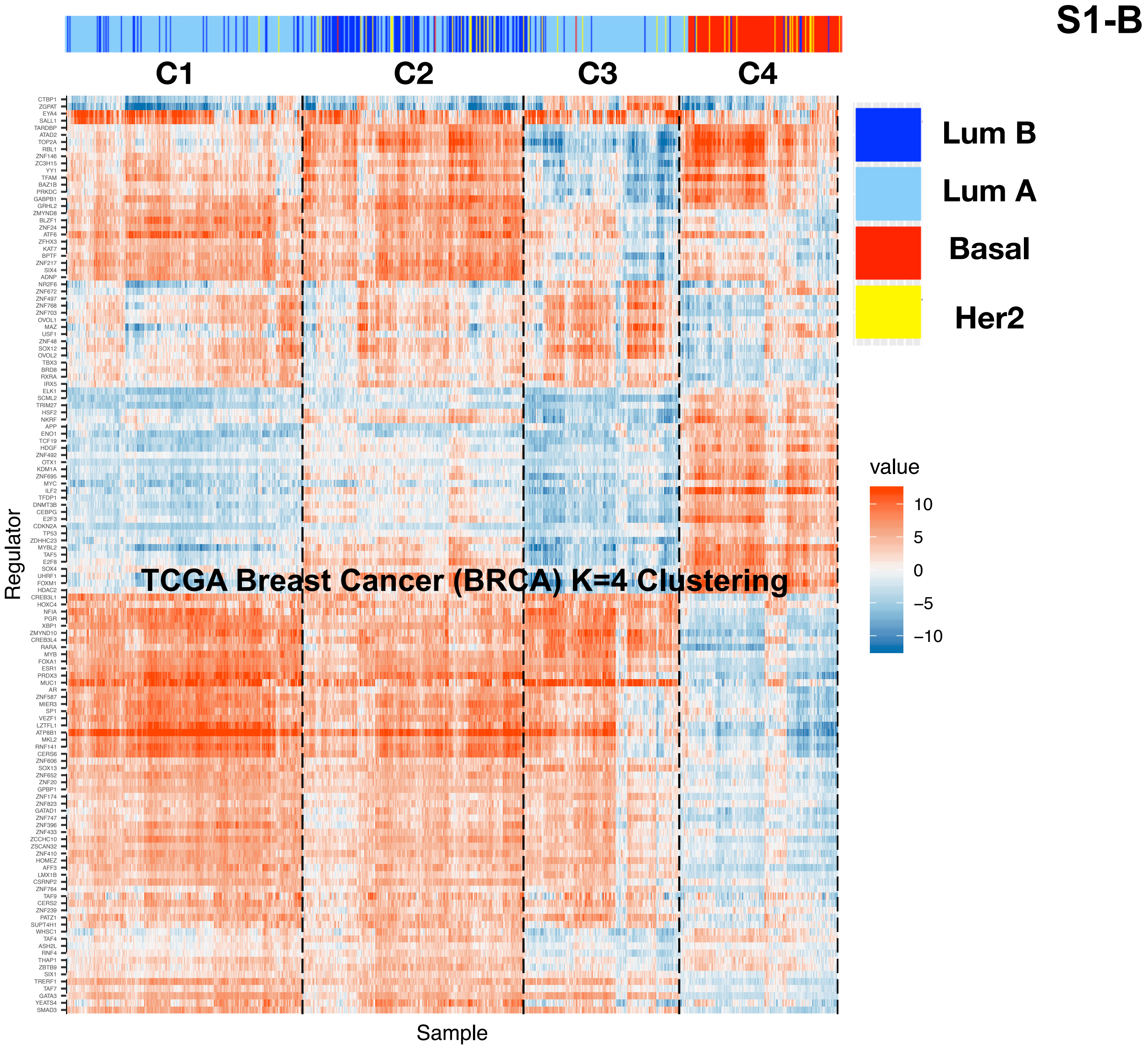

S1-B
C1
C2
C3
C4
Lum B
Lum A
Basal
Her2
value
10
5
0
−5
−10
TCGA Breast Cancer (BRCA) K=4 Clustering
Regulator
Sample

S1-C

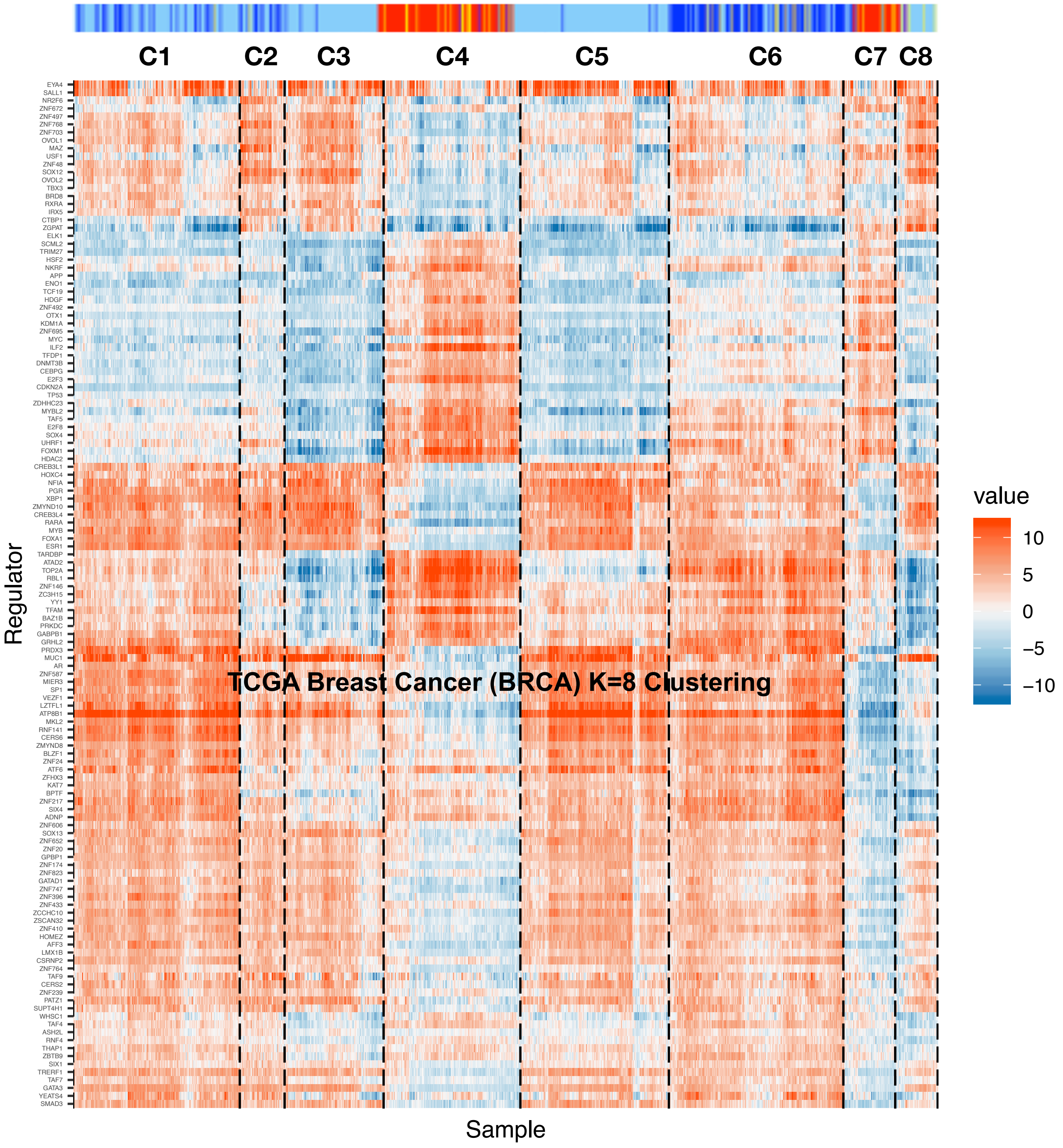

C1
C2
C3
C4
C5
C6
C7
C8
TCGA Breast Cancer (BRCA) K=8 Clustering
Regulator
Sample
value
10
5
0
−5
−10

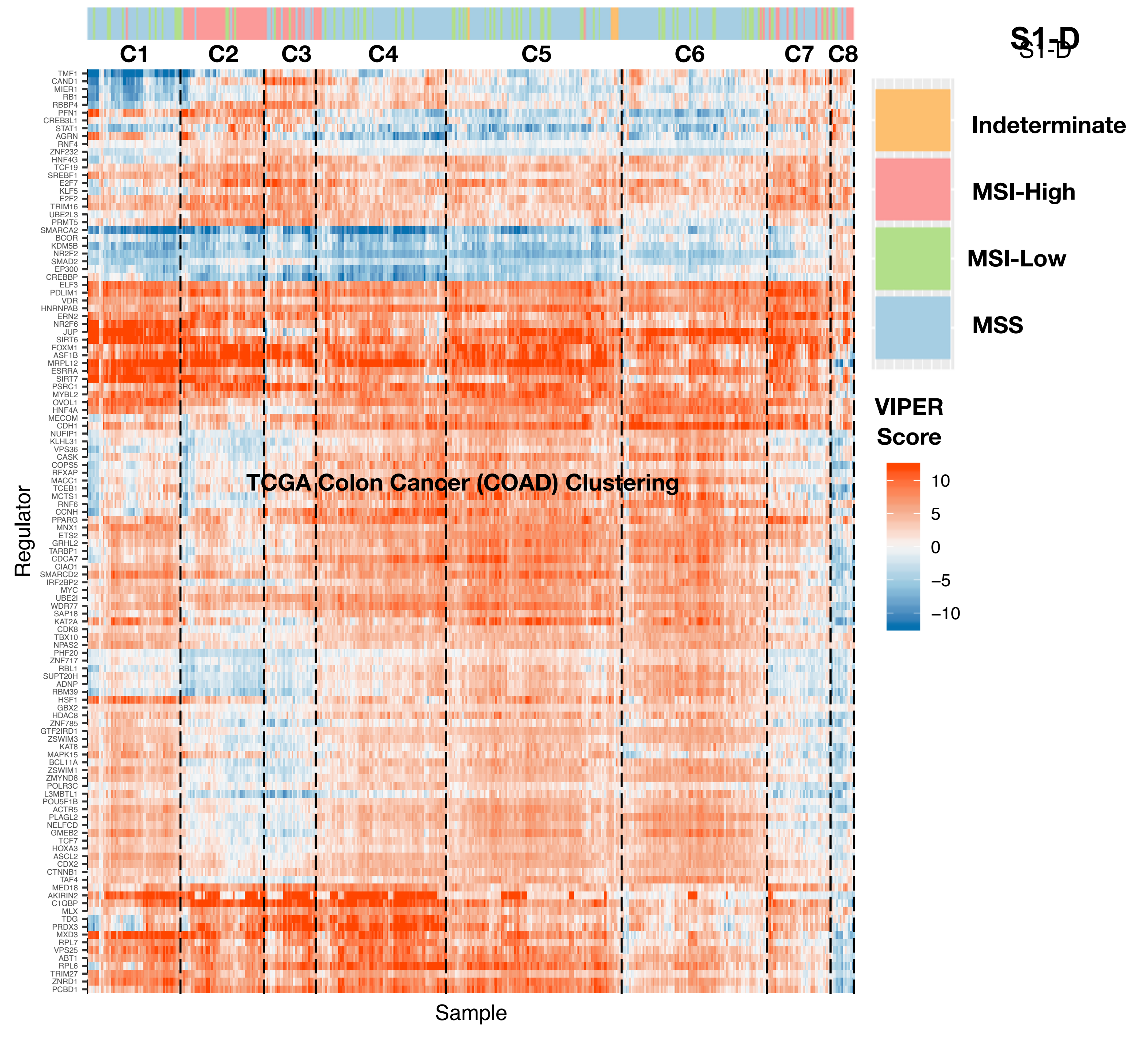
S1-D
C1
C2
C3
C4
C5
C6
C7
C8
TCGA Colon Cancer (COAD) Clustering
Regulator
Sample
TMF1
CAND1
MIER1
RB1
RBBP4
PFN1
CREB3L1
STAT1
AGRN
RNF4
ZNF232
HNF4G
TCF19
SREBF1
E2F7
KLF5
E2F2
TRIM16
UBE2L3
PRMT5
SMARCA2
BCOR
KDM5B
NR2F2
SMAD2
EP300
CREBBP
ELF3
PDLIM1
VDR
HNRNPAB
ERN2
NR2F6
JUP
SIRT6
FOXM1
ASF1B
MRPL12
ESRRA
SIRT7
PSRC1
MYBL2
OVOL1
HNF4A
MECOM
CDH1
NUFIP1
KLHL31
VPS36
CASK
COPS5
RFXAP
MACC1
TCEB1
MCTS1
RNF6
CCNH
PPARG
MNX1
ETS2
GRHL2
TARBP1
CDCA7
CIAO1
SMARCD2
IRF2BP2
MYC
UBE2I
WDR77
SAP18
KAT2A
CDK8
TBX10
NPAS2
PHF20
ZNF717
RBL1
SUPT20H
ADNP
RBM39
HSF1
GBX2
HDAC8
ZNF785
GTF2IRD1
ZSWIM3
KAT8
MAPK15
BCL11A
ZSWIM1
ZMYND8
POLR3C
L3MBTL1
POU5F1B
ACTR5
PLAGL2
NELFCD
GMEB2
TCF7
HOXA3
ASCL2
CDX2
CTNNB1
TAF4
MED18
AKIRIN2
C1QBP
MLX
TDG
PRDX3
MXD3
RPL7
VPS25
ABT1
RPL6
TRIM27
ZNRD1
PCBD1
Indeterminate
MSI-High
MSI-Low
MSS
VIPER Score
10
5
0
−5
−10

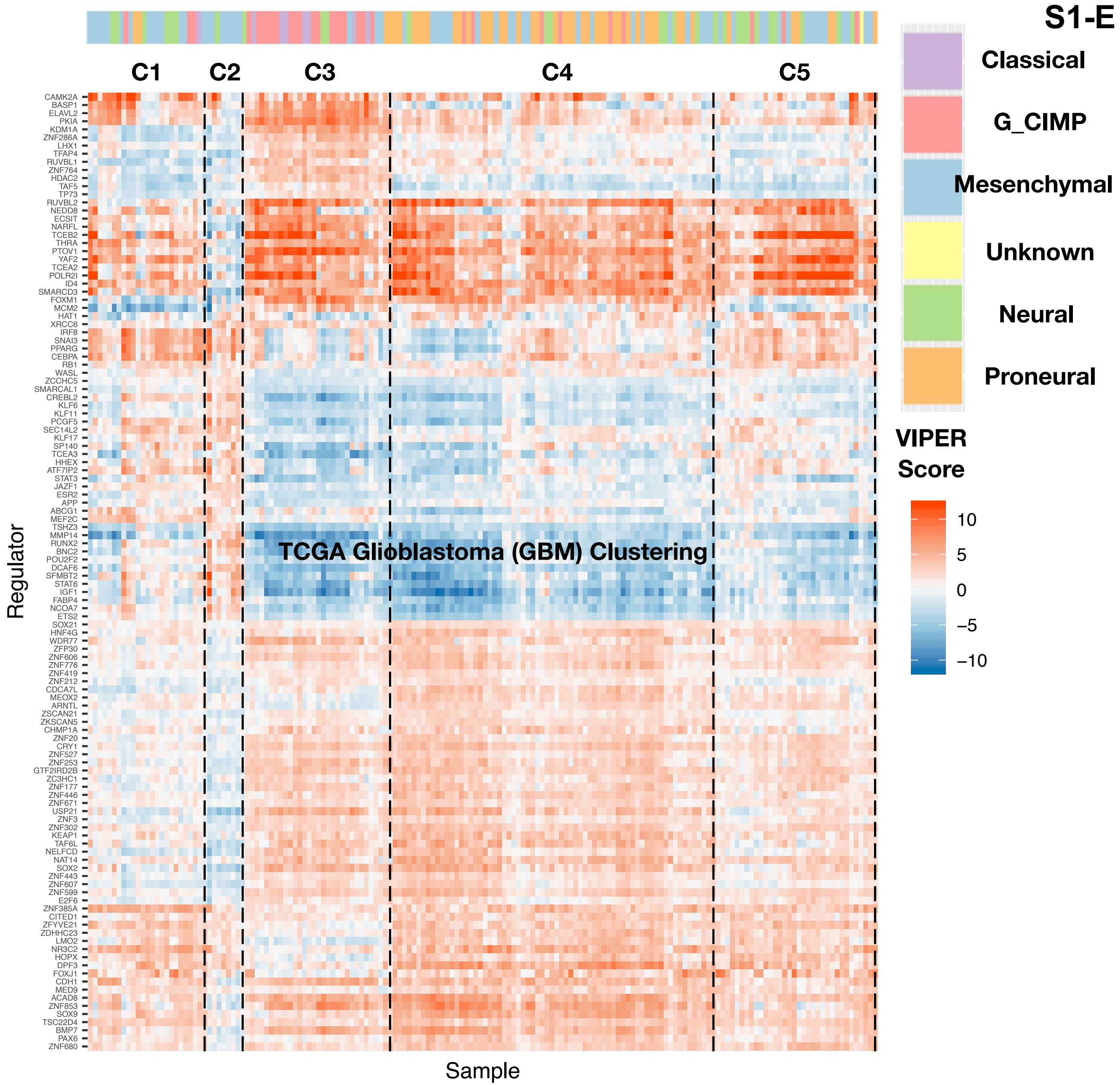
S1-E
C1
C2
C3
C4
C5
Classical
G_CIMP
Mesenchymal
Unknown
Neural
Proneural
VIPER Score
10
5
0
−5
−10
TCGA Glioblastoma (GBM) Clustering
Regulator
Sample
CAMK2A
BASP1
ELAVL2
PKIA
KDM1A
ZNF286A
LHX1
TFAP4
RUVBL1
ZNF764
HDAC2
TAF5
TP73
RUVBL2
NEDD8
ECSIT
NARFL
TCEB2
THRA
PTOV1
YAF2
TCEA2
POLR2I
ID4
SMARCD3
FOXM1
MCM2
HAT1
XRCC6
IRF8
SNAI3
PPARG
CEBPA
RB1
WASL
ZCCHC5
SMARCAL1
CREBL2
KLF6
KLF11
PCGF5
SEC14L2
KLF17
SP140
TCEA3
HHEX
ATF7IP2
STAT3
JAZF1
ESR2
APP
ABCG1
MEF2C
TSHZ3
MMP14
RUNX2
BNC2
POU2F2
DCAF6
SFMBT2
STAT6
IGF1
FABP4
NCOA7
ETS2
SOX21
HNF4G
WDR77
ZFP30
ZNF606
ZNF776
ZNF419
ZNF212
CDCA7L
MEOX2
ARNTL
ZSCAN21
ZKSCAN5
CHMP1A
ZNF20
CRY1
ZNF527
ZNF253
GTF2IRD2B
ZC3HC1
ZNF177
ZNF446
ZNF671
USP21
ZNF3
ZNF302
KEAP1
TAF6L
NELFCD
NAT14
SOX2
ZNF443
ZNF607
ZNF599
E2F6
ZNF385A
CITED1
ZFYVE21
ZDHHC23
LMO2
NR3C2
HOPX
DPF3
FOXJ1
CDH1
MED9
ACAD8
ZNF853
SOX9
TSC22D4
BMP7
PAX6
ZNF680

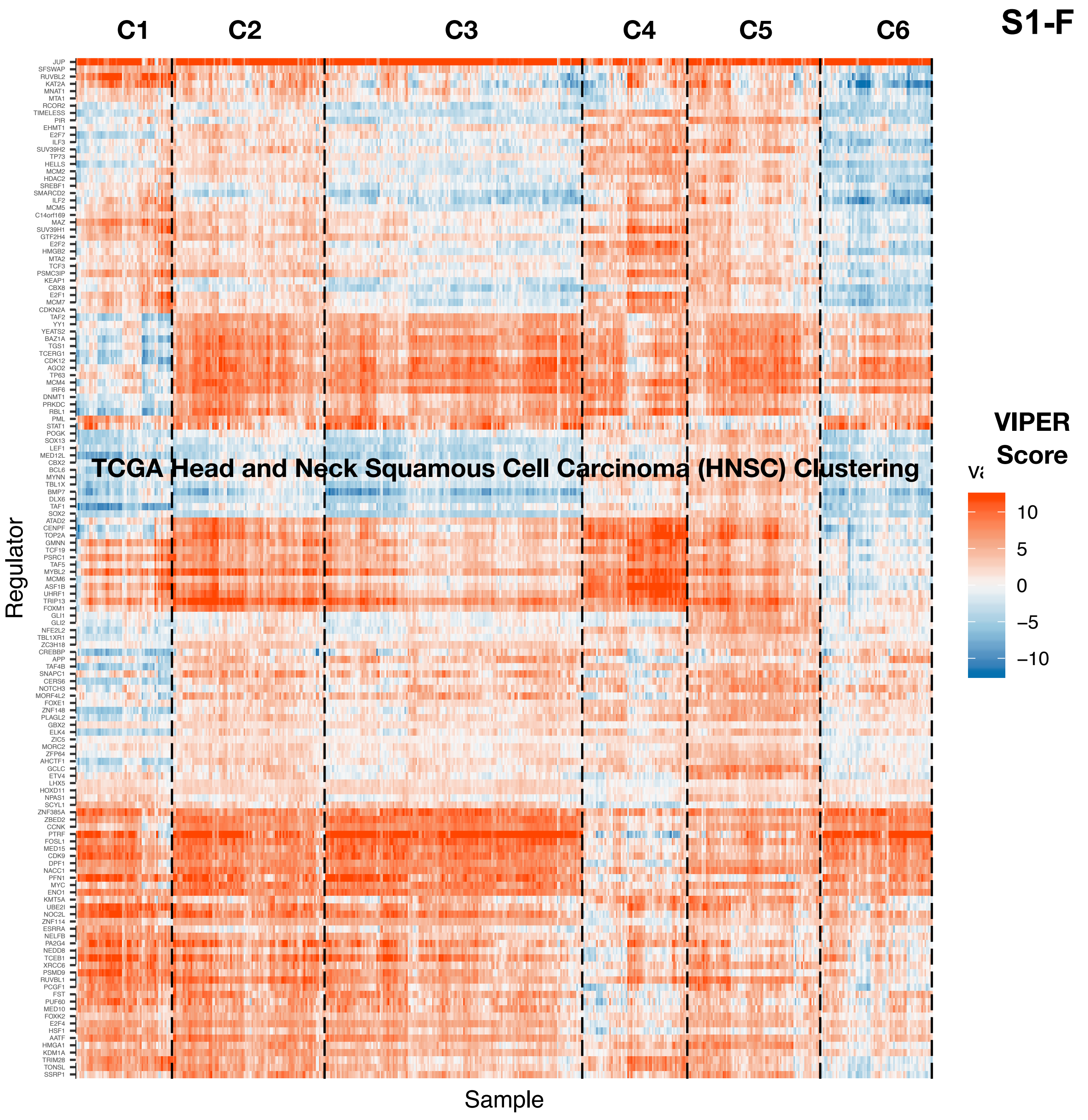
S1-F
TCGA Head and Neck Squamous Cell Carcinoma (HNSC) Clustering
C1
C2
C3
C4
C5
C6
Regulator
Sample
VIPER Score
10
5
0
−5
−10
JUP
SFSWAP
RUVBL2
KAT2A
MNAT1
MTA1
RCOR2
TIMELESS
PIR
EHMT1
E2F7
ILF3
SUV39H2
TP73
HELLS
MCM2
HDAC2
SREBF1
SMARCD2
ILF2
MCM5
C14orf169
MAZ
SUV39H1
GTF2H4
E2F2
HMGB2
MTA2
TCF3
PSMC3IP
KEAP1
CBX8
E2F1
MCM7
CDKN2A
TAF2
YY1
YEATS2
BAZ1A
TGS1
TCERG1
CDK12
AGO2
TP63
MCM4
IRF6
DNMT1
PRKDC
RBL1
PML
STAT1
POGK
SOX13
LEF1
MED12L
CBX2
BCL6
MYNN
TBL1X
BMP7
DLX6
TAF1
SOX2
ATAD2
CENPF
TOP2A
GMNN
TCF19
PSRC1
TAF5
MYBL2
MCM6
ASF1B
UHRF1
TRIP13
FOXM1
GLI1
GLI2
NFE2L2
TBL1XR1
ZC3H18
CREBBP
APP
TAF4B
SNAPC1
CERS6
NOTCH3
MORF4L2
FOXE1
ZNF148
PLAGL2
GBX2
ELK4
ZIC5
MORC2
ZFP64
AHCTF1
GCLC
ETV4
LHX5
HOXD11
NPAS1
SCYL1
ZNF385A
ZBED2
CCNK
PTRF
FOSL1
MED15
CDK9
DPF1
NACC1
PFN1
MYC
ENO1
KMT5A
UBE2I
NOC2L
ZNF114
ESRRA
NELFB
PA2G4
NEDD8
TCEB1
XRCC6
PSMD9
RUVBL1
PCGF1
FST
PUF60
MED10
FOXK2
E2F4
HSF1
AATF
HMGA1
KDM1A
TRIM28
TONSL
SSRP1

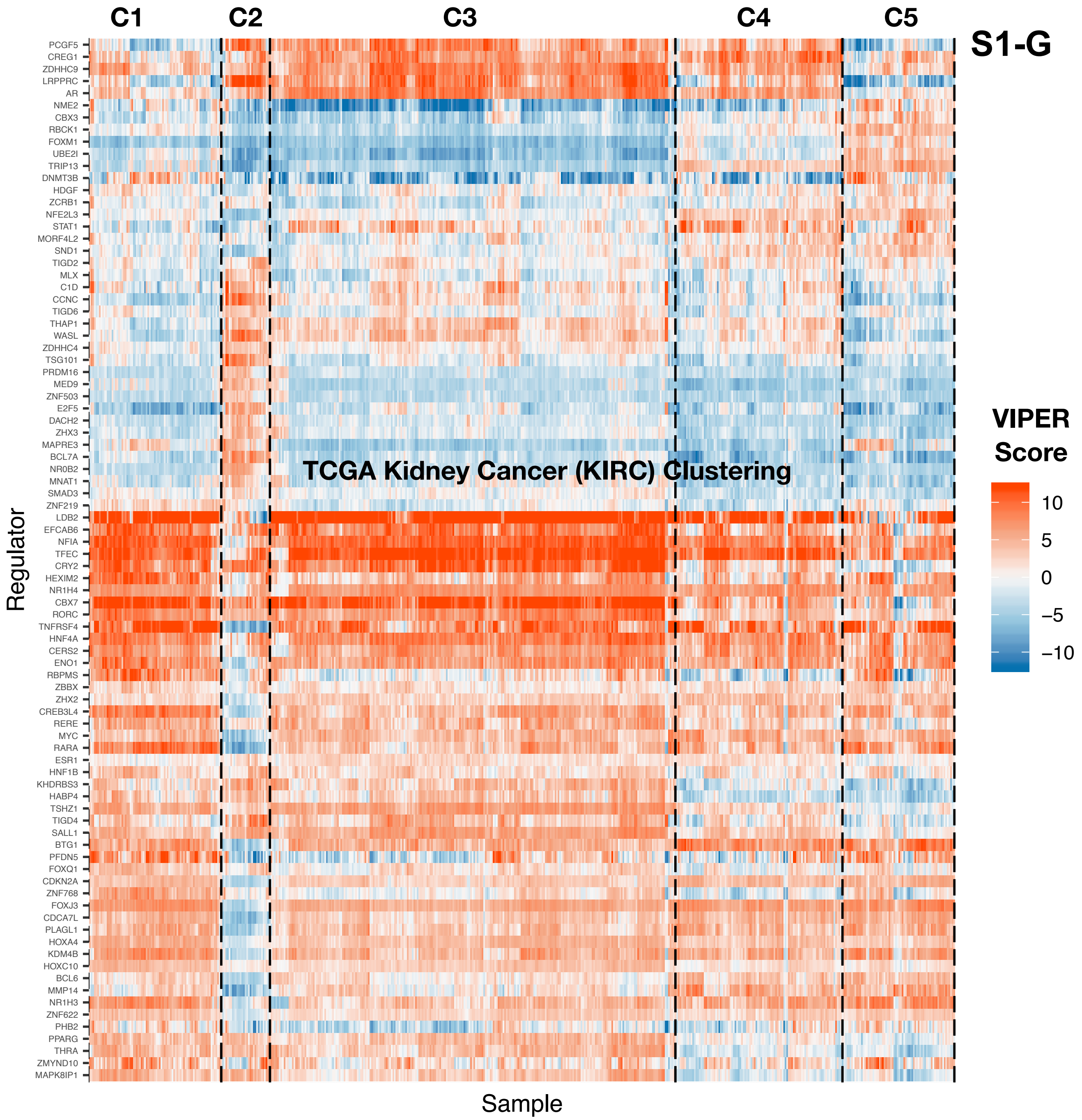
C1
C2
C3
C4
C5
S1-G
TCGA Kidney Cancer (KIRC) Clustering
Regulator
Sample
VIPER Score
10
5
0
−5
−10
PCGF5
CREG1
ZDHHC9
LRPPRC
AR
NME2
CBX3
RBCK1
FOXM1
UBE2I
TRIP13
DNMT3B
HDGF
ZCRB1
NFE2L3
STAT1
MORF4L2
SND1
TIGD2
MLX
C1D
CCNC
TIGD6
THAP1
WASL
ZDHHC4
TSG101
PRDM16
MED9
ZNF503
E2F5
DACH2
ZHX3
MAPRE3
BCL7A
NR0B2
MNAT1
SMAD3
ZNF219
LDB2
EFCAB6
NFIA
TFEC
CRY2
HEXIM2
NR1H4
CBX7
RORC
TNFRSF4
HNF4A
CERS2
ENO1
RBPMS
ZBBX
ZHX2
CREB3L4
RERE
MYC
RARA
ESR1
HNF1B
KHDRBS3
HABP4
TSHZ1
TIGD4
SALL1
BTG1
PFDN5
FOXQ1
CDKN2A
ZNF768
FOXJ3
CDCA7L
PLAGL1
HOXA4
KDM4B
HOXC10
BCL6
MMP14
NR1H3
ZNF622
PHB2
PPARG
THRA
ZMYND10
MAPK8IP1

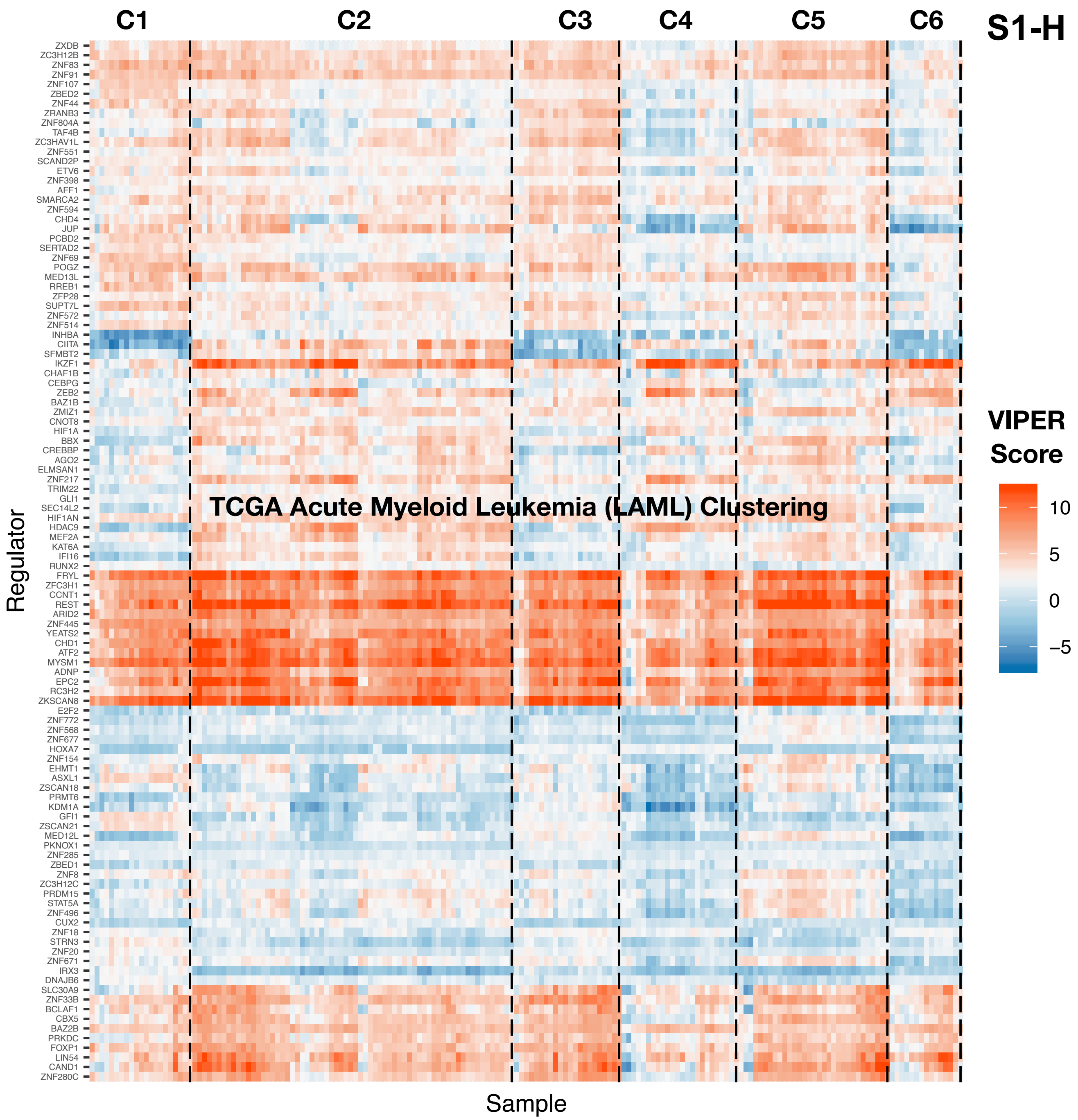

S1-H
C1
C2
C3
C4
C5
C6
TCGA Acute Myeloid Leukemia (LAML) Clustering
Regulator
Sample
VIPER Score
10
5
0
−5
ZXDB
ZC3H12B
ZNF83
ZNF91
ZNF107
ZBED2
ZNF44
ZRANB3
ZNF804A
TAF4B
ZC3HAV1L
ZNF551
SCAND2P
ETV6
ZNF398
AFF1
SMARCA2
ZNF594
CHD4
JUP
PCBD2
SERTAD2
ZNF69
POGZ
MED13L
RREB1
ZFP28
SUPT7L
ZNF572
ZNF514
INHBA
CIITA
SFMBT2
IKZF1
CHAF1B
CEBPG
ZEB2
BAZ1B
ZMIZ1
CNOT8
HIF1A
BBX
CREBBP
AGO2
ELMSAN1
ZNF217
TRIM22
GLI1
SEC14L2
HIF1AN
HDAC9
MEF2A
KAT6A
IFI16
RUNX2
FRYL
ZFC3H1
CCNT1
REST
ARID2
ZNF445
YEATS2
CHD1
ATF2
MYSM1
ADNP
EPC2
RC3H2
ZKSCAN8
E2F2
ZNF772
ZNF568
ZNF677
HOXA7
ZNF154
EHMT1
ASXL1
ZSCAN18
PRMT6
KDM1A
GFI1
ZSCAN21
MED12L
PKNOX1
ZNF285
ZBED1
ZNF8
ZC3H12C
PRDM15
STAT5A
ZNF496
CUX2
ZNF18
STRN3
ZNF20
ZNF671
IRX3
DNAJB6
SLC30A9
ZNF33B
BCLAF1
CBX5
BAZ2B
PRKDC
FOXP1
LIN54
CAND1
ZNF280C

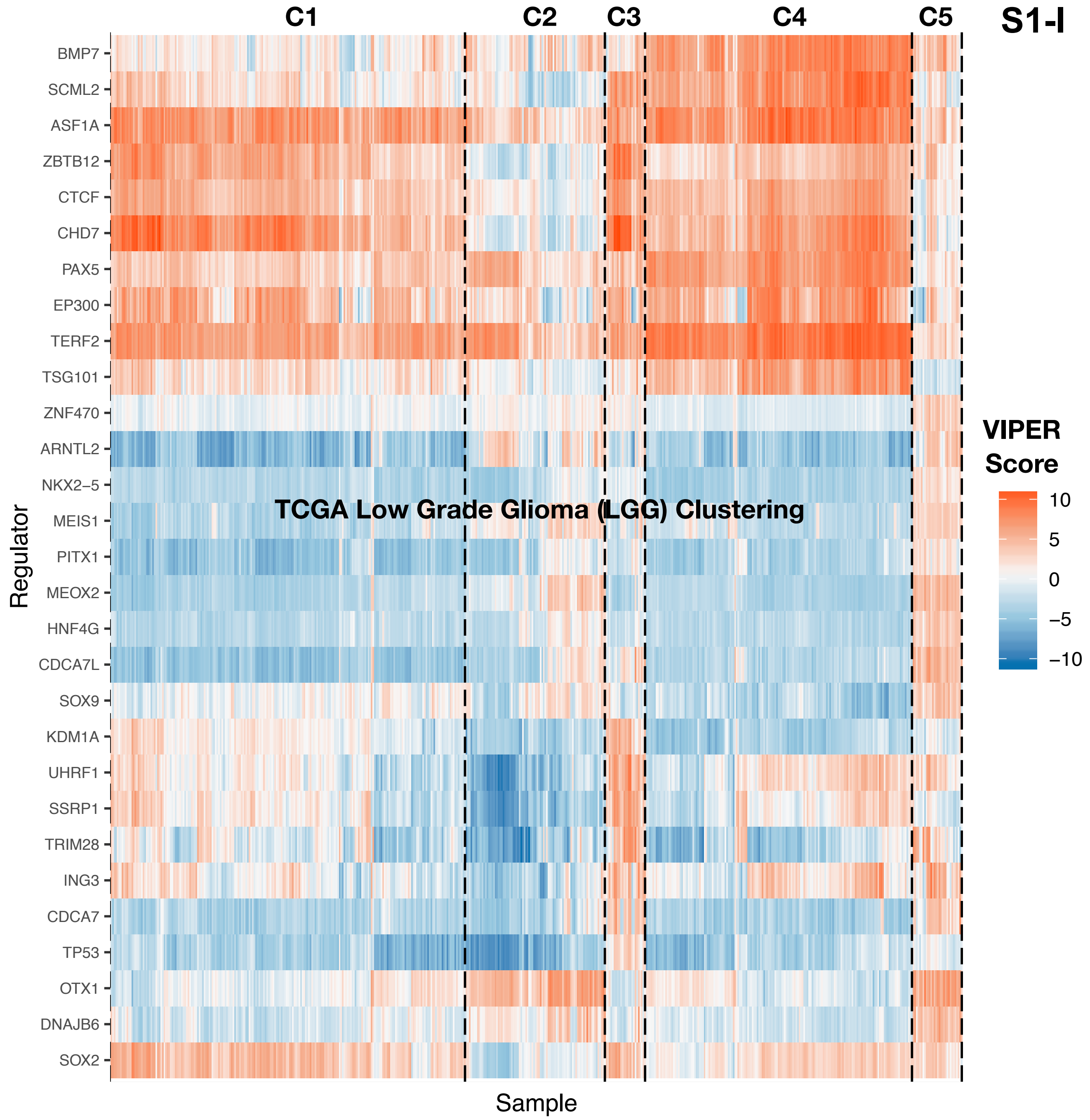

S1-I
TCGA Low Grade Glioma (LGG) Clustering
C1
C2
C3
C4
C5
Regulator
BMP7
SCML2
ASF1A
ZBTB12
CTCF
CHD7
PAX5
EP300
TERF2
TSG101
ZNF470
ARNTL2
NKX2–5
MEIS1
PITX1
MEOX2
HNF4G
CDCA7L
SOX9
KDM1A
UHRF1
SSRP1
TRIM28
ING3
CDCA7
TP53
OTX1
DNAJB6
SOX2
Sample
VIPER Score
10
5
0
−5
−10

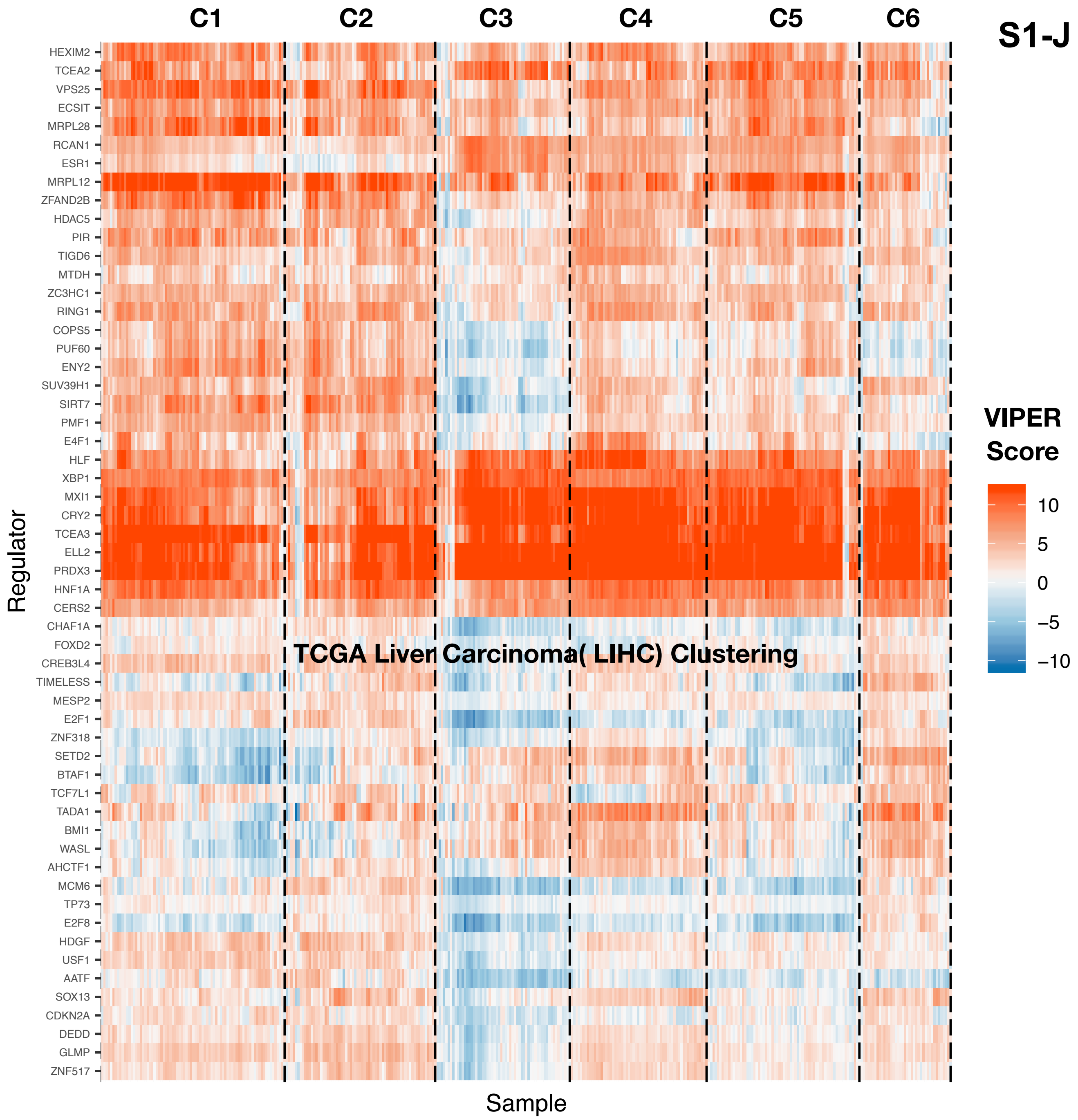
C1
C2
C3
C4
C5
C6
S1-J
HEXIM2
TCEA2
VPS25
ECSIT
MRPL28
RCAN1
ESR1
MRPL12
ZFAND2B
HDAC5
PIR
TIGD6
MTDH
ZC3HC1
RING1
COPS5
PUF60
ENY2
SUV39H1
SIRT7
PMF1
E4F1
HLF
XBP1
MXI1
CRY2
TCEA3
ELL2
PRDX3
HNF1A
CERS2
CHAF1A
FOXD2
CREB3L4
TIMELESS
MESP2
E2F1
ZNF318
SETD2
BTAF1
TCF7L1
TADA1
BMI1
WASL
AHCTF1
MCM6
TP73
E2F8
HDGF
USF1
AATF
SOX13
CDKN2A
DEDD
GLMP
ZNF517
Regulator
Sample
TCGA Liver Carcinoma( LIHC) Clustering
VIPER Score
10
5
0
−5
−10

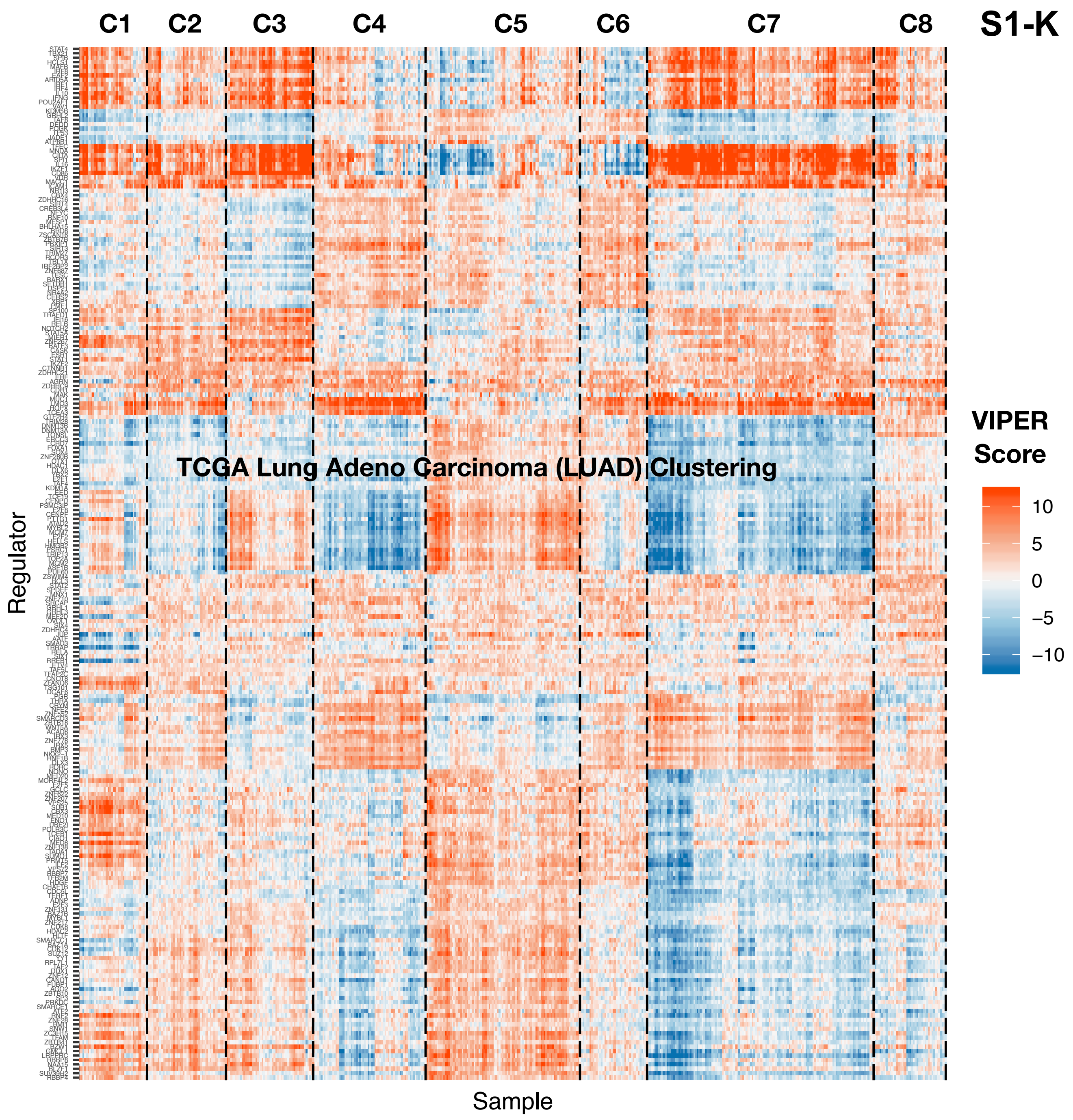

S1-K
C1
C2
C3
C4
C5
C6
C7
C8
TCGA Lung Adeno Carcinoma (LUAD) Clustering
Regulator
Sample
VIPER Score
10
5
0
−5
−10

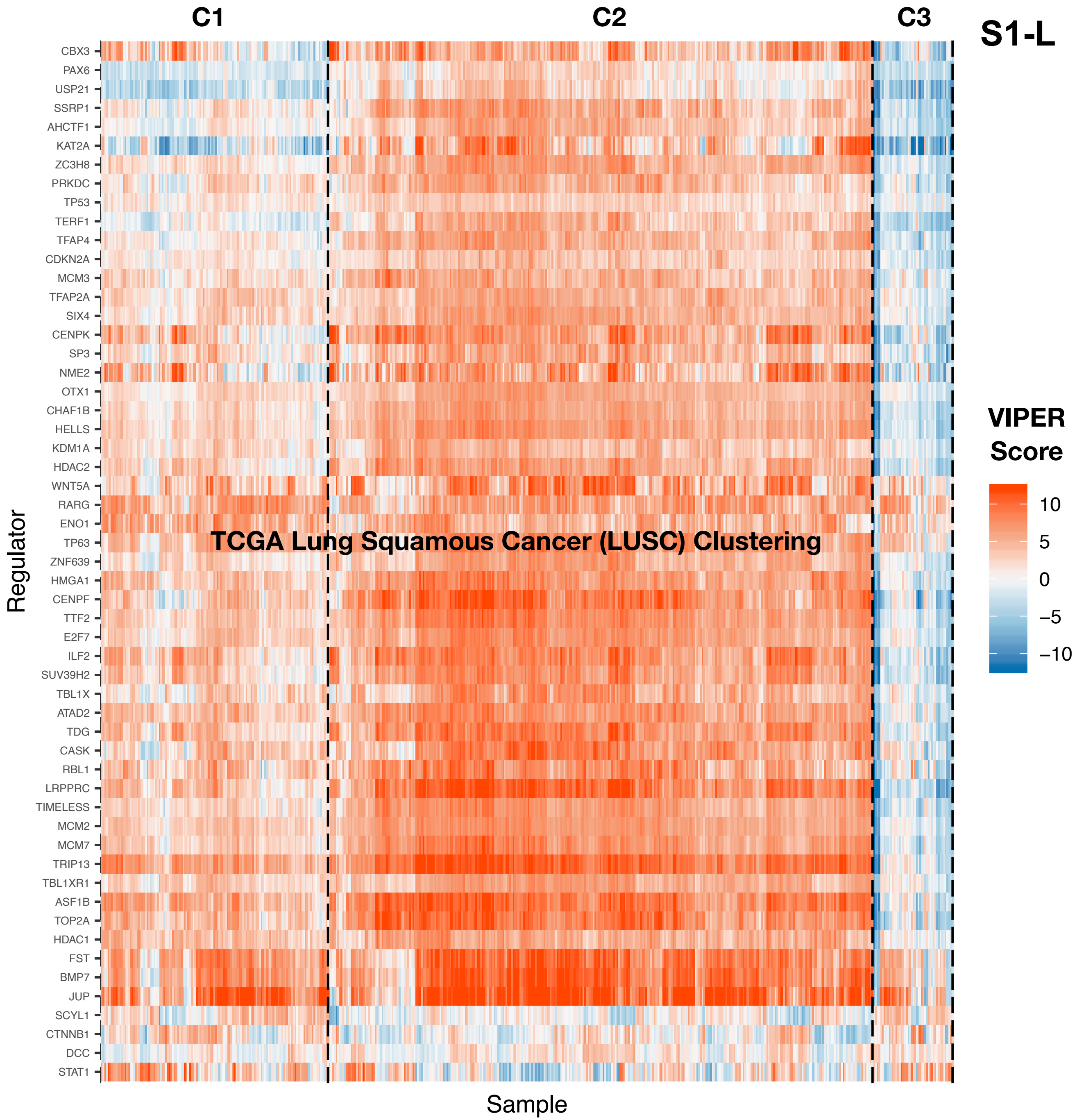
S1-L
C1
C2
C3
TCGA Lung Squamous Cancer (LUSC) Clustering
Regulator
Sample
CBX3
PAX6
USP21
SSRP1
AHCTF1
KAT2A
ZC3H8
PRKDC
TP53
TERF1
TFAP4
CDKN2A
MCM3
TFAP2A
SIX4
CENPK
SP3
NME2
OTX1
CHAF1B
HELLS
KDM1A
HDAC2
WNT5A
RARG
ENO1
TP63
ZNF639
HMGA1
CENPF
TTF2
E2F7
ILF2
SUV39H2
TBL1X
ATAD2
TDG
CASK
RBL1
LRPPRC
TIMELESS
MCM2
MCM7
TRIP13
TBL1XR1
ASF1B
TOP2A
HDAC1
FST
BMP7
JUP
SCYL1
CTNNB1
DCC
STAT1
VIPER Score
10
5
0
−5
−10

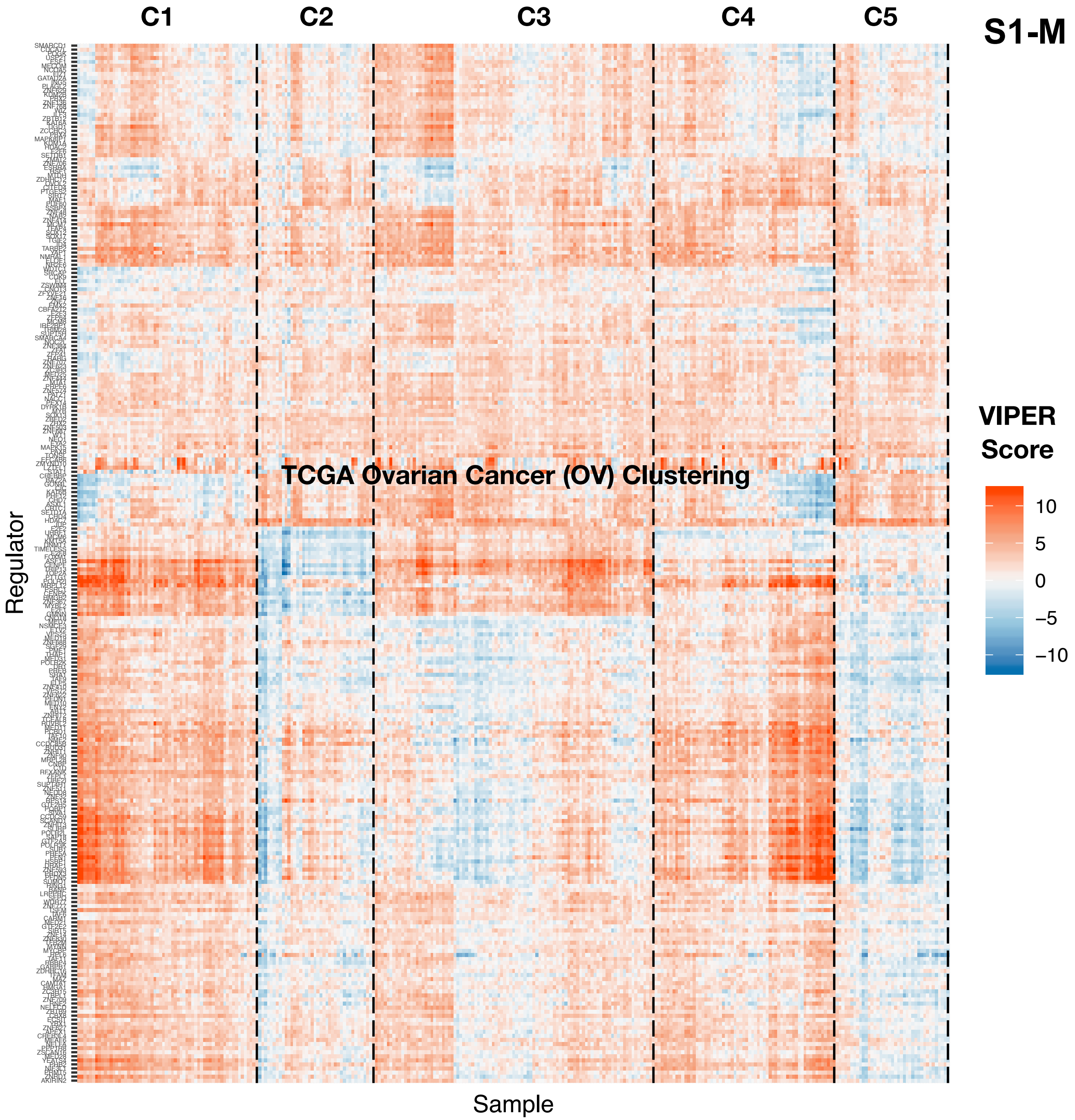

S1-M
C1
C2
C3
C4
C5
TCGA Ovarian Cancer (OV) Clustering
Regulator
Sample
VIPER Score
10
5
0
−5
−10

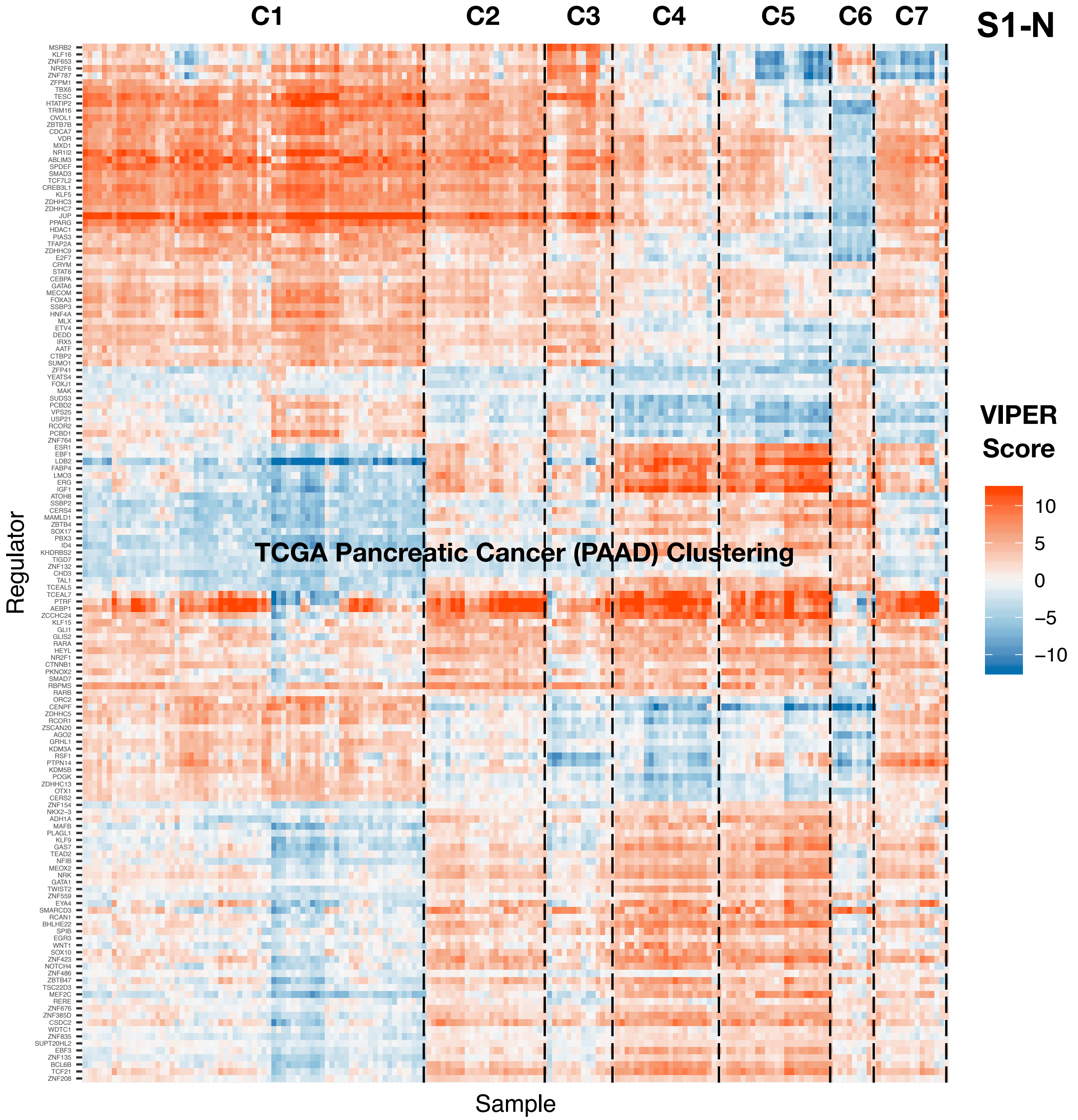
S1-N
C1
C2
C3
C4
C5
C6
C7
TCGA Pancreatic Cancer (PAAD) Clustering
Regulator
Sample
VIPER Score
10
5
0
−5
−10
MSRB2
KLF16
ZNF653
NR2F6
ZNF787
ZFPM1
TBX6
TESC
HTATIP2
TRIM16
OVOL1
ZBTB7B
CDCA7
VDR
MXD1
NR1I2
ABLIM3
SPDEF
SMAD3
TCF7L2
CREB3L1
KLF5
ZDHHC3
ZDHHC7
JUP
PPARG
HDAC1
PIAS3
TFAP2A
ZDHHC9
E2F7
CRYM
STAT6
CEBPA
GATA6
MECOM
FOXA3
SSBP3
HNF4A
MLX
ETV4
DEDD
IRX5
AATF
CTBP2
SUMO1
ZFP41
YEATS4
FOXJ1
MAK
SUDS3
PCBD2
VPS25
USP21
RCOR2
PCBD1
ZNF764
ESR1
EBF1
LDB2
FABP4
LMO3
ERG
IGF1
ATOH8
SSBP2
CERS4
MAMLD1
ZBTB4
SOX17
PBX3
ID4
KHDRBS2
TIGD7
ZNF132
CHD3
TAL1
TCEAL5
TCEAL7
PTRF
AEBP1
ZCCHC24
KLF15
GLI1
GLIS2
RARA
HEYL
NR2F1
CTNNB1
PKNOX2
SMAD7
RBPMS
RARB
ORC2
CENPF
ZDHHC5
RCOR1
ZSCAN20
AGO2
GRHL1
KDM3A
RSF1
PTPN14
KDM5B
POGK
ZDHHC13
OTX1
CERS2
ZNF154
NKX2-3
ADH1A
MAFB
PLAGL1
KLF9
GAS7
TEAD2
NFIB
MEOX2
NRK
GATA1
TWIST2
ZNF559
EYA4
SMARCD3
RCAN1
BHLHE22
SPIB
EGR3
WNT1
SOX10
ZNF423
NOTCH4
ZNF486
ZBTB47
TSC22D3
MEF2C
RERE
ZNF676
ZNF385D
CSDC2
WDTC1
ZNF835
SUPT20HL2
EBF3
ZNF135
BCL6B
TCF21
ZNF208

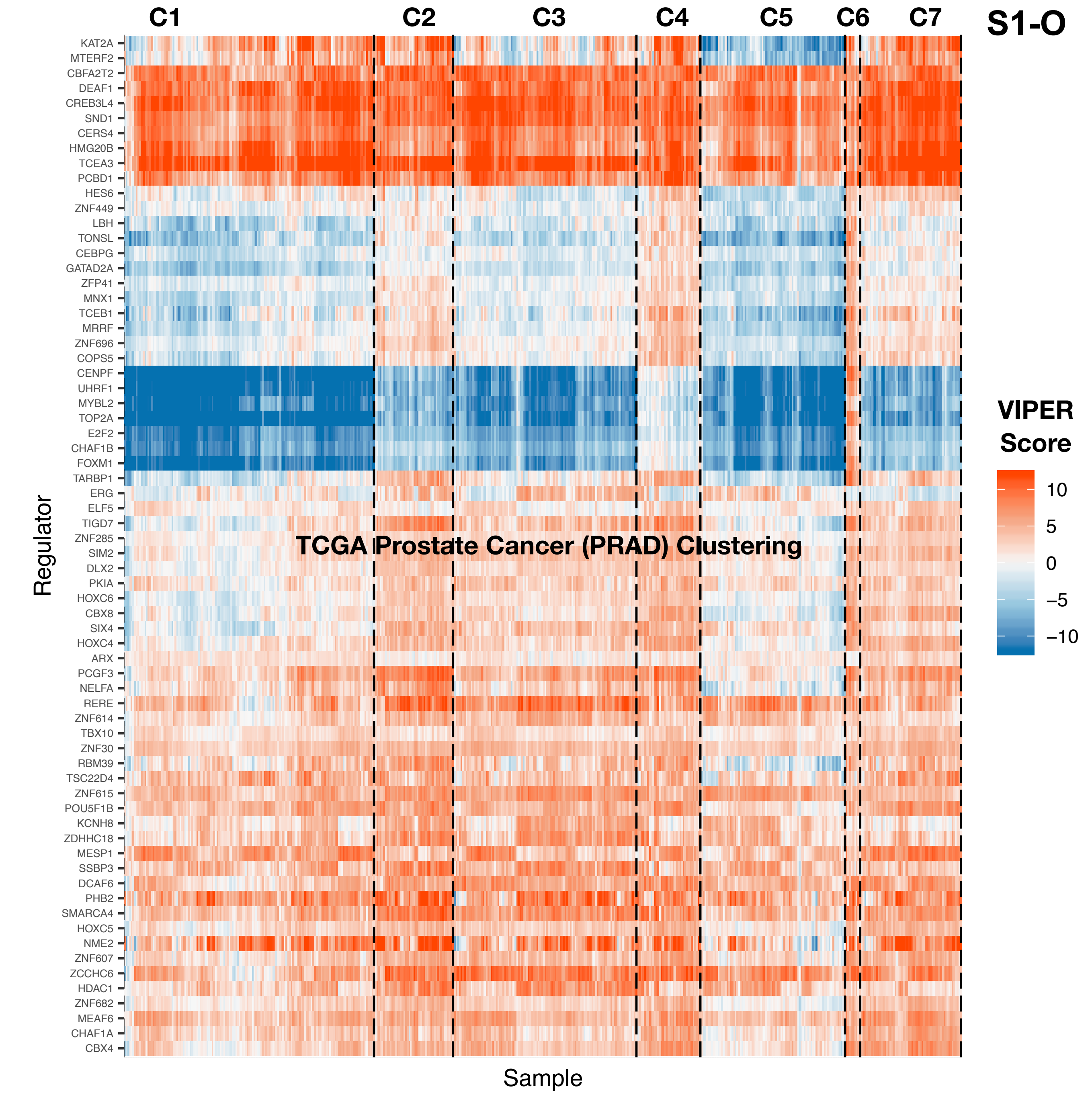
S1-O
C1
C2
C3
C4
C5
C6
C7
TCGA Prostate Cancer (PRAD) Clustering
KAT2A
MTERF2
CBFA2T2
DEAF1
CREB3L4
SND1
CERS4
HMG20B
TCEA3
PCBD1
HES6
ZNF449
LBH
TONSL
CEBPG
GATAD2A
ZFP41
MNX1
TCEB1
MRRF
ZNF696
COPS5
CENPF
UHRF1
MYBL2
TOP2A
E2F2
CHAF1B
FOXM1
TARBP1
ERG
ELF5
TIGD7
ZNF285
SIM2
DLX2
PKIA
HOXC6
CBX8
SIX4
HOXC4
ARX
PCGF3
NELFA
RERE
ZNF614
TBX10
ZNF30
RBM39
TSC22D4
ZNF615
POU5F1B
KCNH8
ZDHHC18
MESP1
SSBP3
DCAF6
PHB2
SMARCA4
HOXC5
NME2
ZNF607
ZCCHC6
HDAC1
ZNF682
MEAF6
CHAF1A
CBX4
Regulator
Sample
VIPER Score
10
5
0
−5
−10

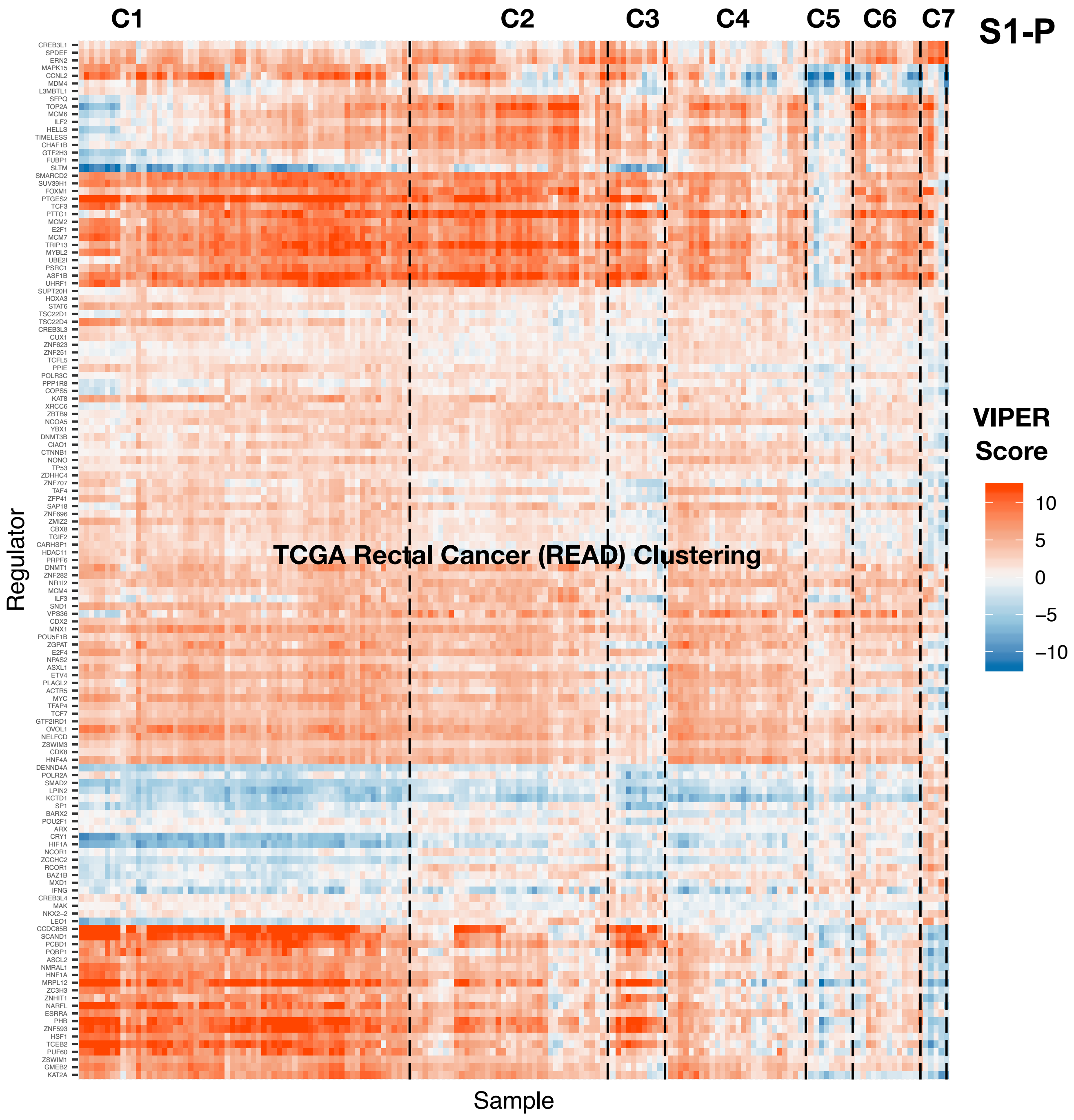

C1
C2
C3
C4
C5
C6
C7
S1-P
TCGA Rectal Cancer (READ) Clustering
Regulator
Sample
CREB3L1
SPDEF
ERN2
MAPK15
CCNL2
MDM4
L3MBTL1
SFPQ
TOP2A
MCM6
ILF2
HELLS
TIMELESS
CHAF1B
GTF2H3
FUBP1
SLTM
SMARCD2
SUV39H1
FOXM1
PTGES2
TCF3
PTTG1
MCM2
E2F1
MCM7
TRIP13
MYBL2
UBE2I
PSRC1
ASF1B
UHRF1
SUPT20H
HOXA3
STAT6
TSC22D1
TSC22D4
CREB3L3
CUX1
ZNF623
ZNF251
TCFL5
PPIE
POLR3C
PPP1R8
COPS5
KAT8
XRCC6
ZBTB9
NCOA5
YBX1
DNMT3B
CIAO1
CTNNB1
NONO
TP53
ZDHHC4
ZNF707
TAF4
ZFP41
SAP18
ZNF696
ZMIZ2
CBX8
TGIF2
CARHSP1
HDAC11
PRPF6
DNMT1
ZNF282
NR1I2
MCM4
ILF3
SND1
VPS36
CDX2
MNX1
POU5F1B
ZGPAT
E2F4
NPAS2
ASXL1
ETV4
PLAGL2
ACTR5
MYC
TFAP4
TCF7
GTF2IRD1
OVOL1
NELFCD
ZSWIM3
CDK8
HNF4A
DENND4A
POLR2A
SMAD2
LPIN2
KCTD1
SP1
BARX2
POU2F1
ARX
CRY1
HIF1A
NCOR1
ZCCHC2
RCOR1
BAZ1B
MXD1
IFNG
CREB3L4
MAK
NKX2−2
LEO1
CCDC85B
SCAND1
PCBD1
PQBP1
ASCL2
NMRAL1
HNF1A
MRPL12
ZC3H3
ZNHIT1
NARFL
ESRRA
PHB
ZNF593
HSF1
TCEB2
PUF60
ZSWIM1
GMEB2
KAT2A
VIPER Score
10
5
0
−5
−10

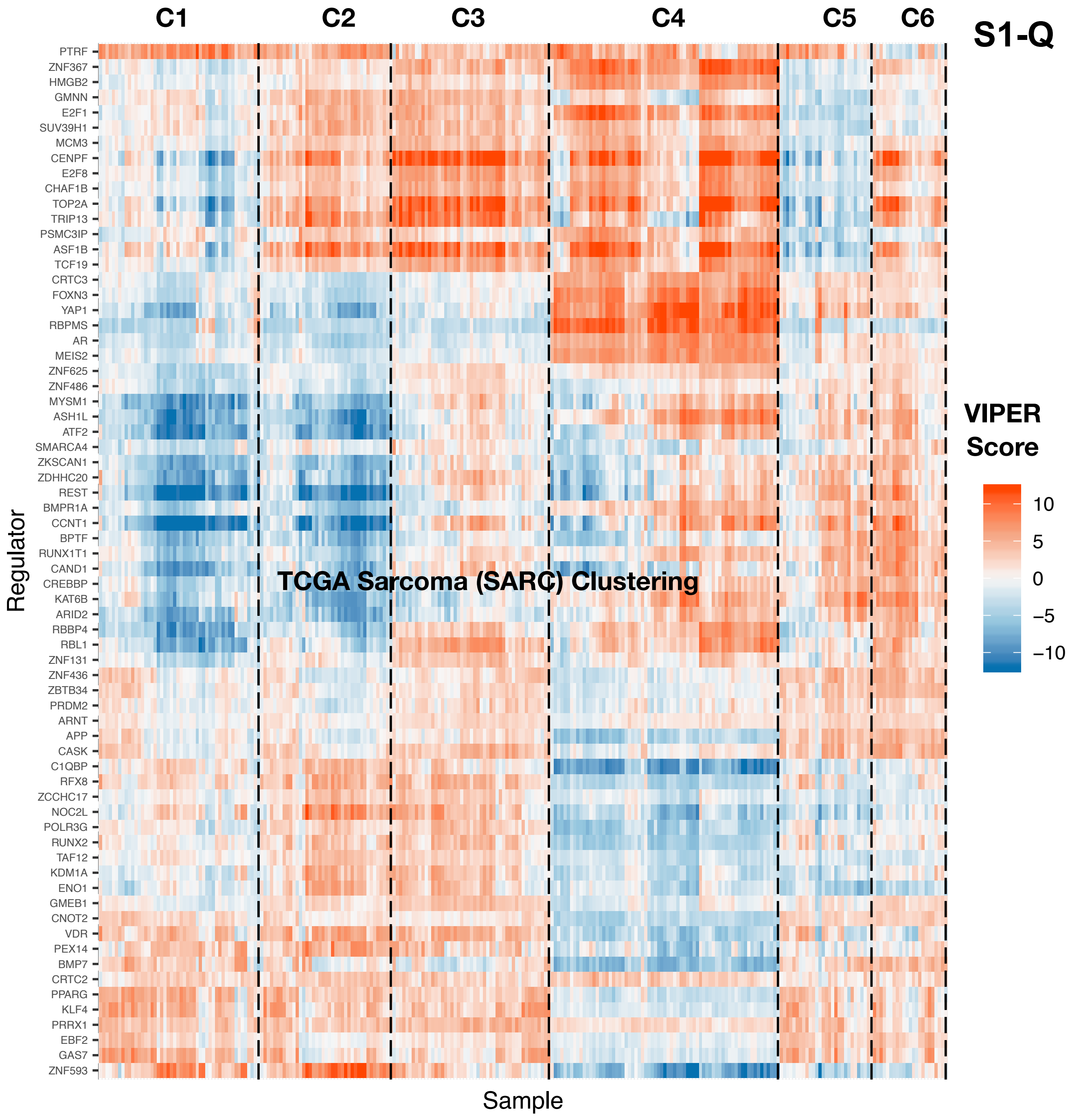

S1-Q
C1
C2
C3
C4
C5
C6
TCGA Sarcoma (SARC) Clustering
Regulator
Sample
VIPER Score
10
5
0
−5
−10
PTRF
ZNF367
HMGB2
GMNN
E2F1
SUV39H1
MCM3
CENPF
E2F8
CHAF1B
TOP2A
TRIP13
PSMC3IP
ASF1B
TCF19
CRTC3
FOXN3
YAP1
RBPMS
AR
MEIS2
ZNF625
ZNF486
MYSM1
ASH1L
ATF2
SMARCA4
ZKSCAN1
ZDHHC20
REST
BMPR1A
CCNT1
BPTF
RUNX1T1
CAND1
CREBBP
KAT6B
ARID2
RBBP4
RBL1
ZNF131
ZNF436
ZBTB34
PRDM2
ARNT
APP
CASK
C1QBP
RFX8
ZCCHC17
NOC2L
POLR3G
RUNX2
TAF12
KDM1A
ENO1
GMEB1
CNOT2
VDR
PEX14
BMP7
CRTC2
PPARG
KLF4
PRRX1
EBF2
GAS7
ZNF593

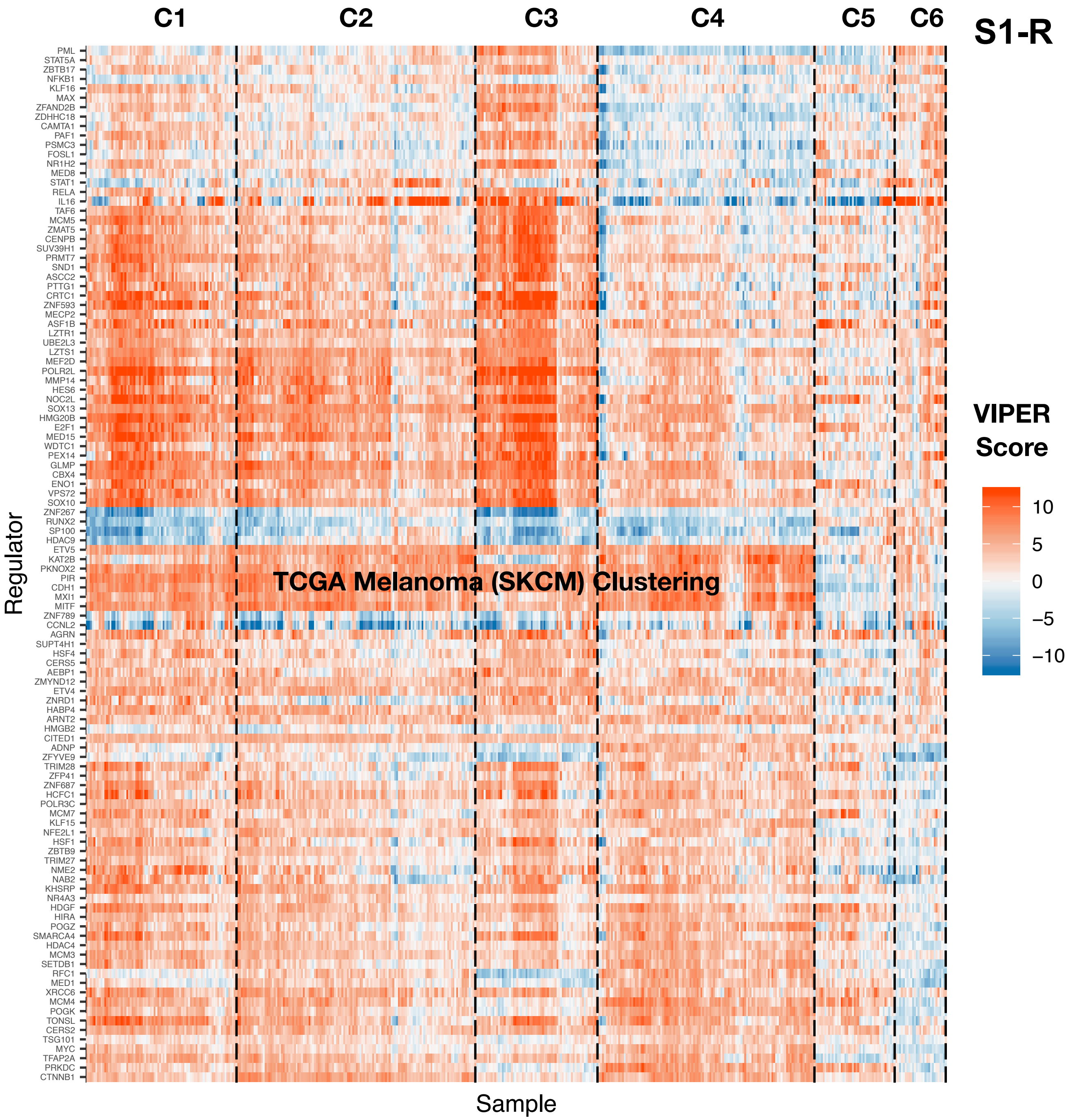

C1
C2
C3
C4
C5
C6
S1-R
TCGA Melanoma (SKCM) Clustering
Regulator
Sample
VIPER Score
10
5
0
−5
−10
PML
STAT5A
ZBTB17
NFKB1
KLF16
MAX
ZFAND2B
ZDHHC18
CAMTA1
PAF1
PSMC3
FOSL1
NR1H2
MED8
STAT1
RELA
IL16
TAF6
MCM5
ZMAT5
CENPB
SUV39H1
PRMT7
SND1
ASCC2
PTTG1
CRTC1
ZNF593
MECP2
ASF1B
LZTR1
UBE2L3
LZTS1
MEF2D
POLR2L
MMP14
HES6
NOC2L
SOX13
HMG20B
E2F1
MED15
WDTC1
PEX14
GLMP
CBX4
ENO1
VPS72
SOX10
ZNF267
RUNX2
SP100
HDAC9
ETV5
KAT2B
PKNOX2
PIR
CDH1
MXI1
MITF
ZNF789
CCNL2
AGRN
SUPT4H1
HSF4
CERS5
AEBP1
ZMYND12
ETV4
ZNRD1
HABP4
ARNT2
HMGB2
CITED1
ADNP
ZFYVE9
TRIM28
ZFP41
ZNF687
HCFC1
POLR3C
MCM7
KLF15
NFE2L1
HSF1
ZBTB9
TRIM27
NME2
NAB2
KHSRP
NR4A3
HDGF
HIRA
POGZ
SMARCA4
HDAC4
MCM3
SETDB1
RFC1
MED1
XRCC6
MCM4
POGK
TONSL
CERS2
TSG101
MYC
TFAP2A
PRKDC
CTNNB1

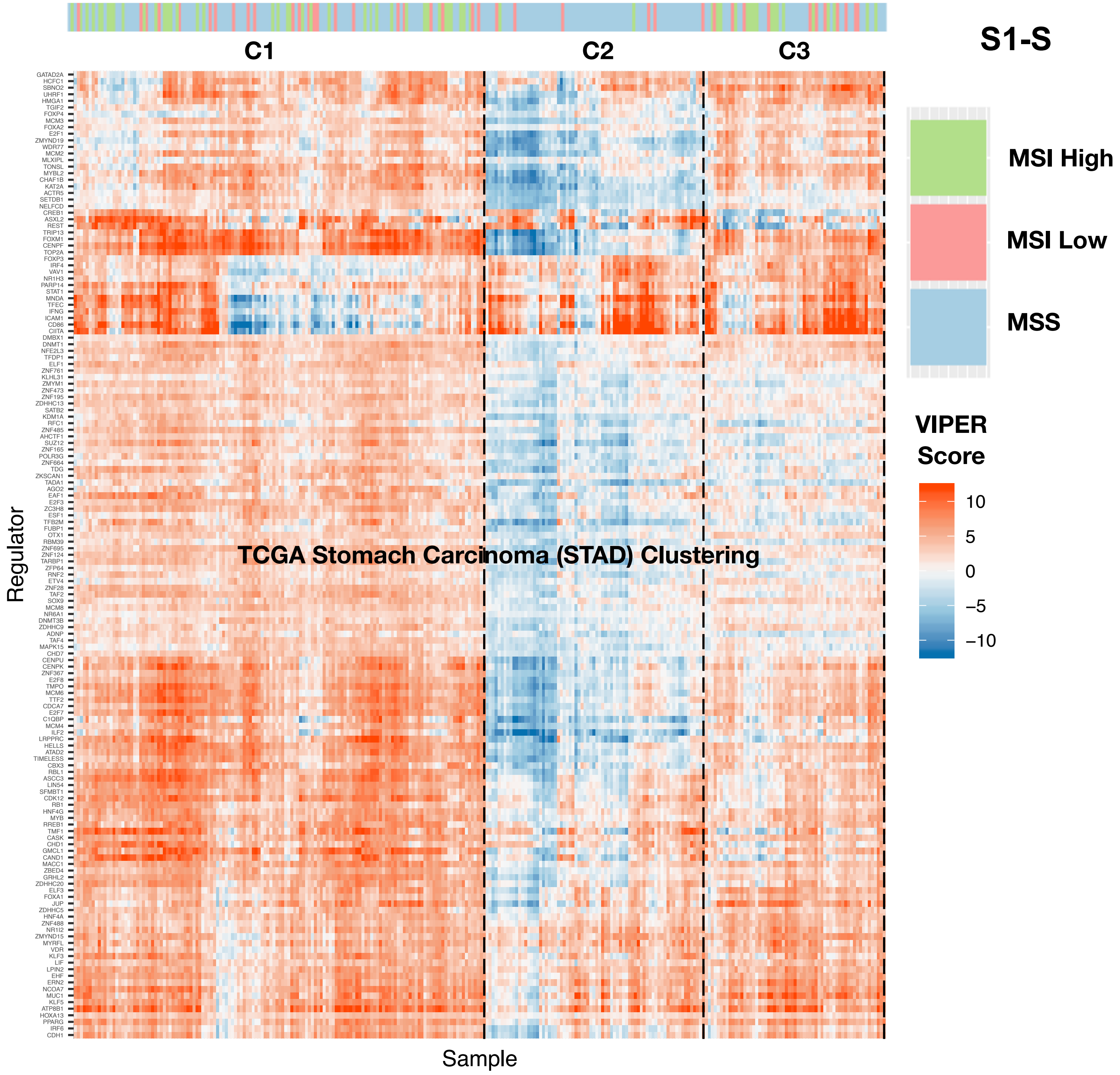
S1-S
C1
C2
C3
TCGA Stomach Carcinoma (STAD) Clustering
Regulator
Sample
MSI High
MSI Low
MSS
VIPER Score
10
5
0
−5
−10
GATAD2A
HCFC1
SBNO2
UHRF1
HMGA1
TGIF2
FOXP4
MCM3
FOXA2
E2F1
ZMYND19
WDR77
MCM2
MLXIPL
TONSL
MYBL2
CHAF1B
KAT2A
ACTR5
SETDB1
NELFCD
CREB1
ASXL2
REST
TRIP13
FOXM1
CENPF
TOP2A
FOXP3
IRF4
VAV1
NR1H3
PARP14
STAT1
MNDA
TFEC
IFNG
ICAM1
CD86
CIITA
DMBX1
DNMT1
NFE2L3
TFDP1
ELF1
ZNF761
KLHL31
ZMYM1
ZNF473
ZNF195
ZDHHC13
SATB2
KDM1A
RFC1
ZNF485
AHCTF1
SUZ12
ZNF165
POLR3G
ZNF664
TDG
ZKSCAN1
TADA1
AGO2
EAF1
E2F3
ZC3H8
ESF1
TFB2M
FUBP1
OTX1
RBM39
ZNF695
ZNF124
TARBP1
ZFP64
RNF2
ETV4
ZNF28
TAF2
SOX9
MCM8
NR6A1
DNMT3B
ZDHHC9
ADNP
TAF4
MAPK15
CHD7
CENPU
CENPK
ZNF367
E2F8
TMPO
MCM6
TTF2
CDCA7
E2F7
C1QBP
MCM4
ILF2
LRPPRC
HELLS
ATAD2
TIMELESS
CBX3
RBL1
ASCC3
LIN54
SFMBT1
CDK12
RB1
HNF4G
MYB
RREB1
TMF1
CASK
CHD1
GMCL1
CAND1
MACC1
ZBED4
GRHL2
ZDHHC20
ELF3
FOXA1
JUP
ZDHHC5
HNF4A
ZNF488
NR1I2
ZMYND15
MYRFL
VDR
KLF3
LIF
LPIN2
EHF
ERN2
NCOA7
MUC1
KLF5
ATP8B1
HOXA13
PPARG
IRF6
CDH1

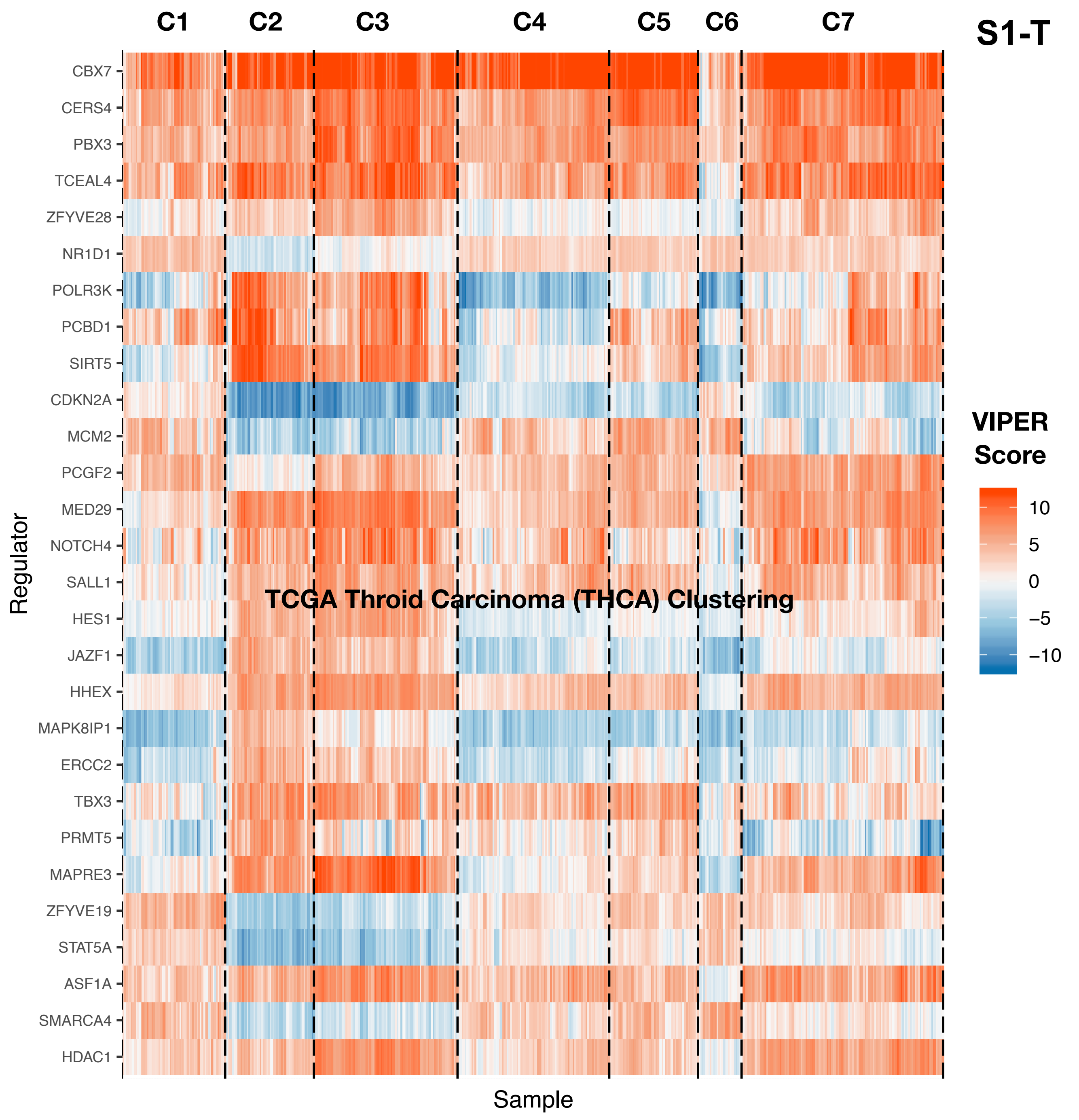

C1
C2
C3
C4
C5
C6
C7
S1-T
CBX7
CERS4
PBX3
TCEAL4
ZFYVE28
NR1D1
POLR3K
PCBD1
SIRT5
CDKN2A
MCM2
PCGF2
MED29
NOTCH4
SALL1
HES1
JAZF1
HHEX
MAPK8IP1
ERCC2
TBX3
PRMT5
MAPRE3
ZFYVE19
STAT5A
ASF1A
SMARCA4
HDAC1
Regulator
Sample
TCGA Throid Carcinoma (THCA) Clustering
VIPER Score
10
5
0
−5
−10

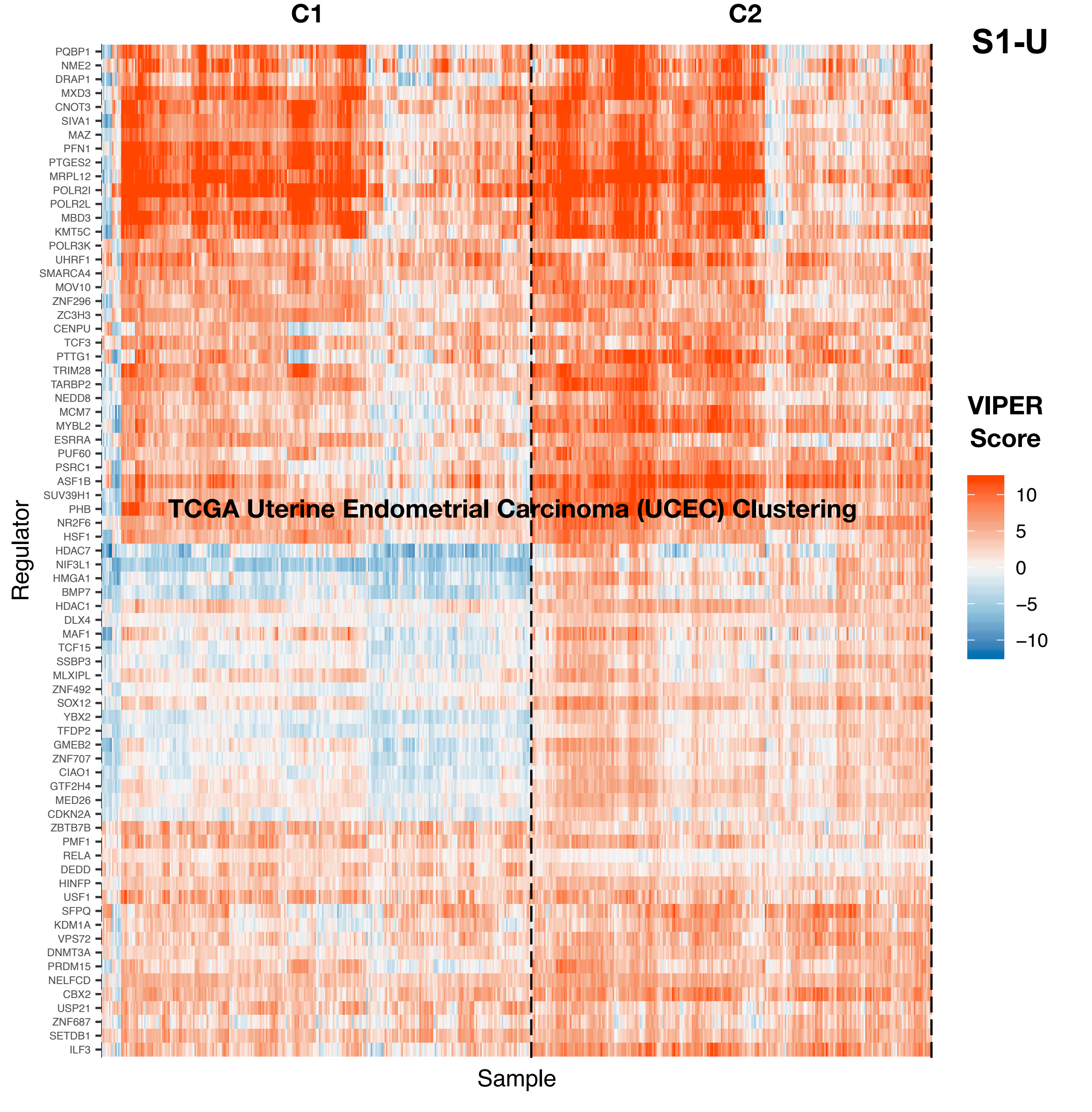

C1
C2
S1-U
TCGA Uterine Endometrial Carcinoma (UCEC) Clustering
Regulator
Sample
VIPER Score
10
5
0
−5
−10
PQBP1
NME2
DRAP1
MXD3
CNOT3
SIVA1
MAZ
PFN1
PTGES2
MRPL12
POLR2I
POLR2L
MBD3
KMT5C
POLR3K
UHRF1
SMARCA4
MOV10
ZNF296
ZC3H3
CENPU
TCF3
PTTG1
TRIM28
TARBP2
NEDD8
MCM7
MYBL2
ESRRA
PUF60
PSRC1
ASF1B
SUV39H1
PHB
NR2F6
HSF1
HDAC7
NIF3L1
HMGA1
BMP7
HDAC1
DLX4
MAF1
TCF15
SSBP3
MLXIPL
ZNF492
SOX12
YBX2
TFDP2
GMEB2
ZNF707
CIAO1
GTF2H4
MED26
CDKN2A
ZBTB7B
PMF1
RELA
DEDD
HINFP
USF1
SFPQ
KDM1A
VPS72
DNMT3A
PRDM15
NELFCD
CBX2
USP21
ZNF687
SETDB1
ILF3

S1-V

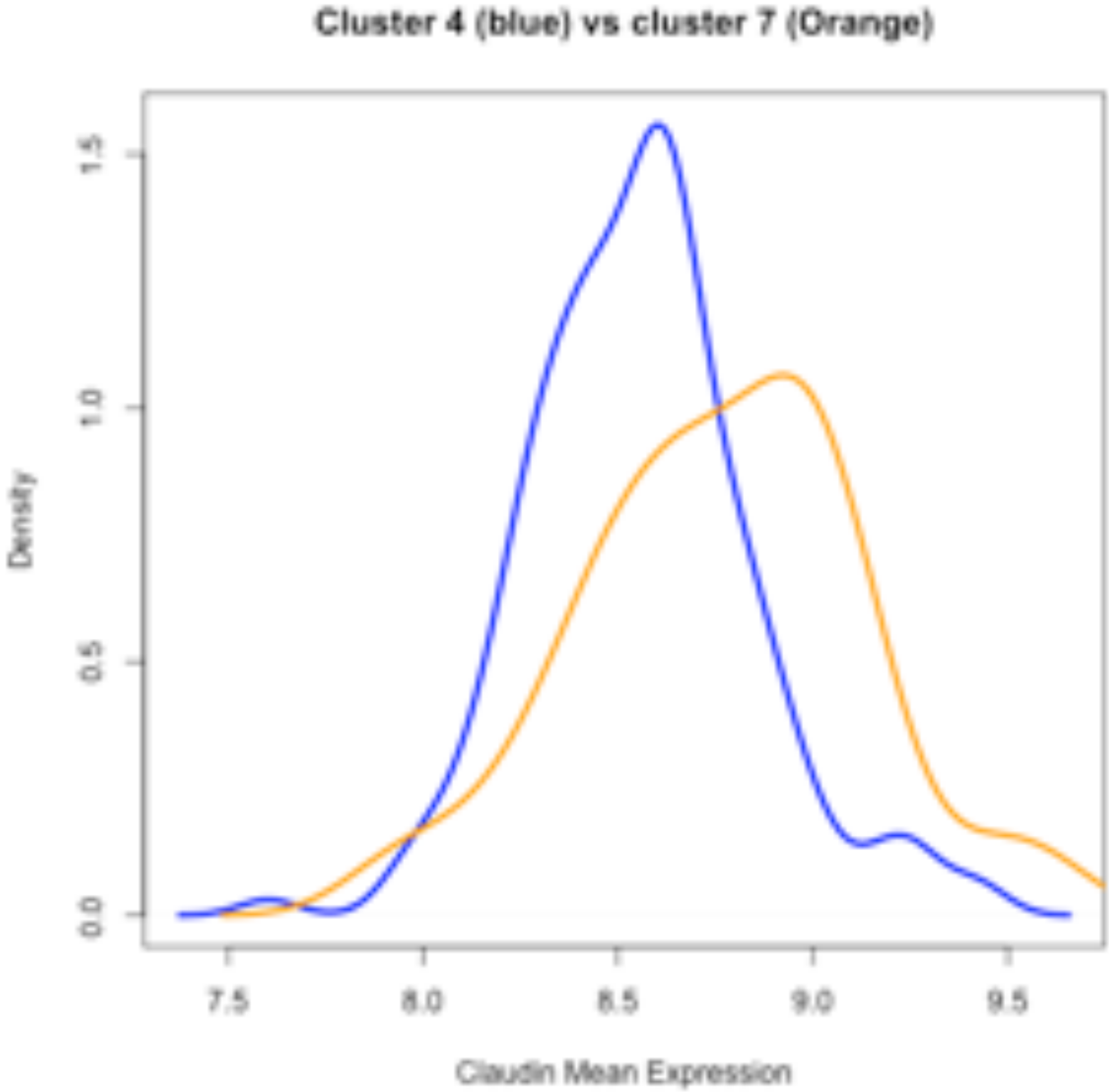

Cluster 4 (blue) vs cluster 7 (Orange)
Density
Claudin Mean Expression
0.0
0.5
1.0
1.5
7.5
8.0
8.5
9.0
9.5

### Figure S2

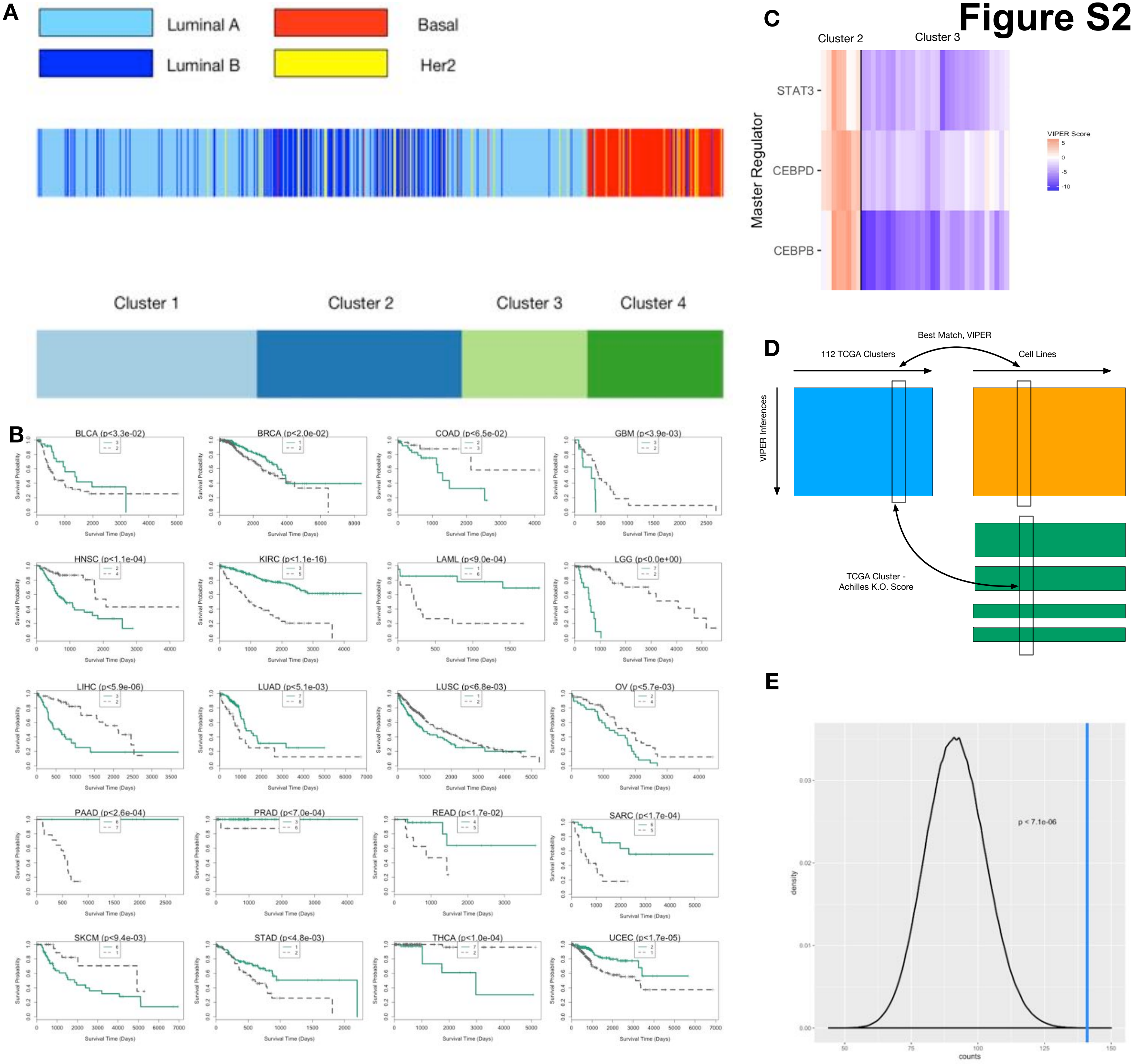
A
Luminal A
Basal
Luminal B
Her2
Cluster 1
Cluster 2
Cluster 3
Cluster 4
B
BLCA (p<3.3e-02)
BRCA (p<2.0e-02)
COAD (p<6.5e-02)
GBM (p<3.9e-03)
HNSC (p<1.1e-04)
KIRC (p<1.1e-16)
LAML (p<9.0e-04)
LGG (p<0.0e+00)
LIHC (p<5.9e-06)
LUAD (p<5.1e-03)
LUSC (p<6.8e-03)
OV (p<5.7e-03)
PAAD (p<2.6e-04)
PRAD (p<7.0e-04)
READ (p<1.7e-02)
SARC (p<1.7e-04)
SKCM (p<9.4e-03)
STAD (p<4.8e-03)
THCA (p<1.0e-04)
UCEC (p<1.7e-05)
Survival Probability
Survival Time (Days)
C
Cluster 2
Cluster 3
Master Regulator
STAT3
CEBPD
CEBPB
VIPER Score
D
Best Match, VIPER
112 TCGA Clusters
Cell Lines
VIPER Inferences
TCGA Cluster -
Achilles K.O. Score
E
p < 7.1e-06
density
counts

### Figure S3

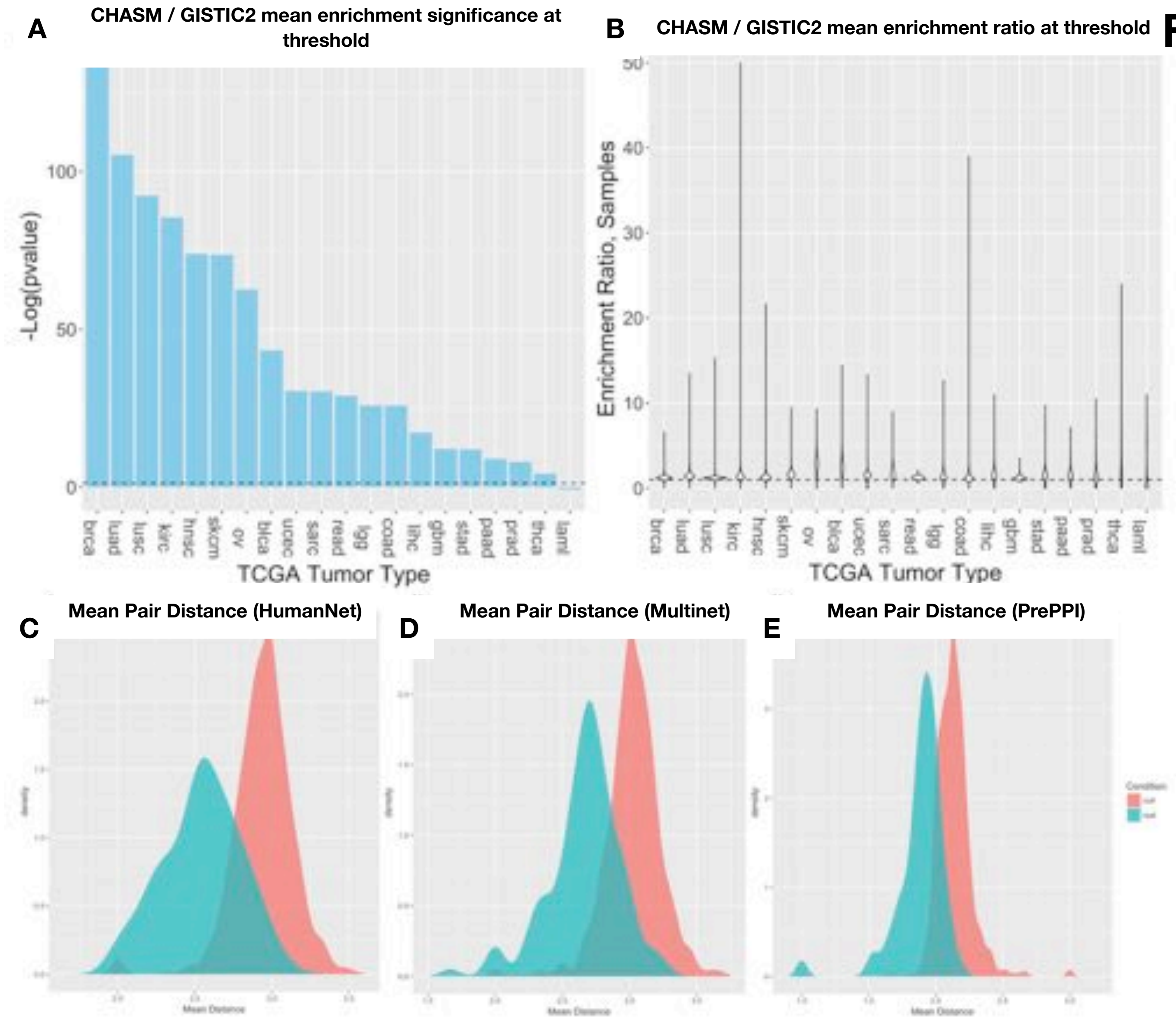
A
CHASM / GISTIC2 mean enrichment significance at threshold
-Log(pvalue)
100
50
0
brca
luad
lusc
kirc
hnsc
skcm
ov
blca
ucec
sarc
read
lgg
coad
lihc
gbm
stad
paad
prad
thca
laml
TCGA Tumor Type
B
CHASM / GISTIC2 mean enrichment ratio at threshold
Enrichment Ratio, Samples
50
40
30
20
10
0
brca
luad
lusc
kirc
hnsc
skcm
ov
blca
ucec
sarc
read
lgg
coad
lihc
gbm
stad
paad
prad
thca
laml
TCGA Tumor Type
C
Mean Pair Distance (HumanNet)
D
Mean Pair Distance (Multinet)
E
Mean Pair Distance (PrePPI)

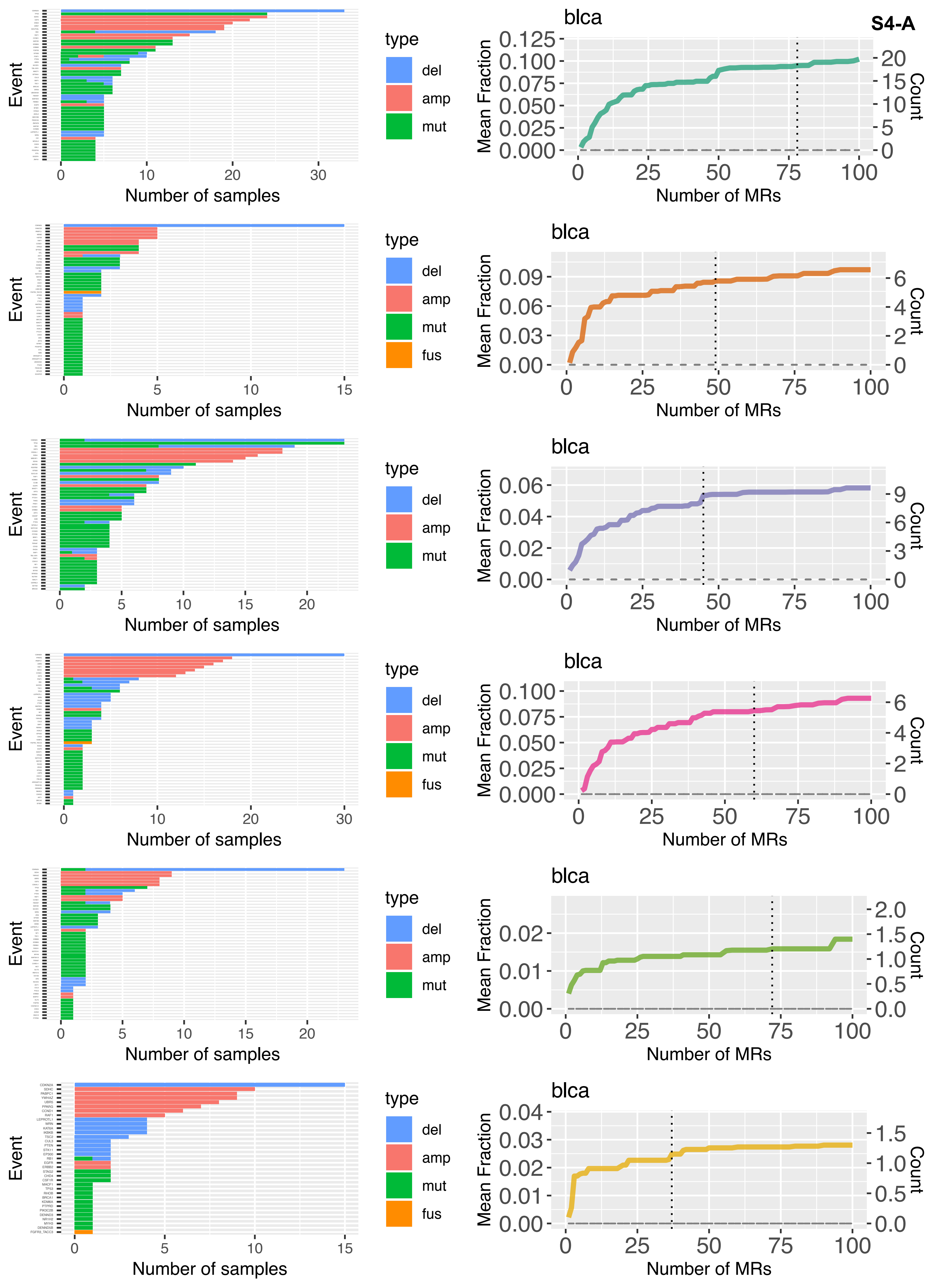
S4-A
Event
Number of samples
type
del
amp
mut
fus
blca
Mean Fraction
Count
Number of MRs
CDKN2A
SDHC
PABPC1
YWHAZ
UBR5
PPARG
CCND1
RAF1
LEPROTL1
WRN
KAT6A
IKBKB
TSC2
CUL3
PTEN
STK11
EP300
RB1
EGFR
ERBB2
STAG2
CHD4
CSF1R
MACF1
TP53
RHOB
BRCA1
KDM6A
PTPRD
PIK3C2B
DENND3
NR1H2
MYH3
DENND5B
FGFR3_TACC3

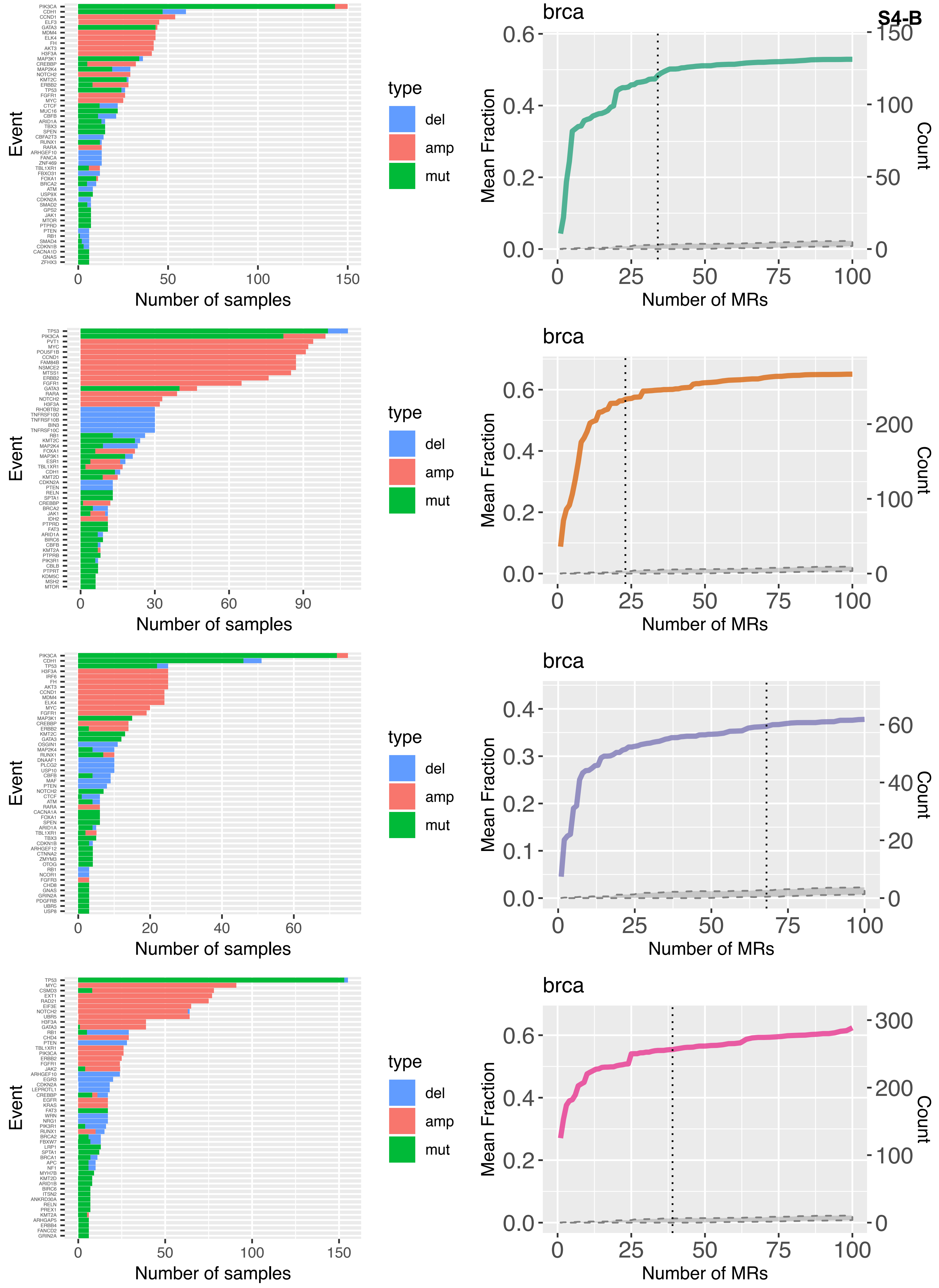
S4-B
brca
Event
Number of samples
type
del
amp
mut
Mean Fraction
Count
Number of MRs

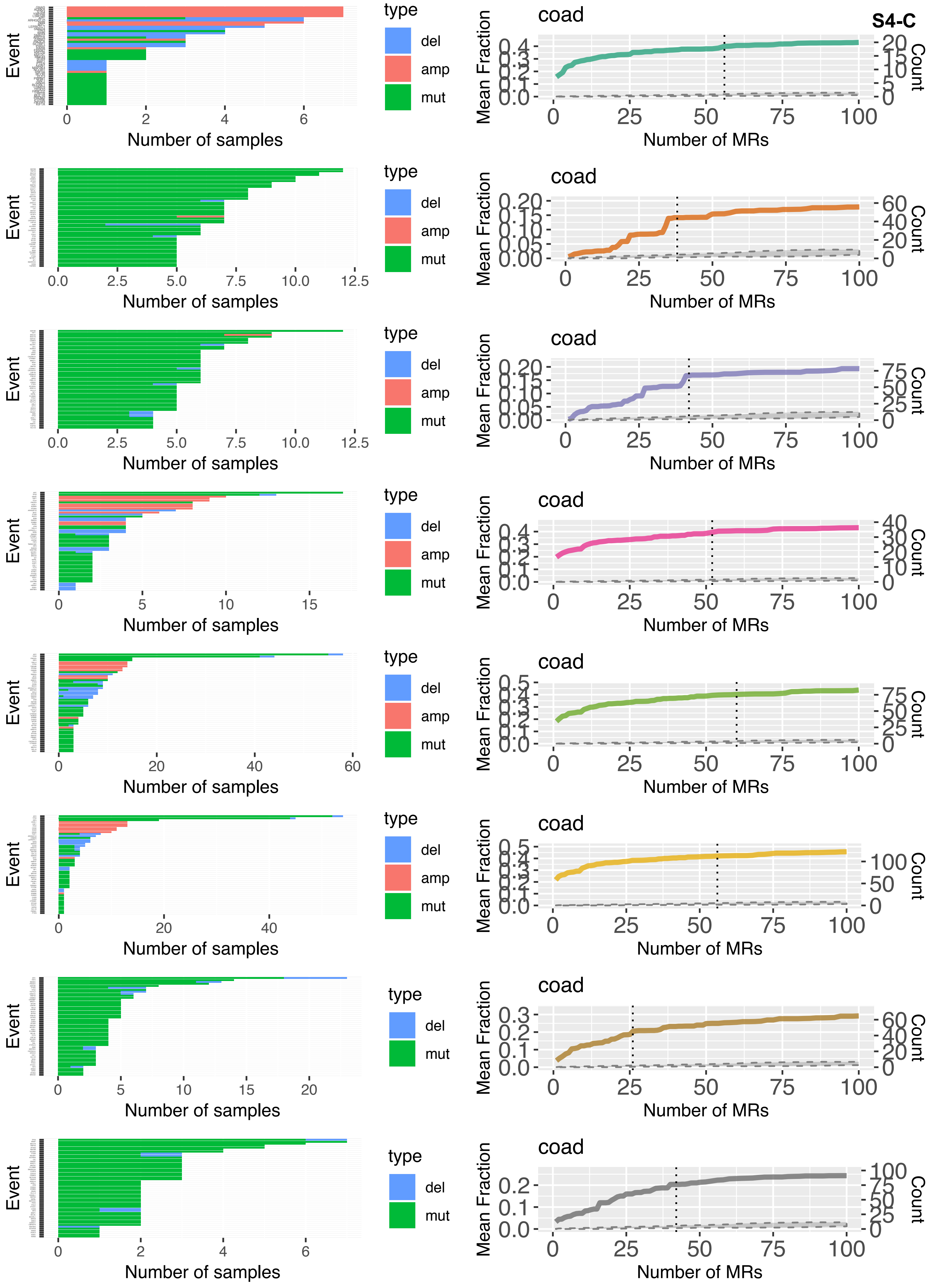
S4-C
Event
Number of samples
type
del
amp
mut
coad
Mean Fraction
Count
Number of MRs
0
2
4
6
0.0
2.5
5.0
7.5
10.0
12.5
0
5
10
15
20
40
60
25
50
75
100
0.0
0.1
0.2
0.3
0.4
0.5
0.00
0.05
0.10
0.15
0.20

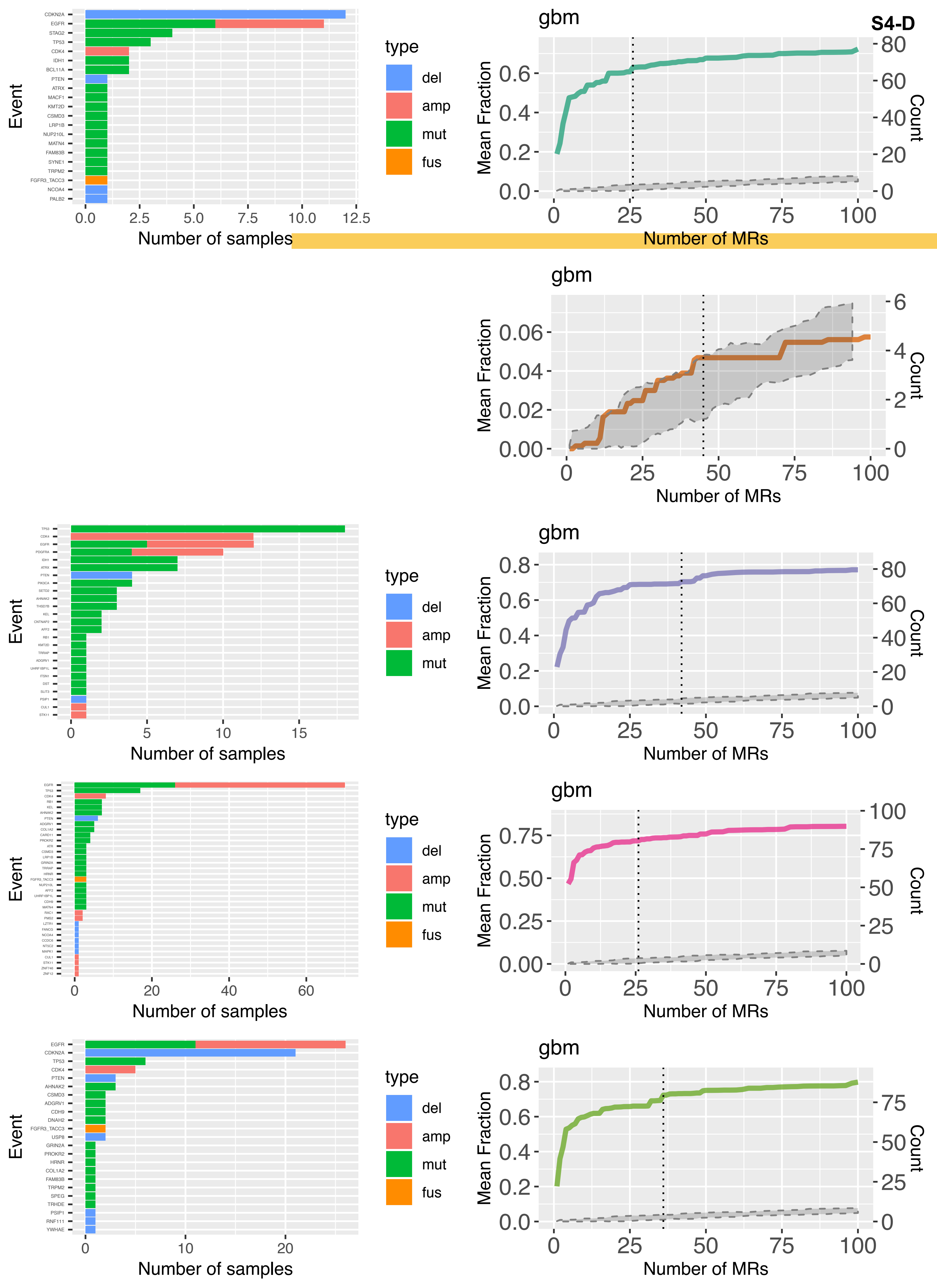

S4-D
gbm
Event
Number of samples
type
del
amp
mut
fus
Mean Fraction
Count
Number of MRs
CDKN2A
EGFR
STAG2
TP53
CDK4
IDH1
BCL11A
PTEN
ATRX
MACF1
KMT2D
CSMD3
LRP1B
NUP210L
MATN4
FAM83B
SYNE1
TRPM2
FGFR3_TACC3
NCOA4
PALB2
EGFR
CDKN2A
TP53
CDK4
PTEN
AHNAK2
CSMD3
ADGRV1
CDH9
DNAH2
FGFR3_TACC3
USP8
GRIN2A
PROKR2
HRNR
COL1A2
FAM83B
TRPM2
SPEG
TRHDE
PSIP1
RNF111
YWHAE

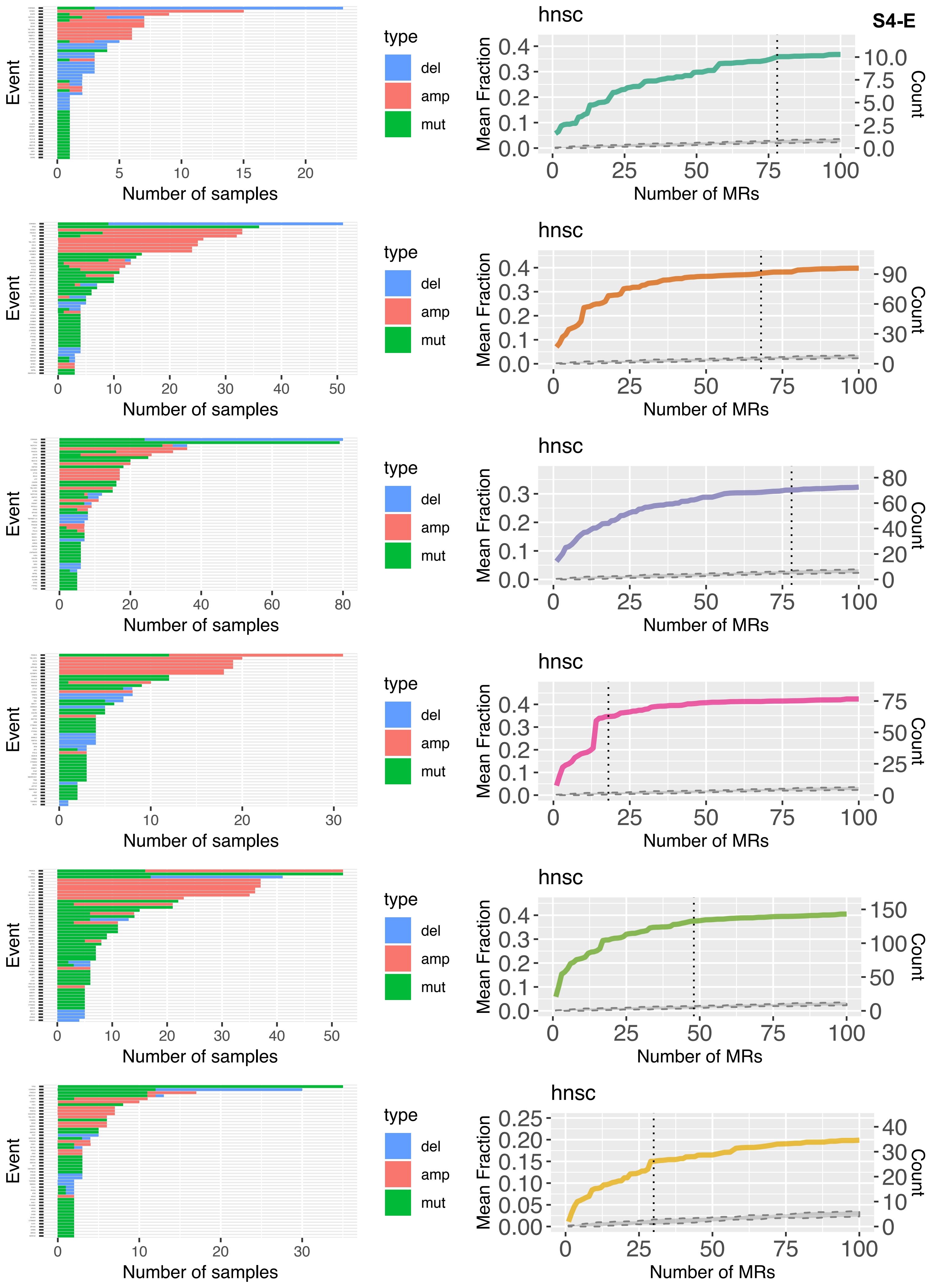

S4-E
hnsc
Event
Number of samples
type
del
amp
mut
Mean Fraction
Count
Number of MRs

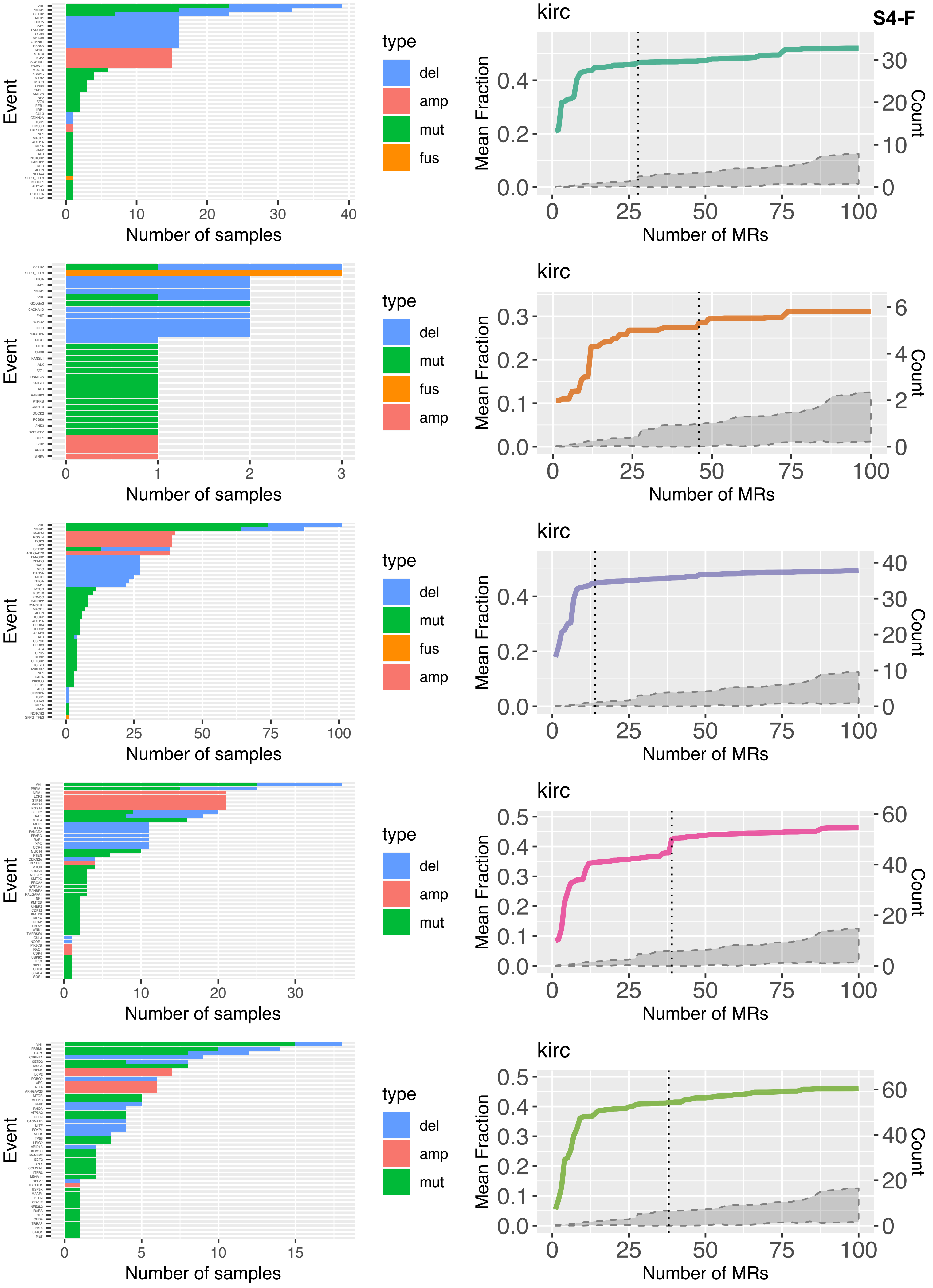
S4-F
Event
Number of samples
type
del
amp
mut
fus
kirc
Mean Fraction
Count
Number of MRs

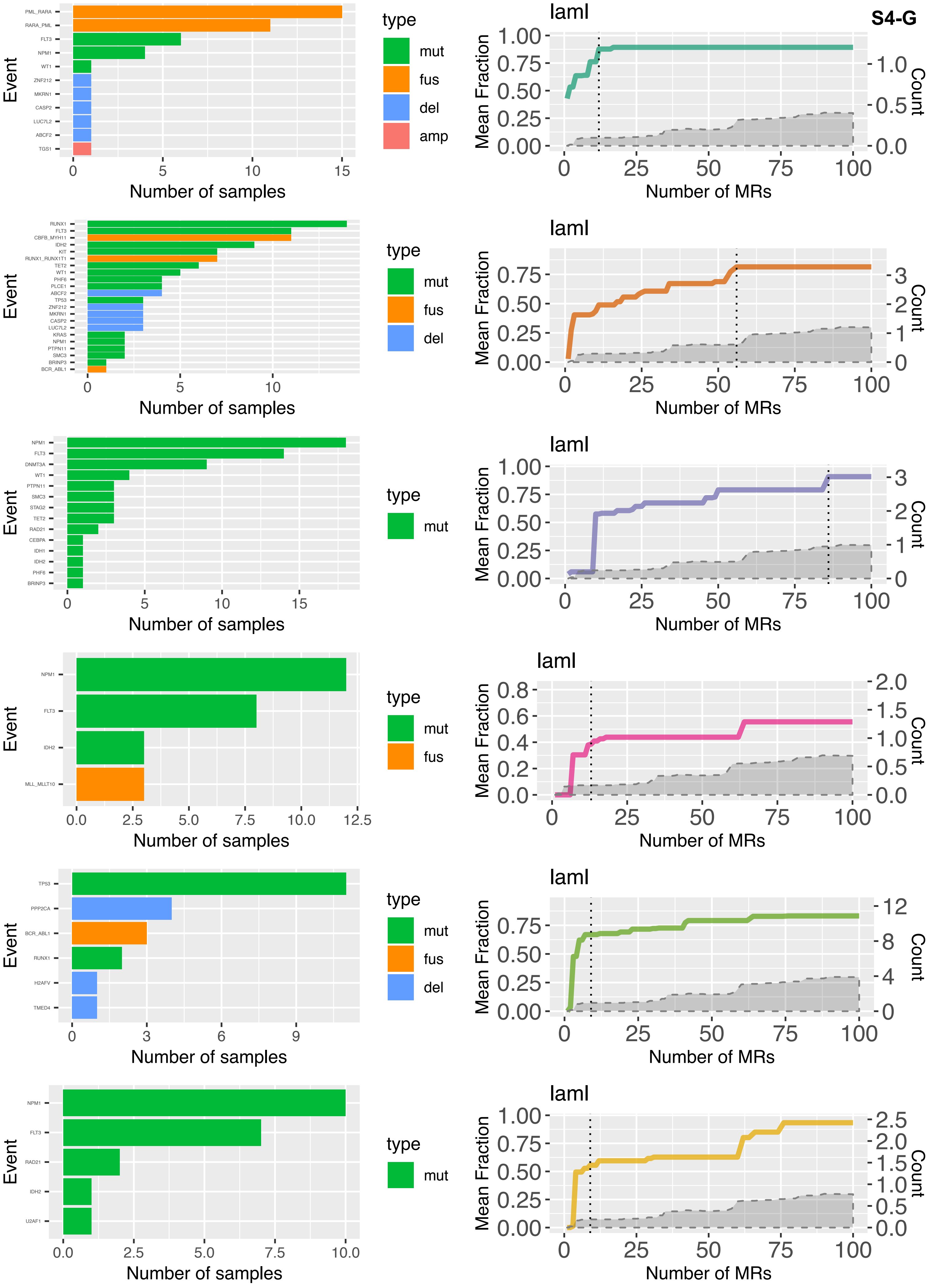
S4-G
laml
Event
Number of samples
type
mut
fus
del
amp
Mean Fraction
Count
Number of MRs
PML_RARA
RARA_PML
FLT3
NPM1
WT1
ZNF212
MKRN1
CASP2
LUC7L2
ABCF2
TGS1
RUNX1
CBFB_MYH11
IDH2
KIT
RUNX1_RUNX1T1
TET2
PHF6
PLCE1
TP53
KRAS
PTPN11
SMC3
BRINP3
BCR_ABL1
DNMT3A
STAG2
RAD21
CEBPA
IDH1
MLL_MLLT10
PPP2CA
H2AFV
TMED4
U2AF1

**S4-H**

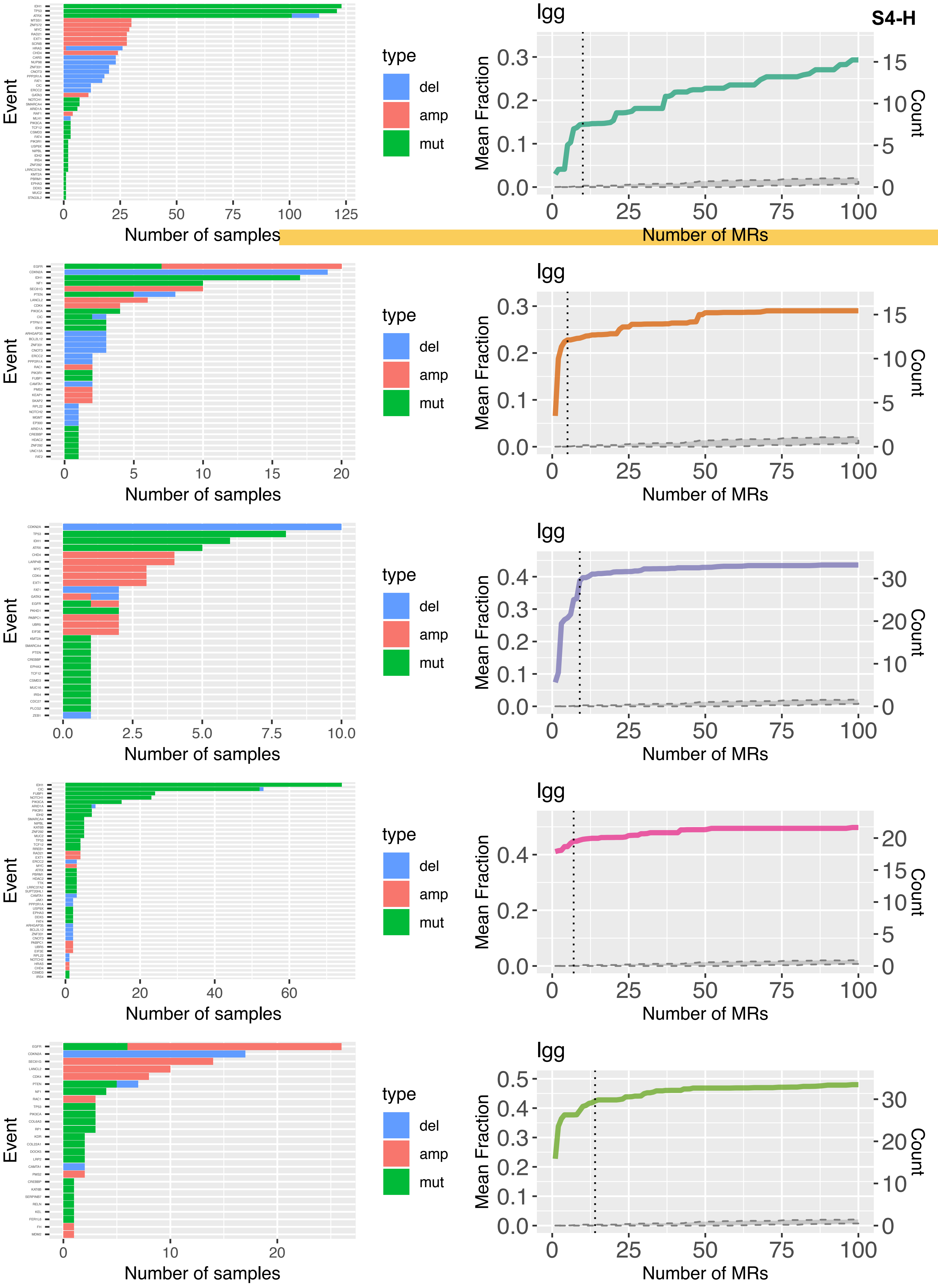
Event
Number of samples
type
del
amp
mut
lgg
Mean Fraction
Count
Number of MRs

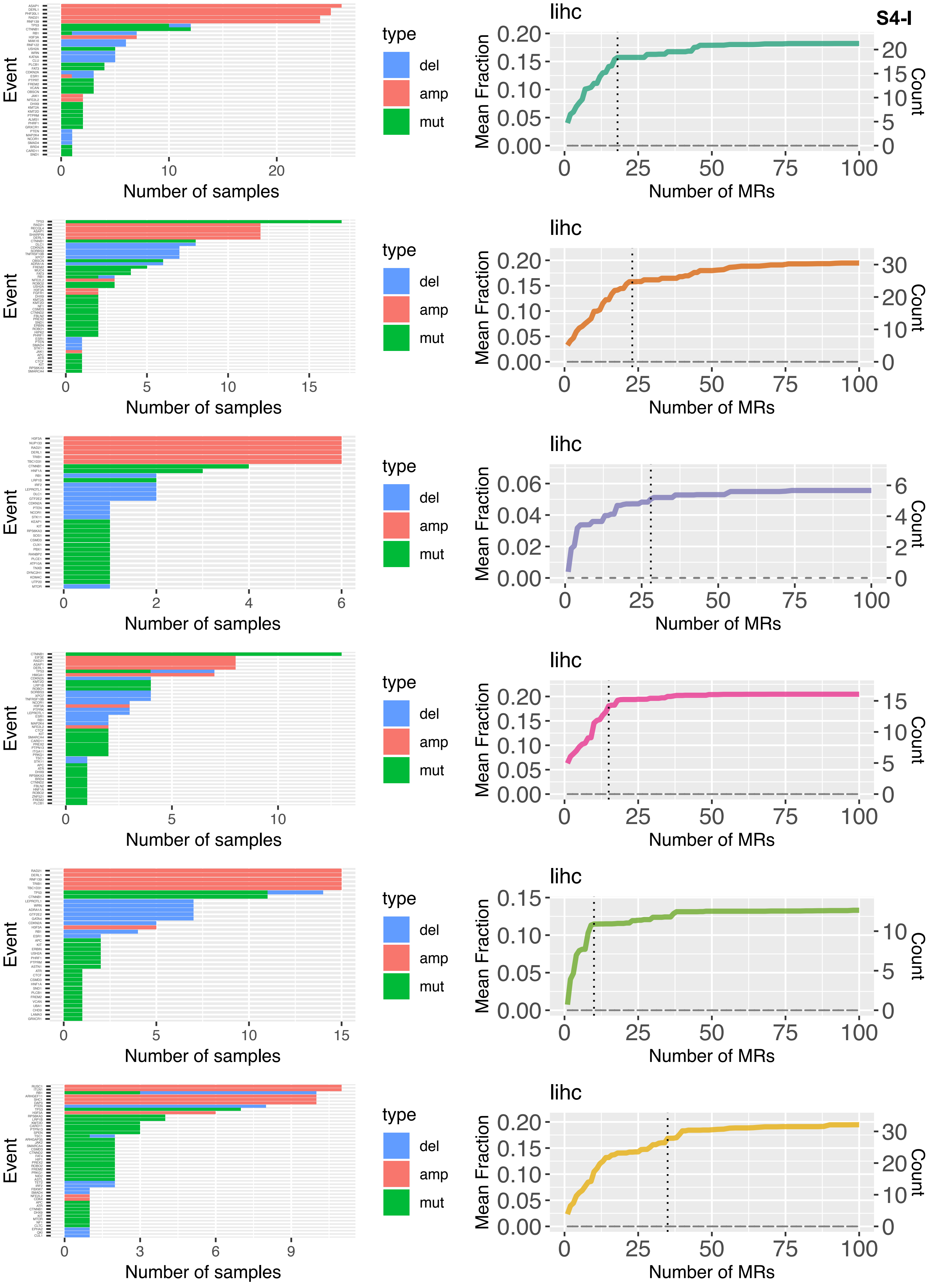
S4-I
Event
Number of samples
type
del
amp
mut
lihc
Mean Fraction
Count
Number of MRs

S4-J

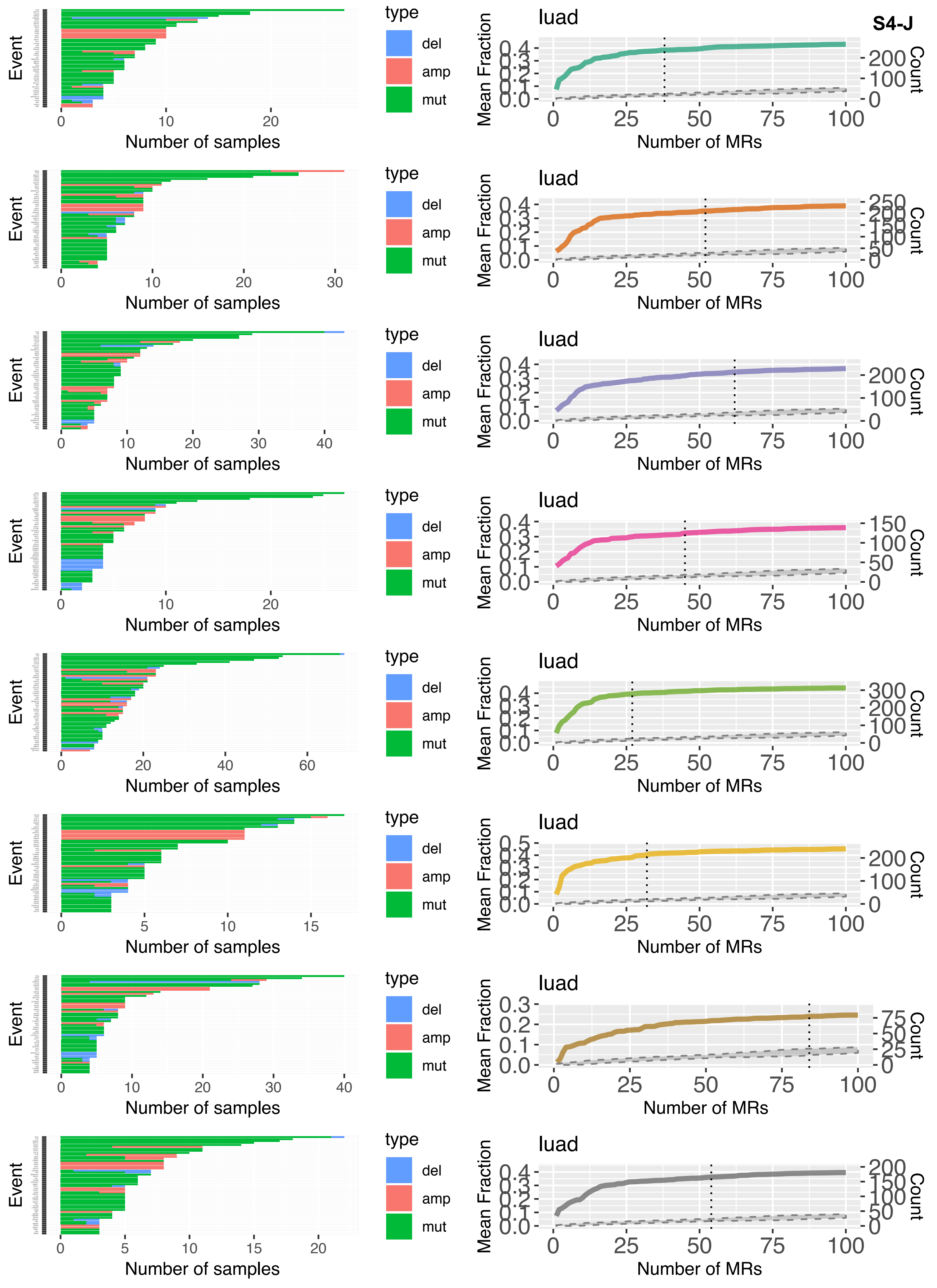
Event
0
10
20
Number of samples
type
del
amp
mut
luad
Mean Fraction
0.0
0.1
0.2
0.3
0.4
Count
0
100
200
0
25
50
75
100
Number of MRs
Event
0
10
20
30
Number of samples
type
del
amp
mut
luad
Mean Fraction
0.0
0.1
0.2
0.3
0.4
Count
0
50
100
150
200
250
0
25
50
75
100
Number of MRs
Event
0
10
20
30
40
Number of samples
type
del
amp
mut
luad
Mean Fraction
0.0
0.1
0.2
0.3
0.4
Count
0
100
200
0
25
50
75
100
Number of MRs
Event
0
10
20
Number of samples
type
del
amp
mut
luad
Mean Fraction
0.0
0.1
0.2
0.3
0.4
Count
0
50
100
150
0
25
50
75
100
Number of MRs
Event
0
20
40
60
Number of samples
type
del
amp
mut
luad
Mean Fraction
0.0
0.1
0.2
0.3
0.4
Count
0
100
200
300
0
25
50
75
100
Number of MRs
Event
0
5
10
15
Number of samples
type
del
amp
mut
luad
Mean Fraction
0.0
0.1
0.2
0.3
0.4
0.5
Count
0
100
200
0
25
50
75
100
Number of MRs
Event
0
10
20
30
40
Number of samples
type
del
amp
mut
luad
Mean Fraction
0.0
0.1
0.2
0.3
Count
0
25
50
75
0
25
50
75
100
Number of MRs
Event
0
5
10
15
20
Number of samples
type
del
amp
mut
luad
Mean Fraction
0.0
0.1
0.2
0.3
0.4
Count
0
50
100
150
200
0
25
50
75
100
Number of MRs

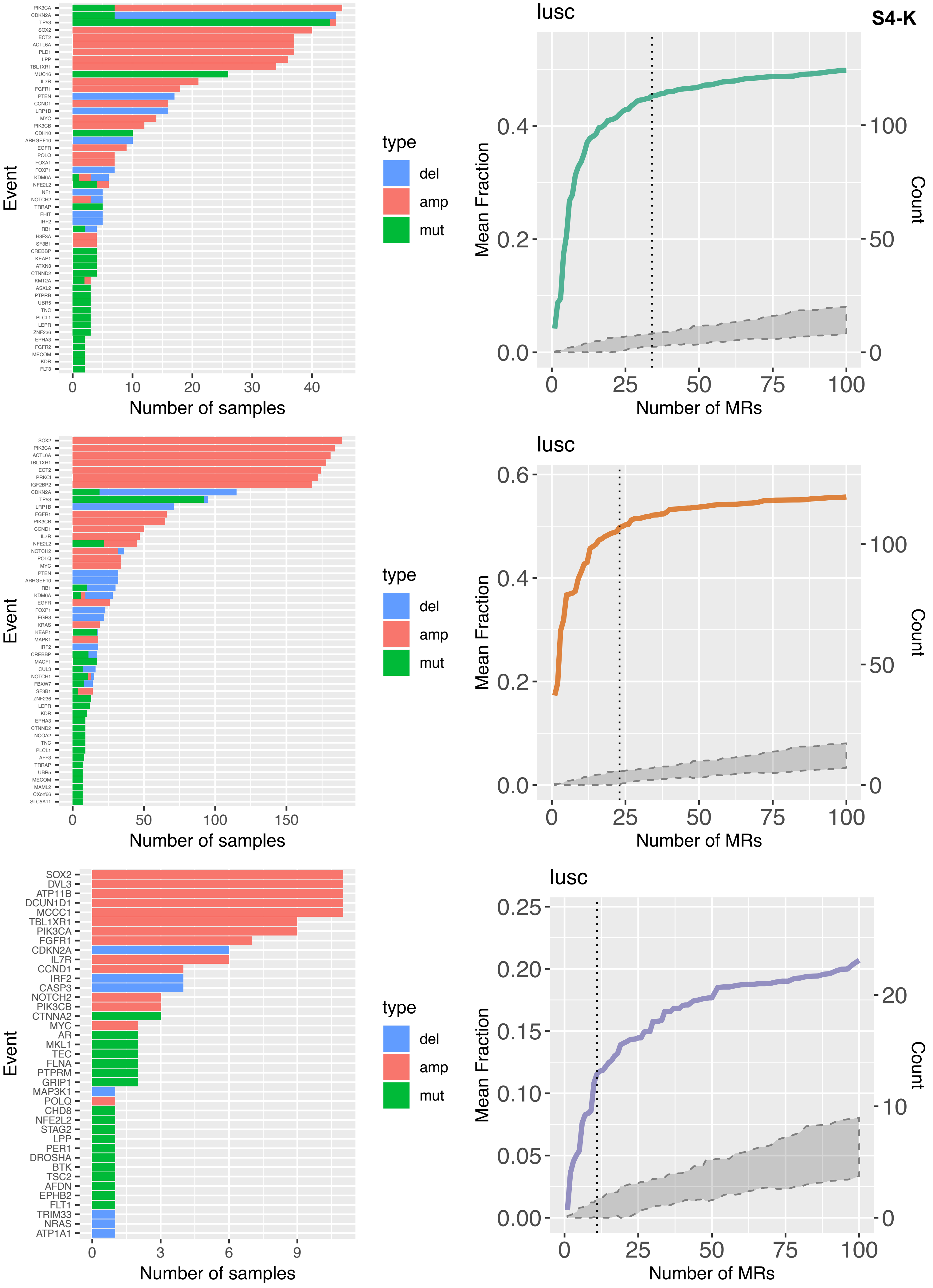
S4-K
lusc
Mean Fraction
Count
Number of MRs
Event
Number of samples
type
del
amp
mut
PIK3CA
CDKN2A
TP53
SOX2
ECT2
ACTL6A
PLD1
LPP
TBL1XR1
MUC16
IL7R
FGFR1
PTEN
CCND1
LRP1B
MYC
PIK3CB
CDH10
ARHGEF10
EGFR
POLQ
FOXA1
FOXP1
KDM6A
NFE2L2
NF1
NOTCH2
TRRAP
FHIT
IRF2
RB1
H3F3A
SF3B1
CREBBP
KEAP1
ATXN3
CTNND2
KMT2A
ASXL2
PTPRB
UBR5
TNC
PLCL1
LEPR
ZNF236
EPHA3
FGFR2
MECOM
KDR
FLT3
SOX2
PIK3CA
ACTL6A
TBL1XR1
ECT2
PRKCI
IGF2BP2
CDKN2A
TP53
LRP1B
FGFR1
PIK3CB
CCND1
IL7R
NFE2L2
NOTCH2
POLQ
MYC
PTEN
ARHGEF10
RB1
KDM6A
EGFR
FOXP1
EGR3
KRAS
KEAP1
MAPK1
IRF2
CREBBP
MACF1
CUL3
NOTCH1
FBXW7
SF3B1
ZNF236
LEPR
KDR
EPHA3
CTNND2
NCOA2
TNC
PLCL1
AFF3
TRRAP
UBR5
MECOM
MAML2
CXorf66
SLC5A11
SOX2
DVL3
ATP11B
DCUN1D1
MCCC1
TBL1XR1
PIK3CA
FGFR1
CDKN2A
IL7R
CCND1
IRF2
CASP3
NOTCH2
PIK3CB
CTNNA2
MYC
AR
MKL1
TEC
FLNA
PTPRM
GRIP1
MAP3K1
POLQ
CHD8
NFE2L2
STAG2
LPP
PER1
DROSHA
BTK
TSC2
AFDN
EPHB2
FLT1
TRIM33
NRAS
ATP1A1

S4-L

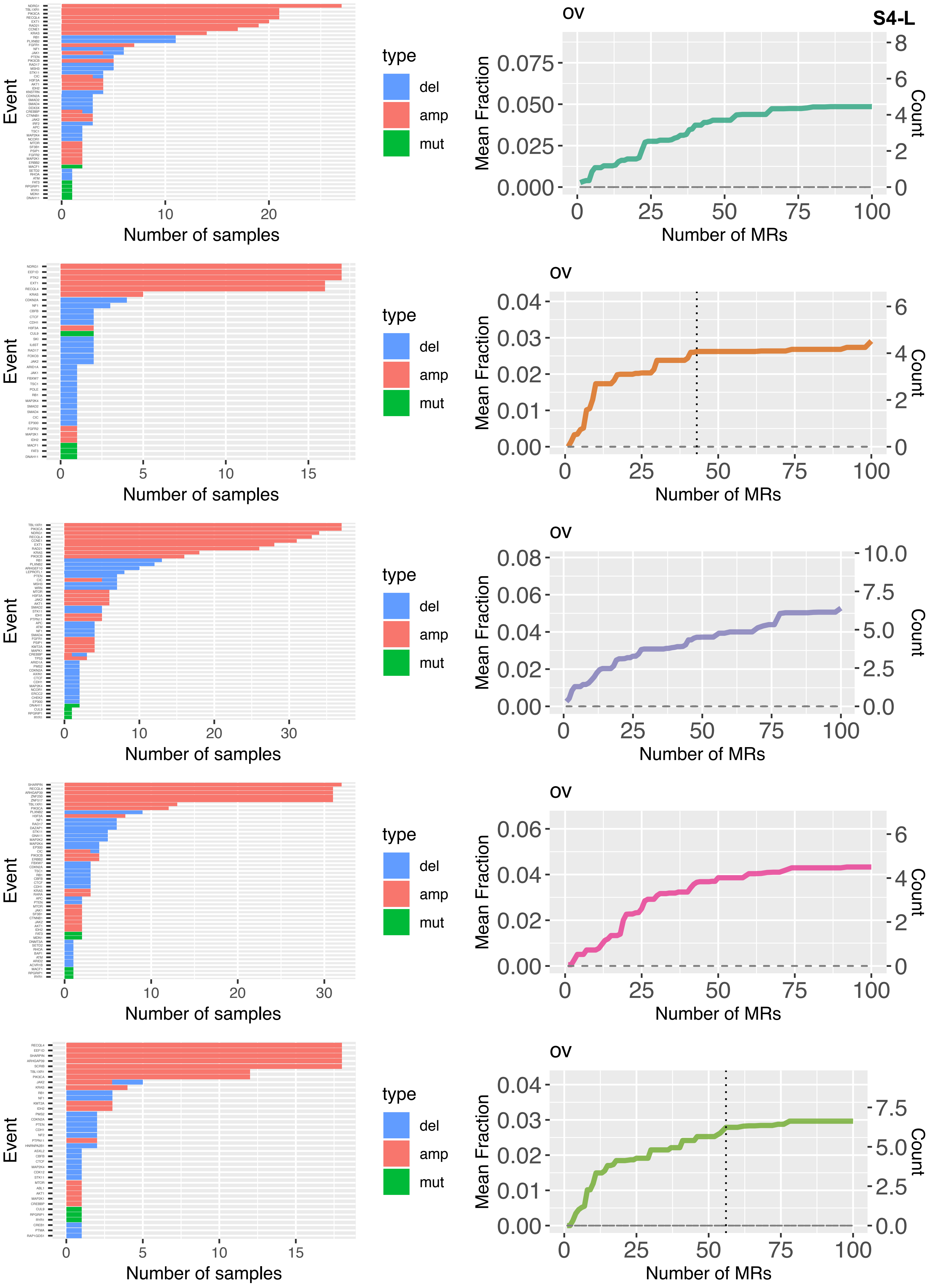
Event
Number of samples
type
del
amp
mut
ov
Mean Fraction
Count
Number of MRs

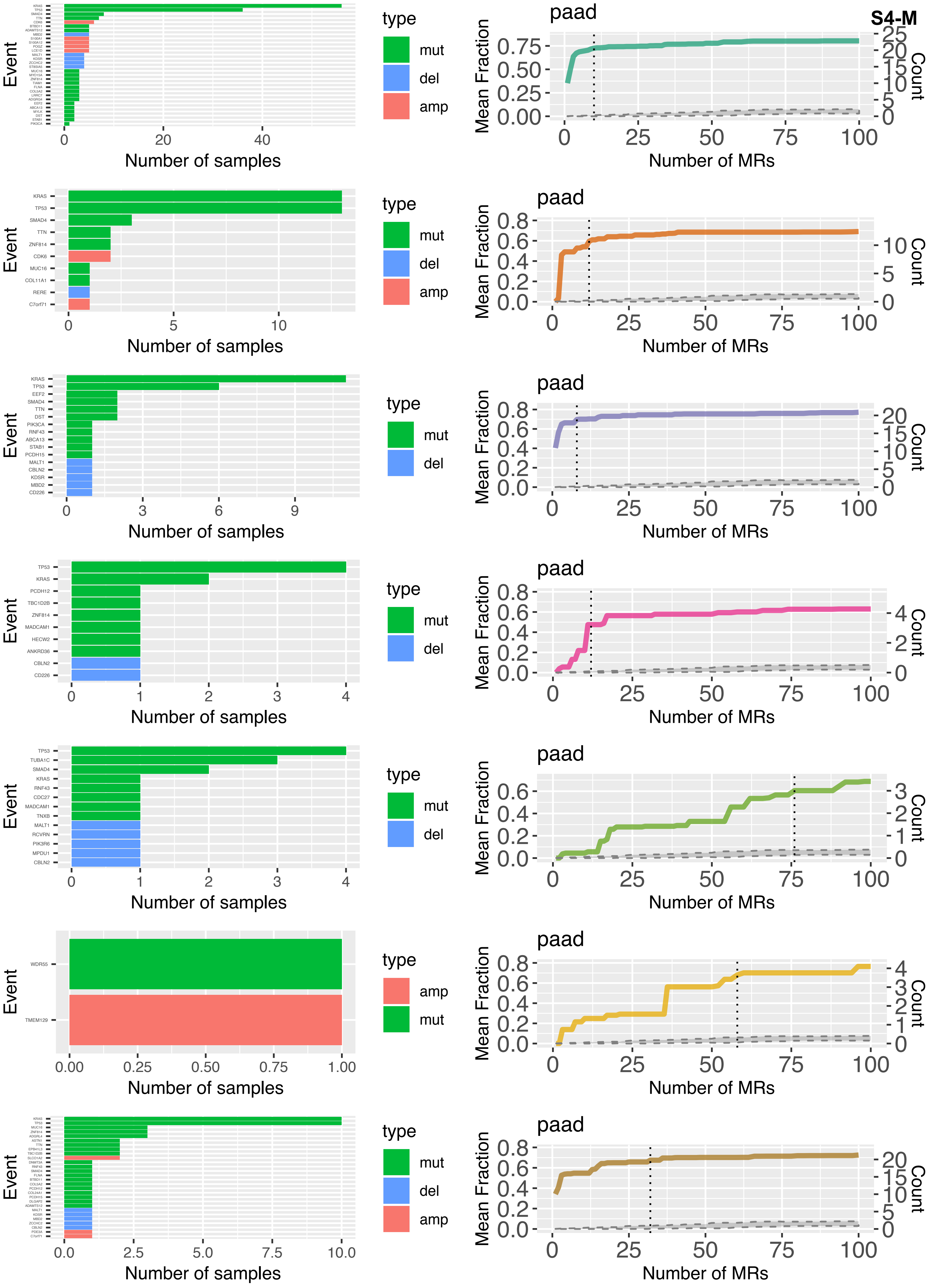
S4-M
paad
Event
Number of samples
type
mut
del
amp
Mean Fraction
Count
Number of MRs
KRAS
TP53
SMAD4
TTN
ZNF814
CDK6
MUC16
COL11A1
RERE
C7orf71
EEF2
DST
PIK3CA
RNF43
ABCA13
STAB1
PCDH15
MALT1
CBLN2
KDSR
MBD2
CD226
PCDH12
TBC1D2B
MADCAM1
HECW2
ANKRD36
TUBA1C
CDC27
TNXB
RCVRN
PIK3R6
MPDU1
WDR55
TMEM129

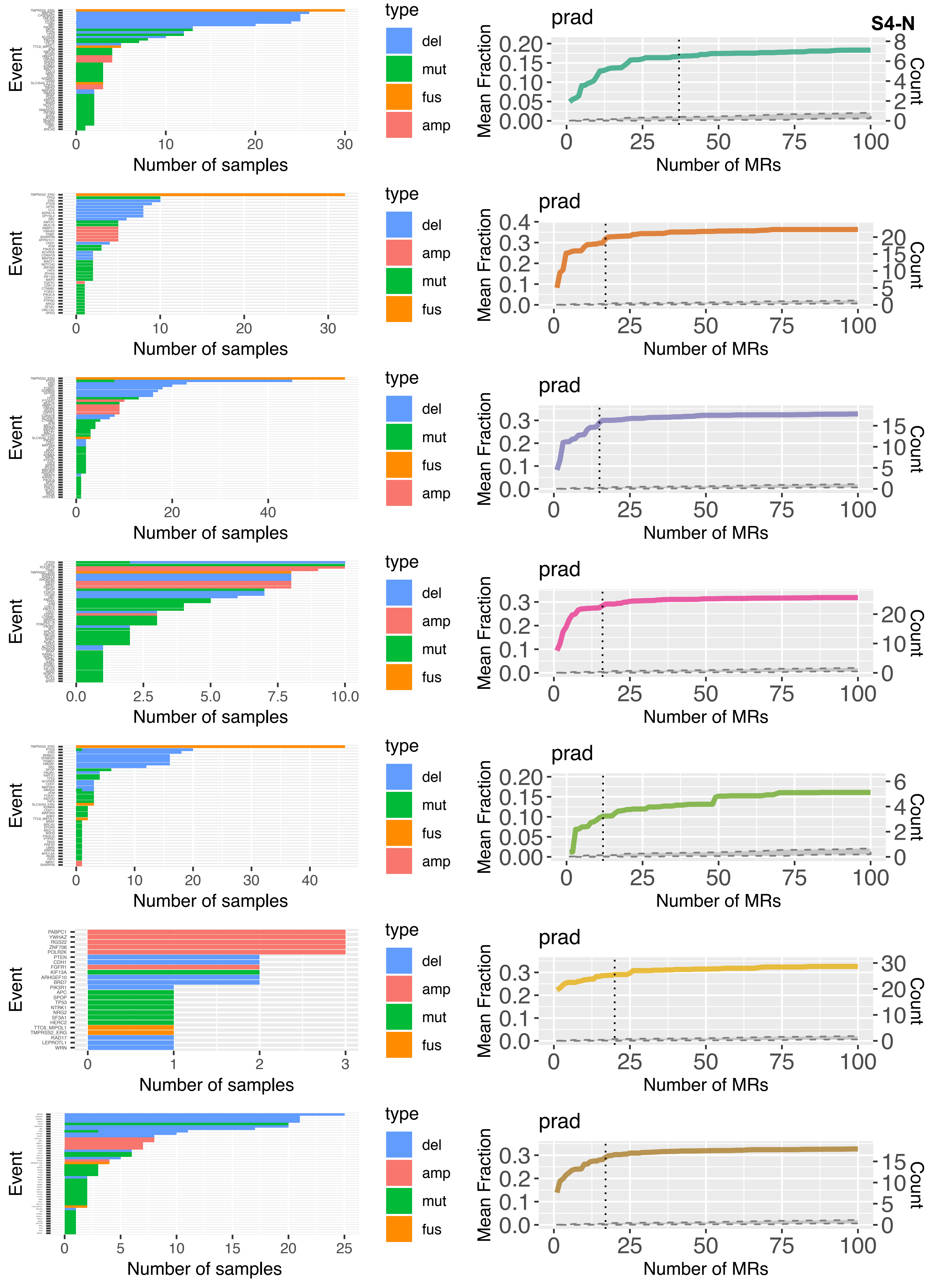
S4-N
Event
Number of samples
type
del
mut
fus
amp
prad
Mean Fraction
Count
Number of MRs
PABPC1
YWHAZ
RGS22
ZNF706
POLR2K
PTEN
CDH1
FGFR1
KIF13A
ARHGEF10
BRD7
PIK3R1
APC
SPOP
TP53
NTRK1
NRG2
SF3A1
HERC2
TTC6_MIPOL1
TMPRSS2_ERG
RAD17
LEPROTL1
WRN

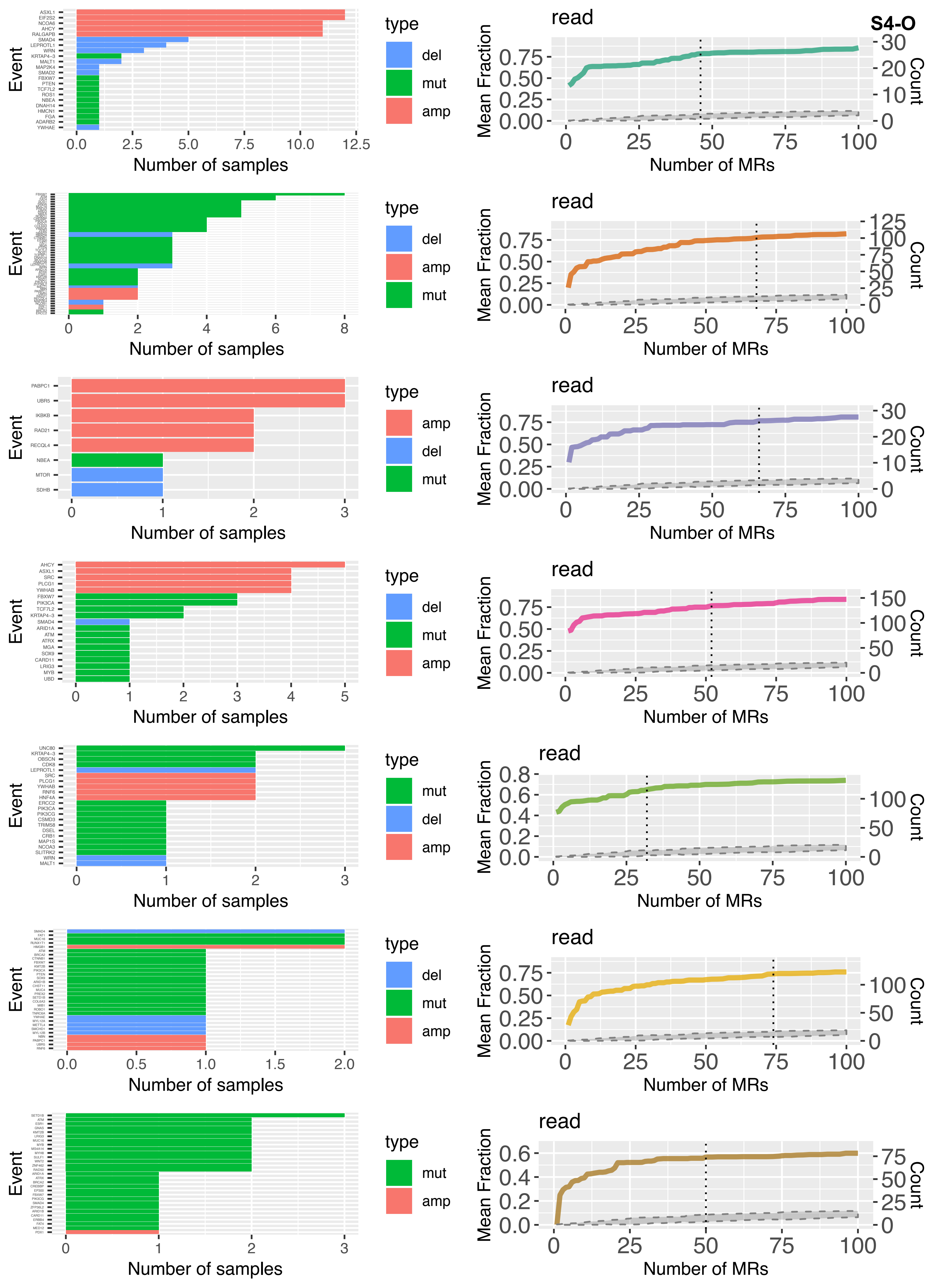
S4-O
Event
ASXL1
EIF2S2
NCOA6
AHCY
RALGAPB
SMAD4
LEPROTL1
WRN
KRTAP4−3
MALT1
MAP2K4
SMAD2
FBXW7
PTEN
TCF7L2
ROS1
NBEA
DNAH14
HMCN1
FGA
ADARB2
YWHAE
0.0 2.5 5.0 7.5 10.0 12.5
Number of samples
type
del
mut
amp
read
Mean Fraction
0.00 0.25 0.50 0.75
Count
0 10 20 30
0 25 50 75 100
Number of MRs
Event
0 2 4 6 8
Number of samples
type
del
amp
mut
read
Mean Fraction
0.00 0.25 0.50 0.75
Count
0 25 50 75 100 125
0 25 50 75 100
Number of MRs
Event
PABPC1
UBR5
IKBKB
RAD21
RECQL4
NBEA
MTOR
SDHB
0 1 2 3
Number of samples
type
amp
del
mut
read
Mean Fraction
0.00 0.25 0.50 0.75
Count
0 10 20 30
0 25 50 75 100
Number of MRs
Event
AHCY
ASXL1
SRC
PLCG1
YWHAB
FBXW7
PIK3CA
TCF7L2
KRTAP4−3
SMAD4
ARID1A
ATM
ATRX
MGA
SOX9
CARD11
LRIG3
MYB
UBD
0 1 2 3 4 5
Number of samples
type
del
mut
amp
read
Mean Fraction
0.00 0.25 0.50 0.75
Count
0 50 100 150
0 25 50 75 100
Number of MRs
Event
UNC80
KRTAP4−3
OBSCN
CDK8
LEPROTL1
SRC
PLCG1
YWHAB
RNF6
HNF4A
ERCC2
PIK3CA
PIK3CG
CSMD3
TRIM58
DSEL
CRB1
MAP1S
NCOA3
SLITRK2
WRN
MALT1
0 1 2 3
Number of samples
type
mut
del
amp
read
Mean Fraction
0.0 0.2 0.4 0.6 0.8
Count
0 50 100
0 25 50 75 100
Number of MRs
Event
0.0 0.5 1.0 1.5 2.0
Number of samples
type
del
mut
amp
read
Mean Fraction
0.00 0.25 0.50 0.75
Count
0 50 100
0 25 50 75 100
Number of MRs
Event
0 1 2 3
Number of samples
type
mut
amp
read
Mean Fraction
0.0 0.2 0.4 0.6
Count
0 25 50 75
0 25 50 75 100
Number of MRs

S4-P

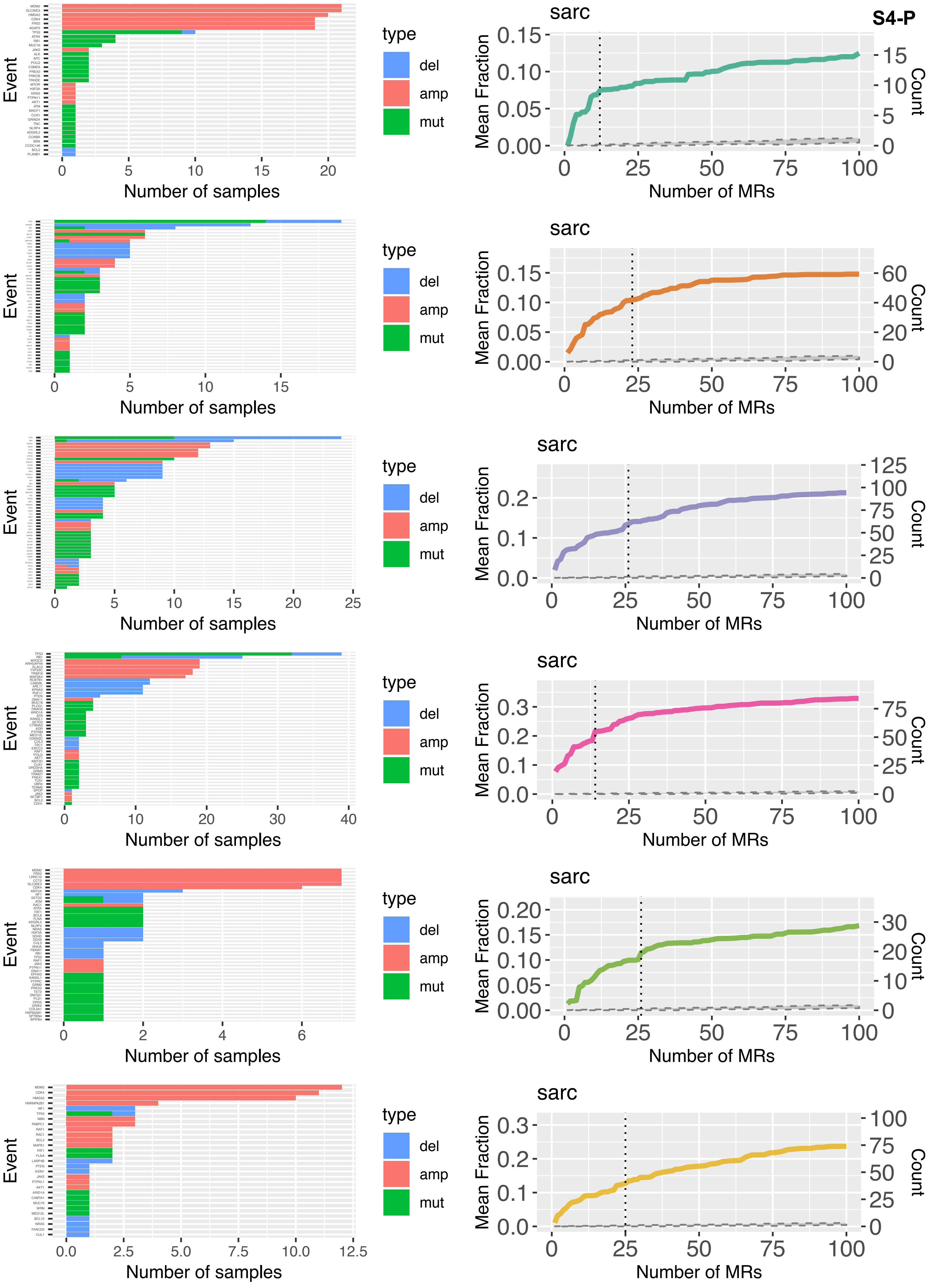
Event
Number of samples
type
del
amp
mut
sarc
Mean Fraction
Count
Number of MRs

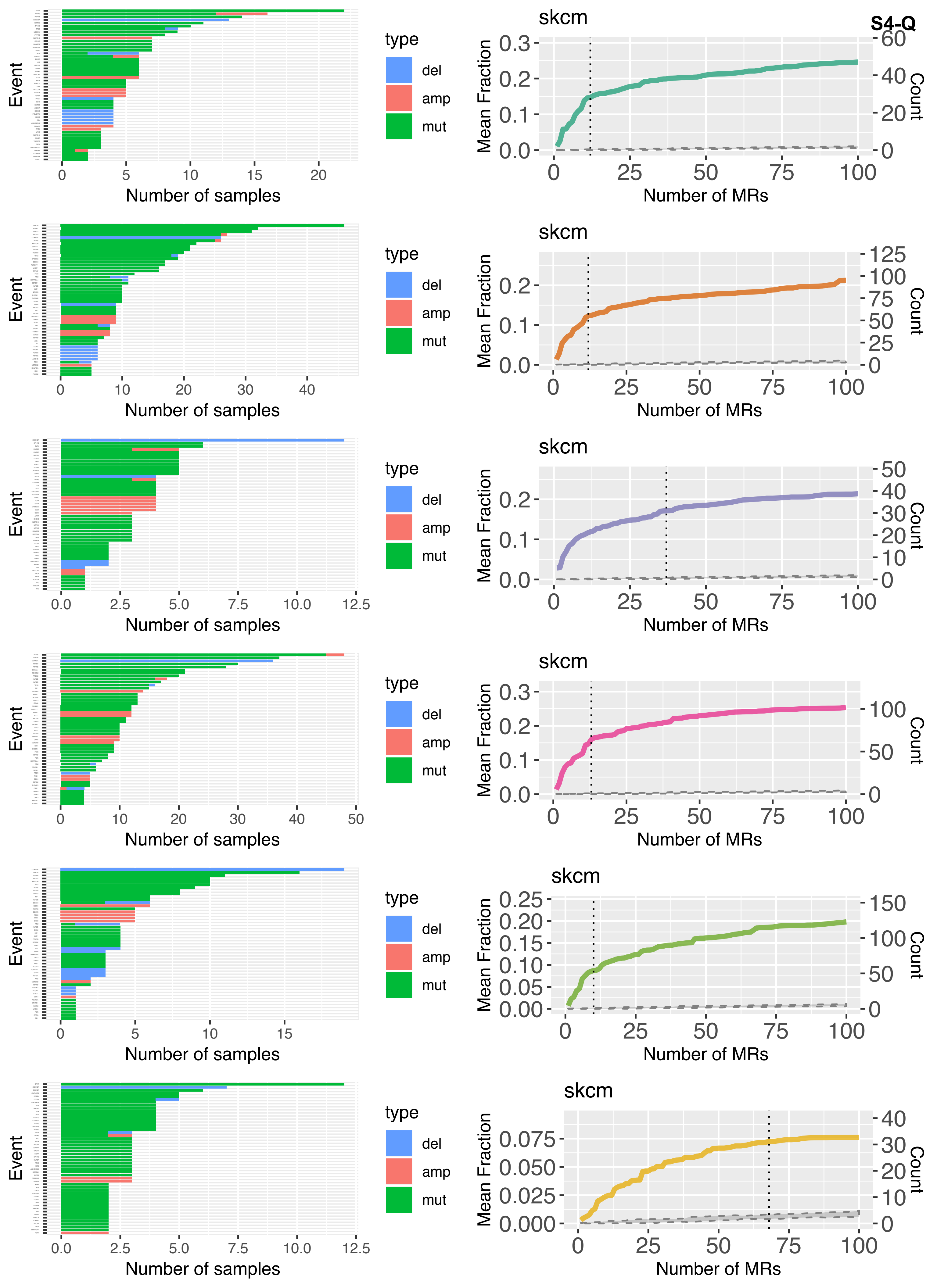
S4-Q
skcm
Event
Number of samples
type
del
amp
mut
Mean Fraction
Count
Number of MRs

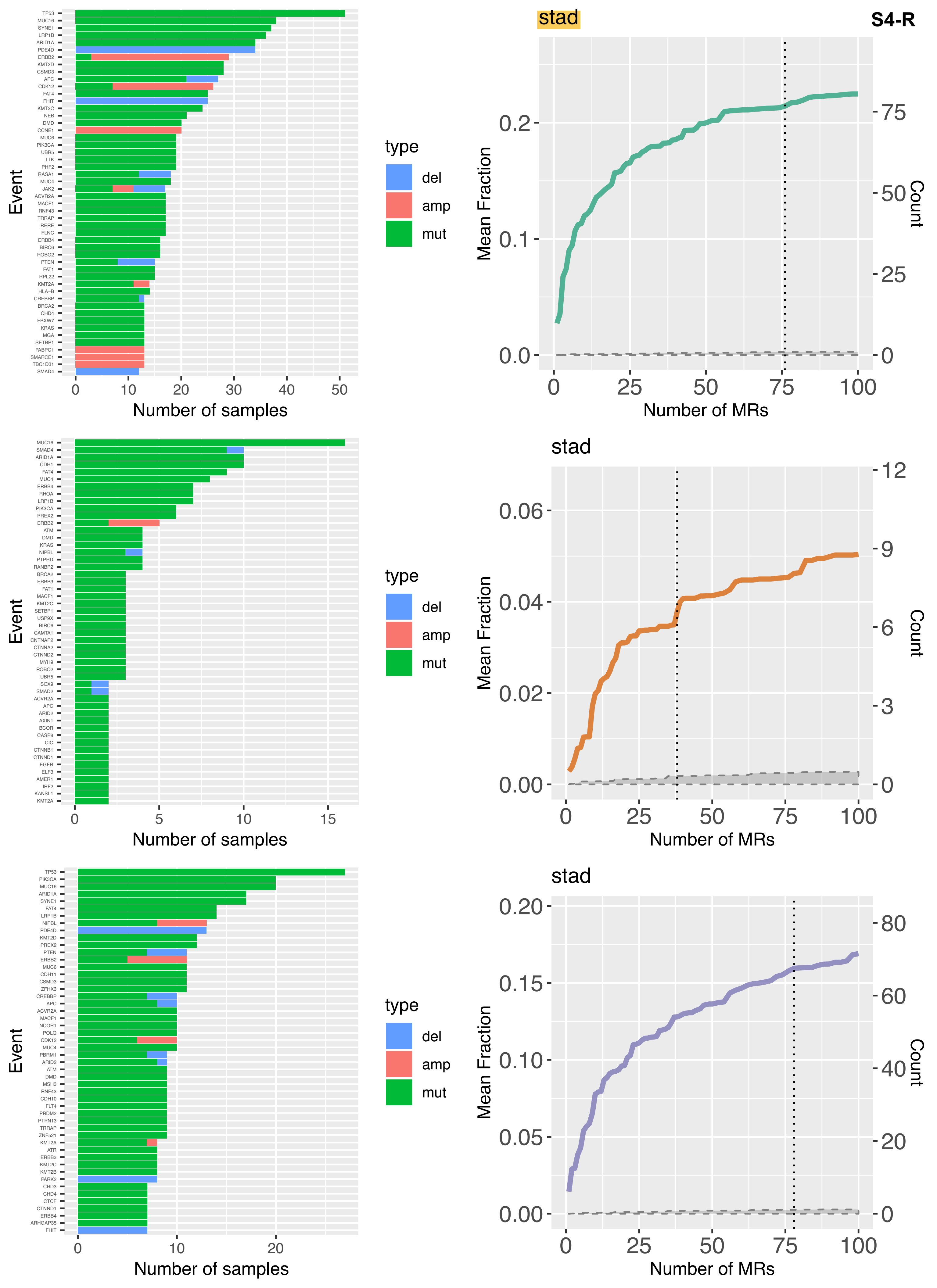
S4-R
stad
Event
Number of samples
type
del
amp
mut
Mean Fraction
Count
Number of MRs
stad
Event
Number of samples
type
del
amp
mut
Mean Fraction
Count
Number of MRs
stad
Event
Number of samples
type
del
amp
mut
Mean Fraction
Count
Number of MRs

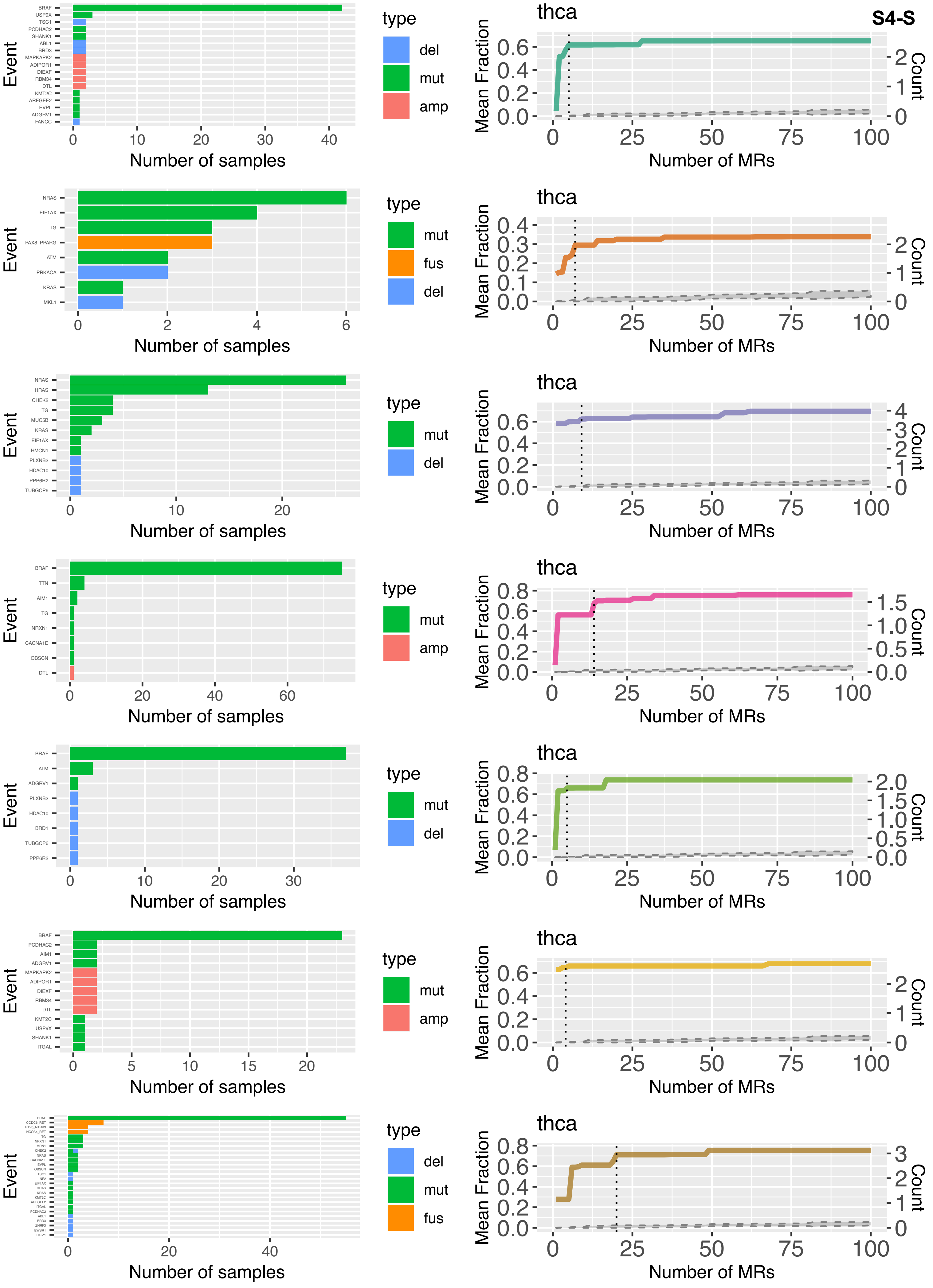
S4-S
Event
BRAF
USP9X
TSC1
PCDHAC2
SHANK1
ABL1
BRD3
MAPKAPK2
ADIPOR1
DIEXF
RBM34
DTL
KMT2C
ARFGEF2
EVPL
ADGRV1
FANCC
Number of samples
type
del
mut
amp
thca
Mean Fraction
Count
Number of MRs
NRAS
EIF1AX
TG
PAX8_PPARG
ATM
PRKACA
KRAS
MKL1
fus
HRAS
CHEK2
MUC5B
HMCN1
PLXNB2
HDAC10
PPP6R2
TUBGCP6
TTN
AIM1
NRXN1
CACNA1E
OBSCN
BRD1
ITGAL

**S4-T**

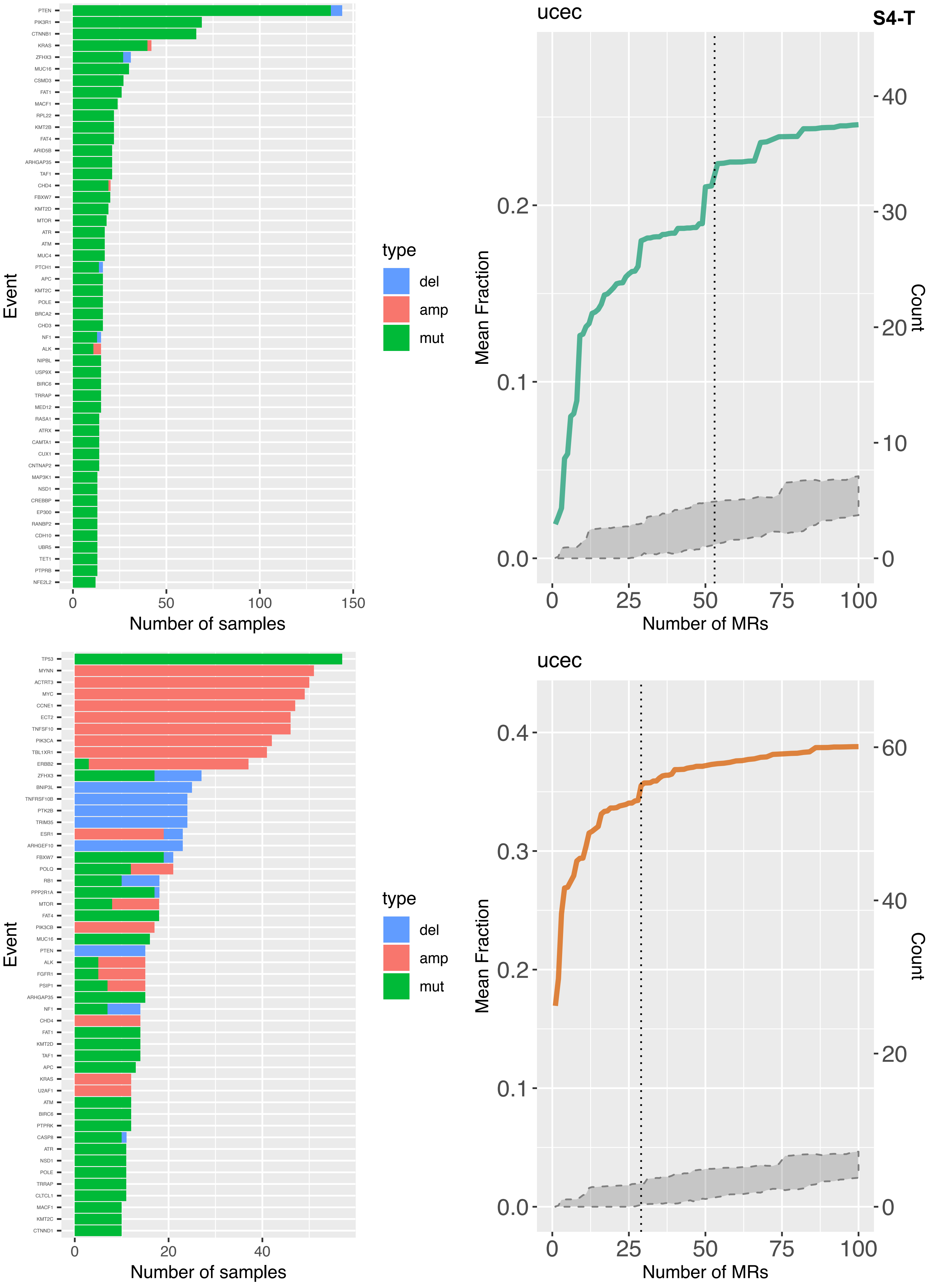

ucec
PTEN
PIK3R1
CTNNB1
KRAS
ZFHX3
MUC16
CSMD3
FAT1
MACF1
RPL22
KMT2B
FAT4
ARID5B
ARHGAP35
TAF1
CHD4
FBXW7
KMT2D
MTOR
ATR
ATM
MUC4
PTCH1
APC
KMT2C
POLE
BRCA2
CHD3
NF1
ALK
NIPBL
USP9X
BIRC6
TRRAP
MED12
RASA1
ATRX
CAMTA1
CUX1
CNTNAP2
MAP3K1
NSD1
CREBBP
EP300
RANBP2
CDH10
UBR5
TET1
PTPRB
NFE2L2
Event
Number of samples
type
del
amp
mut
Mean Fraction
Count
Number of MRs
ucec
TP53
MYNN
ACTRT3
MYC
CCNE1
ECT2
TNFSF10
PIK3CA
TBL1XR1
ERBB2
ZFHX3
BNIP3L
TNFRSF10B
PTK2B
TRIM35
ESR1
ARHGEF10
FBXW7
POLQ
RB1
PPP2R1A
MTOR
FAT4
PIK3CB
MUC16
PTEN
ALK
FGFR1
PSIP1
ARHGAP35
NF1
CHD4
FAT1
KMT2D
TAF1
APC
KRAS
U2AF1
ATM
BIRC6
PTPRK
CASP8
ATR
NSD1
POLE
TRRAP
CLTCL1
MACF1
KMT2C
CTNND1
Event
Number of samples
type
del
amp
mut
Mean Fraction
Count
Number of MRs

### Figure S5

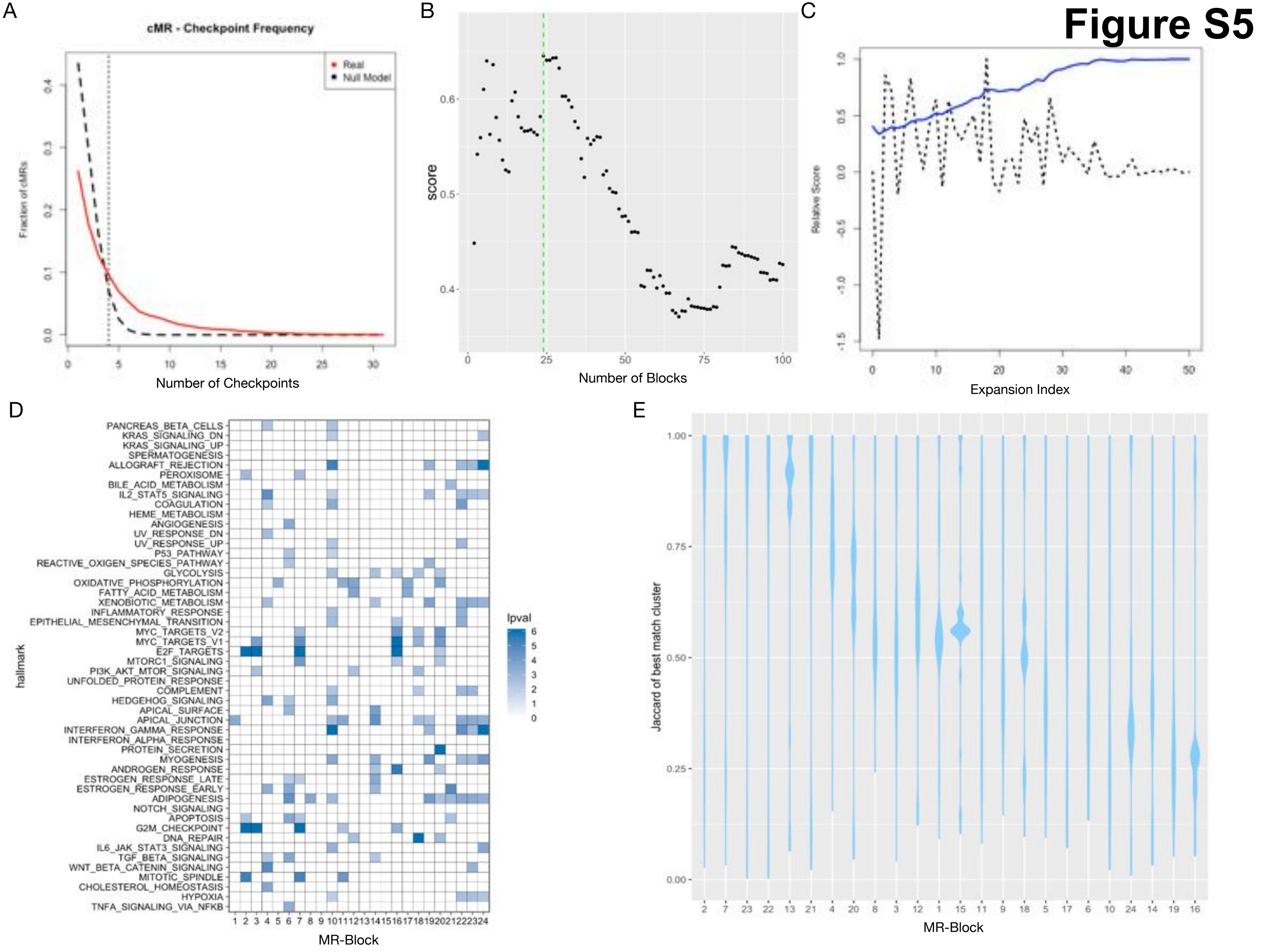

A
cMR - Checkpoint Frequency
Real
Null Model
Fraction of cMRs
Number of Checkpoints
B
score
Number of Blocks
C
Relative Score
Expansion Index
D
hallmark
PANCREAS_BETA_CELLS
KRAS_SIGNALING_DN
KRAS_SIGNALING_UP
SPERMATOGENESIS
ALLOGRAFT_REJECTION
PEROXISOME
BILE_ACID_METABOLISM
IL2_STAT5_SIGNALING
COAGULATION
HEME_METABOLISM
ANGIOGENESIS
UV_RESPONSE_DN
UV_RESPONSE_UP
P53_PATHWAY
REACTIVE_OXIGEN_SPECIES_PATHWAY
GLYCOLYSIS
OXIDATIVE_PHOSPHORYLATION
FATTY_ACID_METABOLISM
XENOBIOTIC_METABOLISM
INFLAMMATORY_RESPONSE
EPITHELIAL_MESENCHYMAL_TRANSITION
MYC_TARGETS_V2
MYC_TARGETS_V1
E2F_TARGETS
MTORC1_SIGNALING
PI3K_AKT_MTOR_SIGNALING
UNFOLDED_PROTEIN_RESPONSE
COMPLEMENT
HEDGEHOG_SIGNALING
APICAL_SURFACE
APICAL_JUNCTION
INTERFERON_GAMMA_RESPONSE
INTERFERON_ALPHA_RESPONSE
PROTEIN_SECRETION
MYOGENESIS
ANDROGEN_RESPONSE
ESTROGEN_RESPONSE_LATE
ESTROGEN_RESPONSE_EARLY
ADIPOGENESIS
NOTCH_SIGNALING
APOPTOSIS
G2M_CHECKPOINT
DNA_REPAIR
IL6_JAK_STAT3_SIGNALING
TGF_BETA_SIGNALING
WNT_BETA_CATENIN_SIGNALING
MITOTIC_SPINDLE
CHOLESTEROL_HOMEOSTASIS
HYPOXIA
TNFA_SIGNALING_VIA_NFKB
lpval
MR-Block
E
Jaccard of best match cluster
MR-Block

### Figure S6

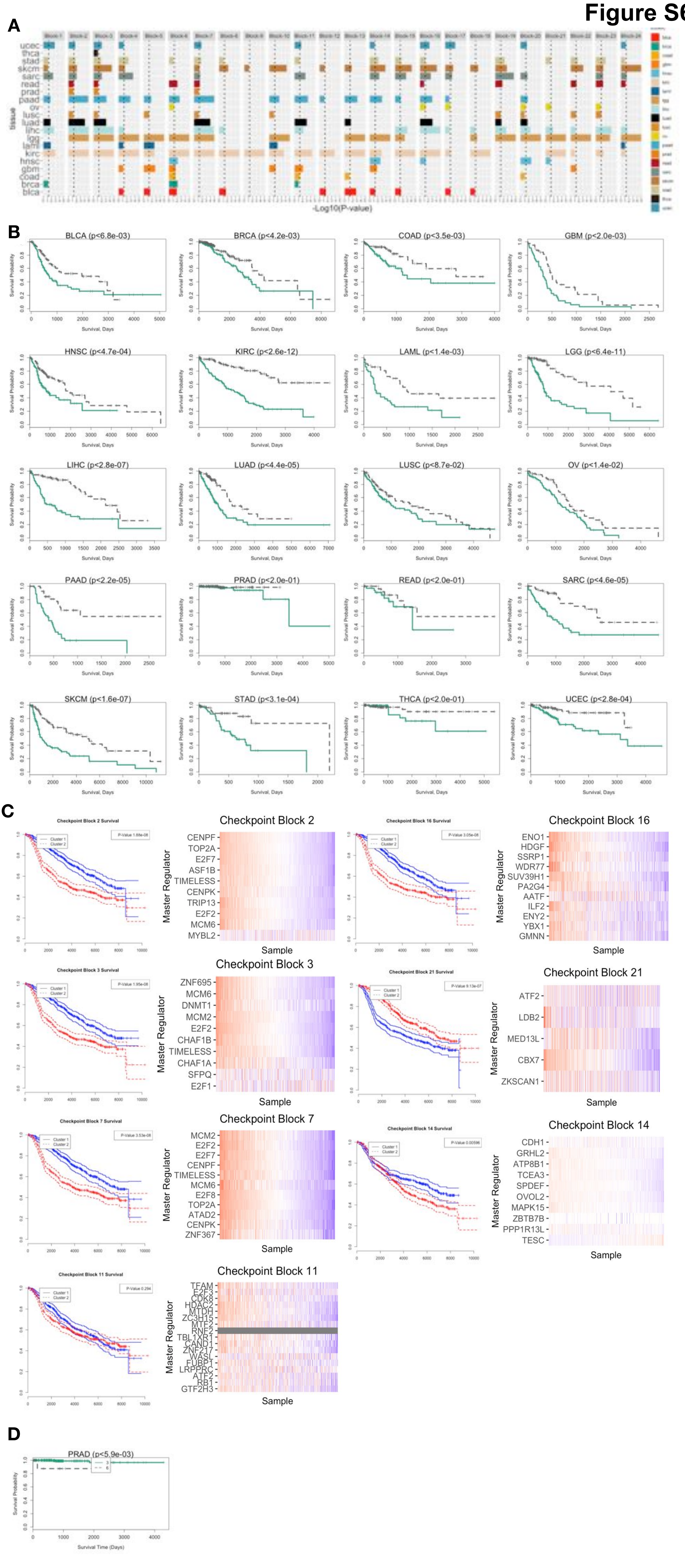


1,3,5
6
